# Diversity, host ranges, and potential drivers of speciation among the inquiline enemies of oak gall wasps

**DOI:** 10.1101/2020.05.08.084699

**Authors:** Anna KG Ward, Sofia I Sheikh, Andrew A Forbes

## Abstract

Animals that exploit an extended phenotype (e.g., residences, resources, etc.) of other animals are called inquilines. Not strictly parasites, inquilines may nevertheless possess specialized traits that adapt them to particular dimensions of the extended phenotype of their “host”. These adaptations to host traits can in turn lead to fitness tradeoffs that restrict the host range of an inquiline such that shifts to novel hosts might trigger inquiline diversification. Speciation *via* host shifting has been studied in many animal parasites, but we know far less about the role of host shifts in inquiline speciation. *Synergus* (Hymenoptera: Cynipidae: Synergini) is a speciose but taxonomically-challenging group of inquilines that feed on the tissue of galls induced by oak gall wasps (Hymenoptera: Cynipidae: Cynipini). Currently too little is known about Nearctic *Synergus* diversity or host associations to evaluate whether and how host use affects their diversification. Here, we report on a large collection of *Synergus* reared from galls of 33 oak gall wasp species in the upper Midwestern United States. We integrated DNA barcodes, morphology, ecology, and phenology to delimit putative species of *Synergus* and describe their host ranges. We find evidence of at least 23 *Synergus* species associated with the 33 gall wasp hosts. At least five previously described *Synergus* species are each a complex of two to five species, while three species fit no prior description. We also find strong evidence that oak tree section and host gall morphology define axes of specialization for *Synergus*. Without over-interpreting our singlegene tree, it is clear that the North American *Synergus* have experienced several transitions among gall hosts and tree habitats and that host-use is correlated with reproductive isolation, though it remains too early to tell whether shifts to new hosts are the initiators of speciation events in *Synergus* inquilines of oak gall wasps, or if host shifts occur after reproductive isolation has already evolved.

## INTRODUCTION

Many plant-feeding insects are attacked by diverse and complex communities of natural enemies (Price 1980; Rodriguez and Hawkins 2000; Lewis et al. 2002). In general, these natural enemy communities are dominated by one or more trophic levels of parasitic wasps (parasitoids) that lay eggs into or onto their hosts, and whose larvae feed on the still-living host’s body. The particular ubiquity of parasitoids (Noyes 2012; Forbes et al. 2018) has led to much research into the mechanisms underlying their evolutionary success, with shifts by specialist parasitoids onto new host species frequently invoked as a potential driver of their speciation (e.g., Stireman et al. 2006; Forbes et al. 2009; Joyce et al. 2010; Kester et al. 2015; Hamerlinck et al. 2016; Hood et al. 2015). A related body of work has employed DNA sequence data to reveal cryptic host-specialization among insects – including parasitoids – initially thought to be oligophagous (Smith et al. 2006, 2008, 2011; Desneux et al. 2009; Forbes et al. 2012; Condon et al. 2008, 2014).

While parasitoids continue to be rewarding systems for evolutionary biologists interested in the relationship between host use and reproductive isolation, another group of insect natural enemies promises equally interesting insights but is far less well studied in this context. Inquilinous insects (inquilines) exploit the homes and/or use resources produced by other insects (Roskam 1979). Unlike parasitoids, which directly parasitize the host insect, inquilines “attack” the extended phenotype of their “host” insect, and their effect can range from a benign commensalism to a malignant kleptoparasitism. Though inquilines are common and taxonomically diverse in the context of some habitats (Sanver and Hawkins 2000), few studies to date have examined life history characters or host ranges for individual inquiline species (Stone et al. 1995; 2017), and fewer still have asked how ecology might influence their evolution and diversity (though see: Abrahamson et al. 2003; Blair et al. 2005).

Defined by their strategy of exploiting a habitat created by another animal, inquilines are a common associate of insects that induce the development of galls – fleshy, highly-structured growths of tissue – on plants (Sanver and Hawkins 2000). Larvae of gall-inducing insects benefit from the assured source of food that their gall provides and the protection from predators that concealment inside the gall offers (Egan et al. 2018). However, just as the gall ensures food and security for the developing galler, so too can it provide a high-quality source of food and protection to a variety of non-galling insects, including inquilines. Inquilines of gall wasps typically lay their eggs into the developing gall, after which their larvae consume the gall tissue. This can lead to malnourishment or starvation of the young galler (Sanver and Hawkins 2000). Inquilines of galls therefore straddle the nexus of herbivore and parasite, feeding on plant tissue but at the same time exploiting (and sometimes killing) their “host” insect. While not as speciose as parasitoids in most gall systems, they can often be among the most numerically dominant of gall associates, and some galls can host several different species of inquiline (Roininen and Price 1996; Forbes et al. 2015; A.W., personal observation).

*Synergus* (Hymenoptera: Cynipidae: Synergini) is a genus of inquiline wasps whose larvae feed on galls made on oak trees (*Quercus*; Fagaceae) by many of the ~1,400 oak gall wasp species (Hymenoptera: Cynipidae: Cynipini) (Pénzes et al. 2018). Most *Synergus* do not create their own primary galls (though see Abe et al. 2011; Ide et al. 2018). Instead, they induce secondary chambers in galls made by other cynipid wasps (Askew 1961; Pénzes et al. 2012). Some *Synergus* do not mortally injure the developing gall wasp, while others sever the nutrient supply and can be lethal to other gall inhabitants (Csoka et al 2005; Acs et al. 2010). *Synergus* are common across oak gall communities: often multiple species attack the same galler species (e.g., Melika 2006; Nazemi et al. 2008; Bird et al. 2013; Forbes et al. 2015; Weinersmith et al. 2020), though in many galls only one *Synergus* is known (Burks 1979) and in still others they may be replaced by other inquiline genera (e.g., Joseph et al. 2011).

Just a few studies have used molecular data to address the evolution and host ranges of *Synergus*, and all of these have focused on the Palearctic region. A three-gene (COI, cytb, and 28S) phylogeny of inquilines associated with oak gall wasps included 23 of the 30 described Western Palearctic *Synergus* species and generally supported the hypothesis that they are monophyletic relative to other gall-associated inquiline genera (Acs et al. 2010). Acs et al. (2010) also suggested that some species, including the apparent generalist *Synergus umbraculus*, might be amalgams of several morphologically convergent lineages. This phylogenetic work did not directly address species host ranges at the host plant or gall wasp species level. Two subsequent studies (Bihari et al. 2011, Stone et al. 2017) addressed the population genetics of one of the lineages of *S. umbraculus* identified by Acs et al. (2010; lineage 19). Both studies found a strong geographic signal without any strong signal of host-associated genetic differentiation. However, all host gall wasps in these studies were in genus *Andricus* and all collections save for one were from oaks in the same taxonomic section *(Quercus* sect. *Quercus*) (Bihari et al. 2011, Stone et al. 2017).

Though much more remains to be addressed relative to the role of host in diversification of Palearctic *Synergus*, even less is known about *Synergus* in the Nearctic despite the latter being more species-rich and with a far greater diversity of potential gall hosts (Pénzes et al. 2012; 2018). Seventy-eight species of *Synergus* have been described in the Nearctic (Lobato-Vila et al. 2019), though the total number is predicted to be far higher (Lobato-Vila and Pujade-Villar 2017). *Synergus* diversity is hypothesized to positively correlate with the diversity of potential gall hosts and the Nearctic has a greater diversity of both oaks (~300 species, compared to just ~40 species in the Western Palearctic; Stone et al. 2002; Lobato-Vila and Pujade-Villar 2017; Hipp et al. 2018) and oak-associated cynipid gall wasps (~700 species, compared with 140 in the western Palearctic; Stone et al. 2002).

Our primary goal in the current study is to characterize patterns of diversity, host range, and ecology for a subset of the Nearctic *Synergus*, with an eye towards understanding how changes in host use might be tied to their diversification. We employ a regionally-focused collection and *Synergus-rearing* effort from galls found growing on 15 species of oak trees, infer reproductive barriers using molecular barcodes, ecology, and morphology, and ask the following questions: 1) Do putative *Synergus* species that attack multiple different gall types harbor additional host- or habitat-associated genetic structure? 2) Along what axes (gall wasp species / gall morphology / tree species), if any, do *Synergus* specialize? We also use our limited molecular data to make a first inference regarding how Nearctic and Palearctic *Synergus* are related to one another.

## METHODS

### Notes on Taxonomy and Phylogeny of Synergus

Morphological characters historically used to sort *Synergus* into taxonomically relevant groups do not necessarily reflect characters shared due to common evolutionary histories. The Palearctic *Synergus* were initially divided into two major morphological groups (Mayr 1872): Section I, which are univoltine and usually not lethal to their associated galler, and Section II, which tend to be both bivoltine and lethal (Wiebes-Rijks 1979). The two sections also differed in patterns of microscopic punctures on their metasomal tergites. These sections were more recently revealed to be paraphyletic (Acs et al 2010). One of the potential reasons for the disconnect between morphological and molecular data is that *Synergus* can be morphologically cryptic as well as exhibit variance in morphology within species (Wiebes-Rijks 1979). In the Nearctic, where almost nothing is known about phylogenetic relationships, *Synergus* have been sorted into three morphologically-based groups primarily on the presence or absence of a smooth, shiny region on the mesopleuron (Lobato-Vila and Pujade-Villar 2017), but no molecular study has assessed this hypothesis, or the relationship between the Palearctic and Nearctic *Synergus* (Penzes et al. 2012).

### Collections

We employed a collection approach largely focused on galls in the Midwestern United States (Figure 1). We collected galls from August 2015 through October 2017. We recorded geographical location, date of collection, tree host and the species of gall wasp (based on gall characteristics, Weld 1959). We stored each collection of galls in an individual cup in an incubator that mimicked seasonal variations in temperature, humidity, and light/dark cycle. Each day, we checked all cups in the incubator and placed any emergent insects into 95% ethanol, recording the date of emergence. We identified each insect to family or genus using a variety of taxonomic resources (Goulet and Huber 1993; Gibson et al. 1997; Wahl 2015). In total we collected >24,000 galls, representing 103 gall wasp species (inferred using gall morphologies and tree host association based on Weld 1959).

**Figure 1.**
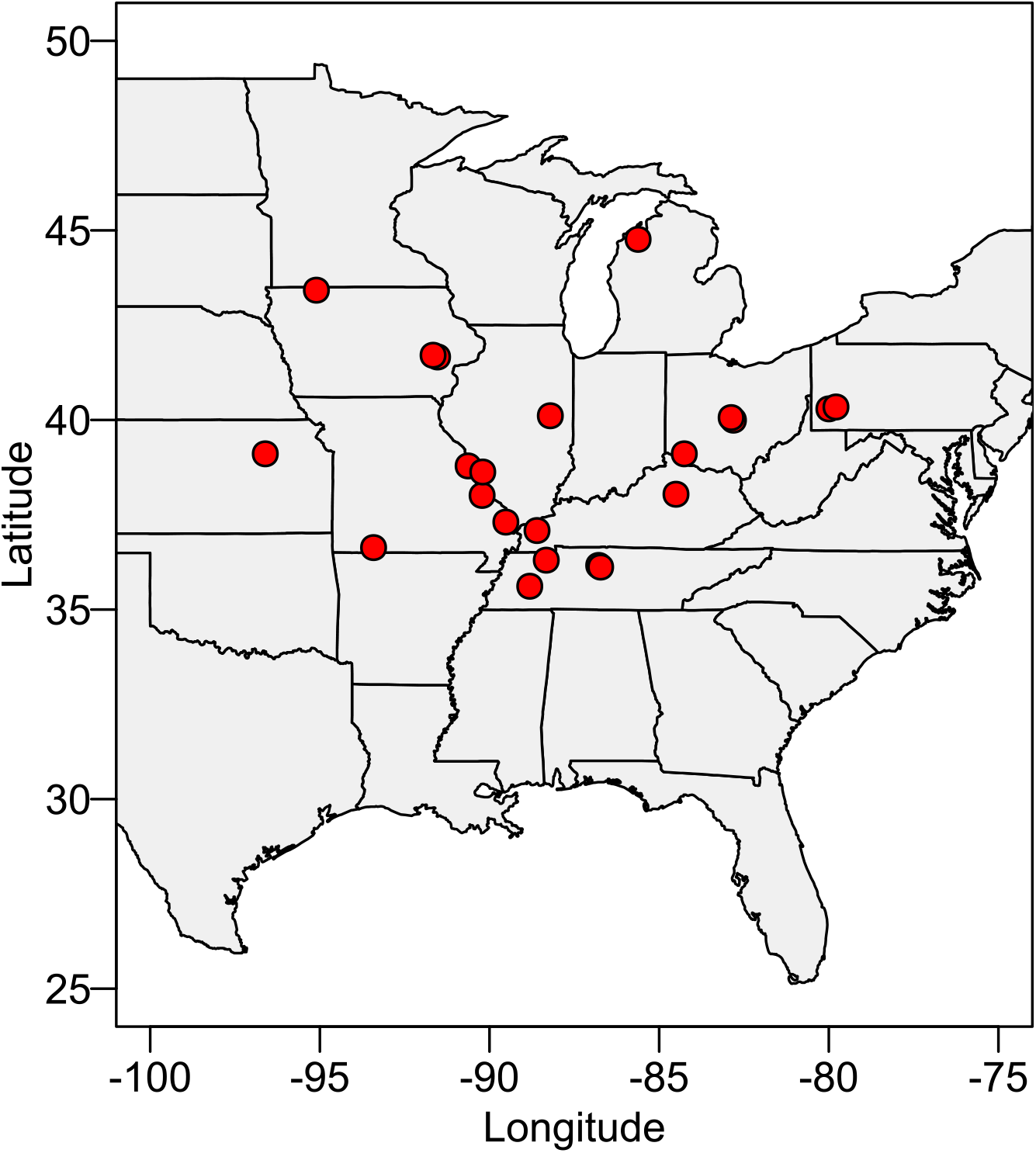
Map of gall collections that yielded *Synergus* samples used in this study. For a list of samples, locations, and tree- and gall-associations, see Supplemental Table 2.

### Identification of putative species

We first separated *Synergus* by gall association and then sorted individuals into morphotypes based on characters used by previous authors to distinguish species (e.g., color, mesopleural striation, antennal segmentation, patterns of syntergal punctures, etc.). Representatives from each morphotype were also identified to species – where possible – using keys by Gillette (1896), Lobato-Vila and Pujade-Villar (2017), and Lobato-Vila et al. (2019), or matched to described species based on morphological descriptions and records of host associations found in Fullaway (1911), McCracken and Egbert (1922), or Muesebeck et al. (1951). From each morphologically defined species, we chose individuals (usually females) to interrogate genetically. To allow for detection of possible cryptic diversity, we chose individuals that represented a diverse set of collection sites and from across each morphotype’s spectrum of eclosion dates. In cases where a *Synergus* morphotype emerged from the same gall wasp species but from different oak tree species, we also chose representatives from each tree habit. We photographed a single forewing and a profile of the body of each wasp used in genetic work (Supplementary figures S1-S46). We also pinned and deposited representatives of most morphotypes into the collection of the University of Iowa Museum of Natural History (Supplementary table 1).

For 96 samples extracted before 2017, we used a DNeasy Blood and Tissue kit (Qiagen) to destructively extract DNA from specimens. For another 55 *Synergus* extracted after 2017, we used a CTAB/PCI approach following the methods from Chen et al. (2010). The mtCOI region was amplified using the primers LepR15’ TAAACTTCTGGATGTCCAAAAAATCA 3’ and LepF1 5’ ATTCAACCAATACATAAAGATATTGG 3’ (Smith et al. 2008) We cleaned DNA using EXOI and SAP before Sanger sequencing both the forward and reverse sequence on an ABI 3730 in the University of Iowa’s Carver Center for Genomics. We aligned sequences in Geneious v.8.1.9 (Biomatters, Inc., San Diego, CA) and model-tested our sequences with jModelTest2 (Darriba et al 2012). We constructed phylogenetic trees using a Bayesian approach (MrBayes v3.2, Ronquist et al 2012) and a maximum likelihood approach (RAxML v8.2, Stamatakis 2014). Sequences were submitted to Genbank: accession numbers MT124785:MT124935 (see Supplemental table 2).

We generated molecular species hypotheses using two different but complementary approaches: Automatic Barcode Gap Discovery (ABGD; Puillandre et al. 2012) and PTP (Poisson Tree Processes model; Zhang et al. 2013). ABGD uses a sequence alignment to infer a confidence interval for genetic divergence within species. Ideally, a clear difference emerges between interspecific and intraspecific comparisons, defining a “barcode gap” for the sample set. PTP (Zhang et al. 2013) is a coalescence-based approach that requires a rooted phylogenetic tree as its input. We used the updated version bPTP, which uses branch lengths as proxies for substitutions and provides posterior Bayesian support values to each node describing the likelihood that descendants of that node belong to the same species. We used our Bayesian tree of *Synergus* samples (Supplemental Fig 47) as the input tree and ran 100,000 MCMC generations with a 10% burn-in. We checked likelihood trace plots to confirm burn-in was sufficient.

## RESULTS

### Summary of collections and species hypotheses

We reared 12,272 individual natural enemies from 88 different gall types. Of these natural enemies, 1,645 were *Synergus* wasps reared from galls of 33 gall wasp species (Supplemental Tables 3&4). Among these, we identified 14 *Synergus* morphospecies, 11 of which corresponded closely to previously described taxa (see “synthesis” below for details). A Bayesian tree of mtCOI from 148 individual *Synergus* (Figure 2; Supplemental Figure 47) demonstrates extensive phylogenetic structure corresponding to host gall wasp and oak section association. For ABGD, there was no clear barcode gap and thus we investigated two different partitions – one more liberal (more splitting) and one more conservative (more clumping). The liberal partition suggested 36 groups whereas the conservative partition suggested 21 groups (Figure 2; Supplemental Figure 48). The highest support solution for bPTP suggested 31 groups (Figure 2; Supplemental Figure 49).

**Figure 2.**
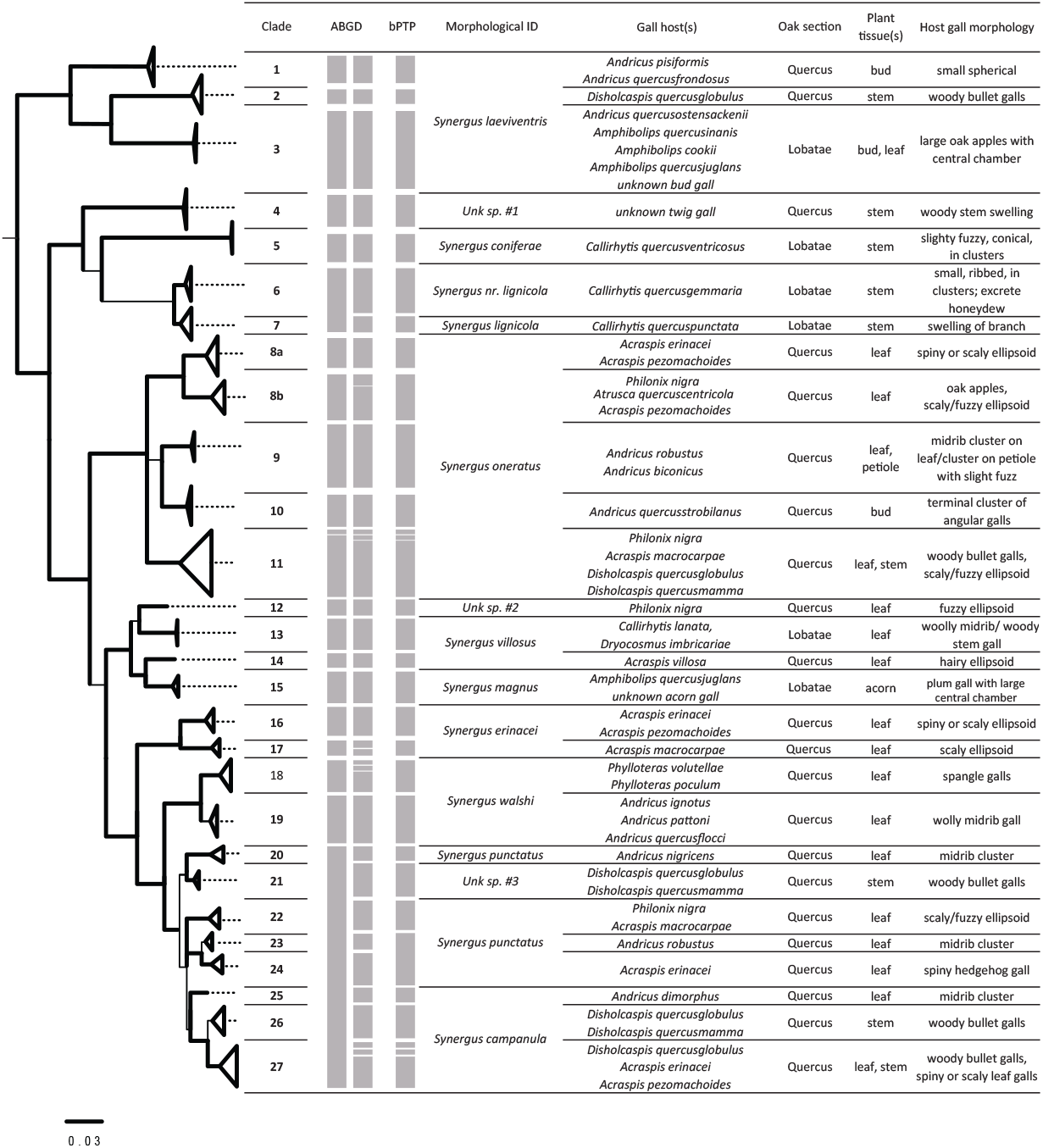
Overview of all data used in inferring *Synergus* species hypotheses for this study. Left: simplified mtCOI phylogeny of *Synergus* included in this study (see Supplemental Figure 47 for full tree). Bold branches indicate support ≥0.9 “Clade” describes putative species assignments based on the sum of information to the right of this column. Grey bars in ABGD (conservative and then liberal partitions) and bPTP columns indicate assignments of individuals into groups by these respective algorithms. “Morphological ID” refers to each collection’s similarity (or lack of similarity) to previously described species. “Gall host”, “oak section”, “plant tissue(s)”, and “Host gall morphology” refer to ecological characters for *Synergus* in each clade, and example photos of galls are shown in Supplemental Figure 51.

### Synthesis

We used concordance between morphology, ABGD, bPTP, and patterns of differing host association in sympatry to define a final count of putative species in our collections (Figure 2). Among the original 14 morphologically-defined species, we find a revised total of 23-27 species. In particular, at least five previously described *Synergus* species in our collections are each composed of at least two or more taxa, each with a more limited range of host galls than morphological species definitions initially implied. What follows is an account of each morphological “species” alongside a summary of its revised status in light of these new results. Species are named below in the order in which they appear in our mtCOI tree, from top to bottom (Figure 2). When a species was reared from >1 different gall types, we discuss apparent morphological or location-based similarities among these gall hosts. We also plot collection and emergence dates of all *Synergus* specimens to offer a sense of their phenology as it relates to host use (Figure 3).

**Figure 3.**
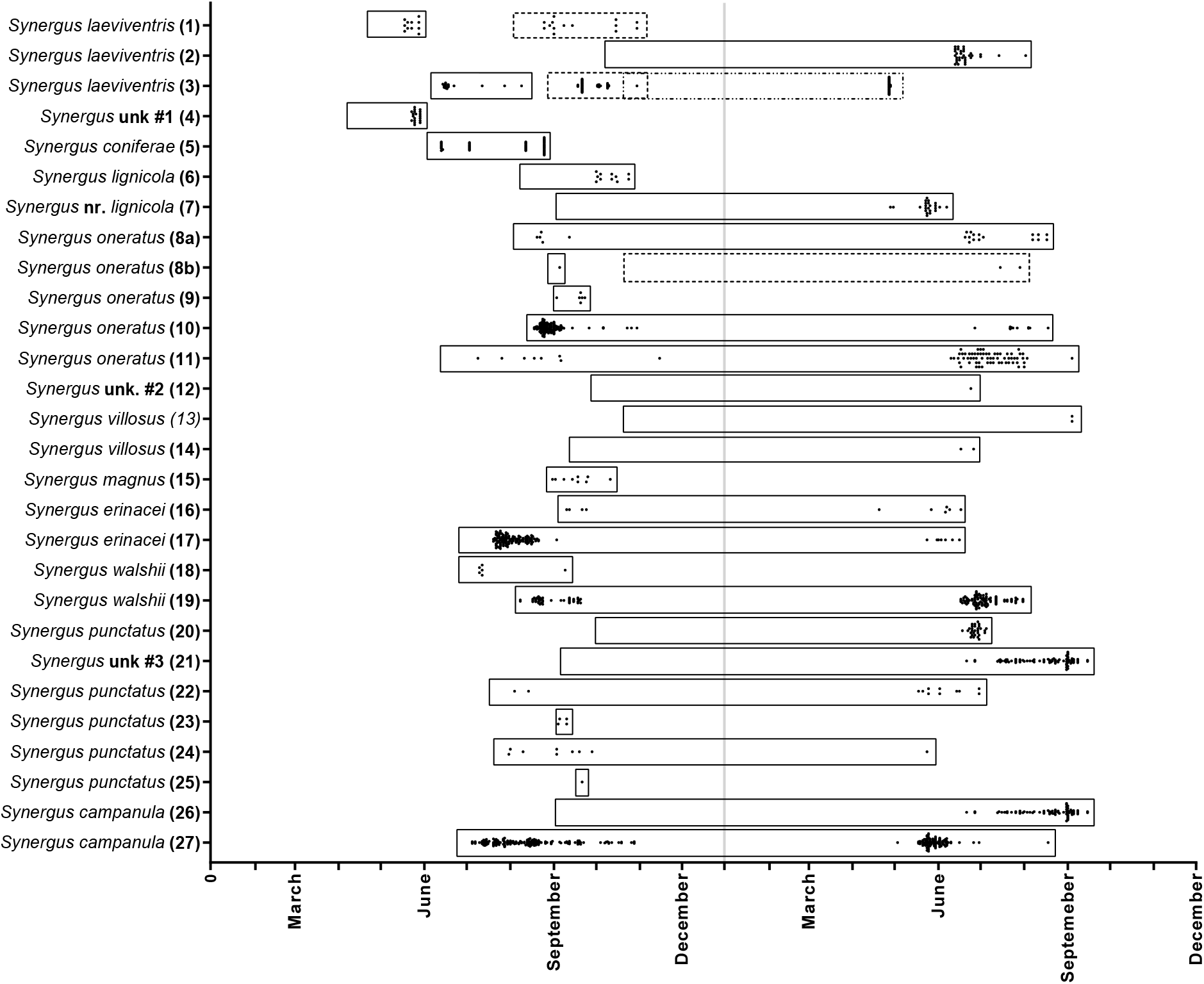
Collection and emergence dates for each of the 27 clades (putative species) of *Synergus* in this study. Dots indicate individual *Synergus* emergences; left-most margins of boxes demarcate the earliest gall collection that produced *Synergus* in each clade. In three cases, boxes with different border styles are used to indicate collection events from galls that occur at different times during the year, and which may indicate the use of temporally distributed gall hosts. Though galls were sometimes collected in different years, each collection was standardized to the year in which the gall first formed.

*Synergus laeviventris* (Osten Sacken) (Clades 1-3) (Figures S1-S6) is a complex of at least three species, differing from one another in their host gall use. One species (Clade 1) attacks the bud galls of *Andricus pisciformis* and *Andricus quercusfrondosus* on white oak (*Quercus alba*) and swamp white oak (*Quercus bicolor*) - both in oak section *Quercus*. A second species (clade 2) uses *Disholcaspis quercusglobulus* bullet galls on white oak (section *Quercus*) and a third (clade 3) attacks several different oak apple galls *(Andricus* and *Amphibolips*) found on red oak (*Quercus rubra*) and pin oak (*Quercus palustris*) – both section *Lobatae*. Previous rearing records associate *S. laeviventris* with the oak apple gall *Amphibolips quercusspongifica*, as well as from three bullet galls: *Disholcaspis quercusglobulus*, *Disholcaspis rubens*, and *Atrusca quercuscentricola* (Gillette 1896). Clades 1 and 3 are unusual among our collections in that we found evidence of two or three temporally distinct generations, each attacking different gall species (Figure 4).

*Synergus* unk. #1 (Clade 4) (Figures S7-S8) was reared from an unknown gall twig gall on a bur oak (*Quercus macrocarpa*; section *Quercus*). This *Synergus* ran closest to *Synergus batatoides* in the Gillette (1896) key, but its color and sculpturing did not match that description. *Synergus batatoides* are associated with *Callirhytis quercusbatatoides* stem galls in southern live oaks *(Quercus virginiana;* section *Virentes)*. Since we do not have collections of *S. batatoides* for comparison, it is not clear whether this is just a morphological variant of a species previous described only from the southern U.S. or a different species.

**Figure 4.**
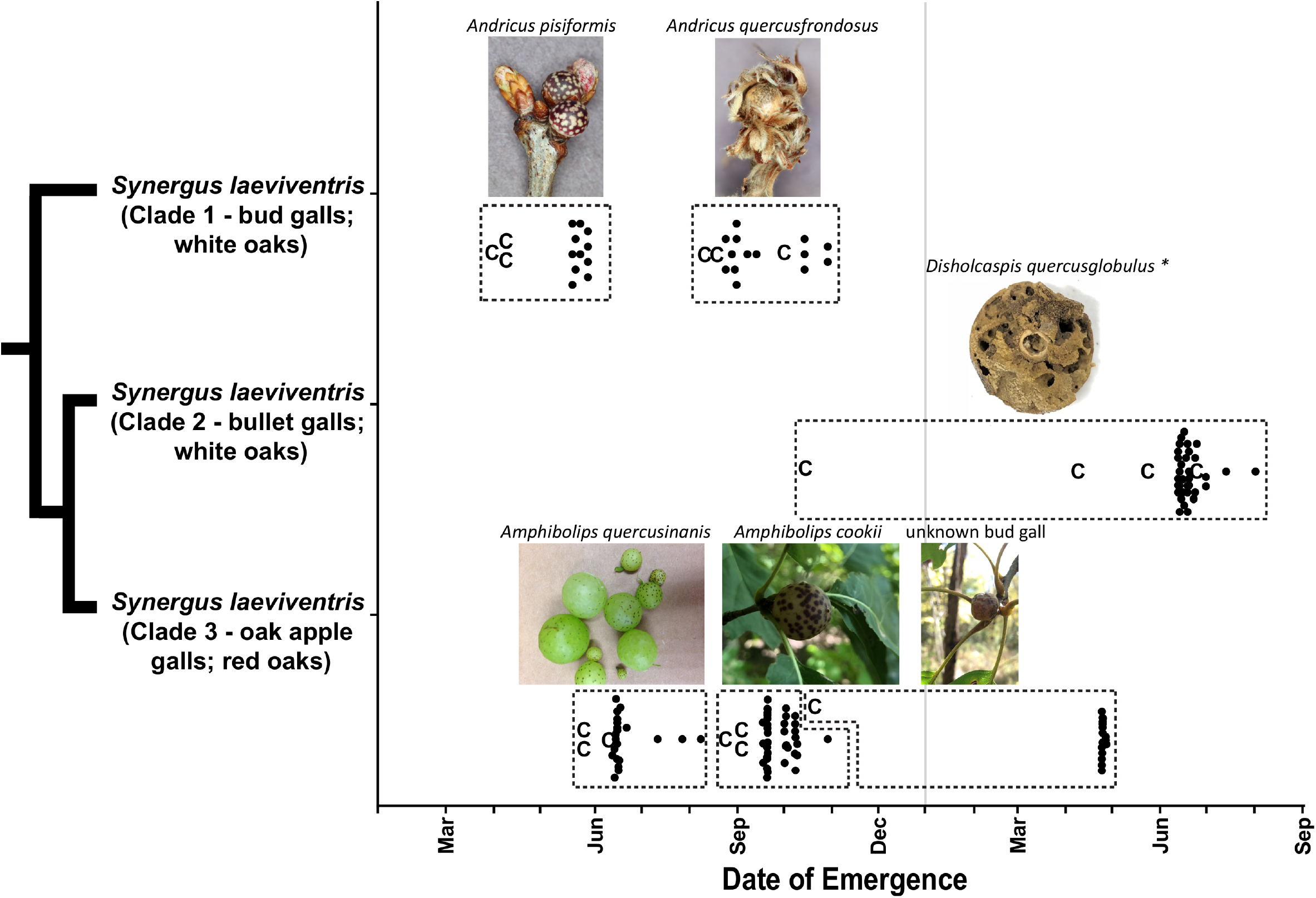
Collection dates, eclosion dates, and gall morphologies of three species previously collected under the name *Synergus laeviventris*. The relationships (as implied by mtCOI) among the three species are show in the tree at left. “**C**” denotes a collection from which at least one *S. laeviventris* later emerged, while black dots are individual *S. laeviventris* emergences. Boxes are used to isolate distinct collection/eclosion relationships from one another, which we interpret as evidence for discrete generations. Gall wasp species are labeled above the photos of their respective galls (* indicates the photo is a cross-section) Dates of eclosion are combined across the three years of collection.

*Synergus coniferae* Ashmead (Clade 5) (Figures S9-S10) was reared from *Callirhytis quercusventricosa* and previously from the same host by Ashmead (1885). We found no additional host or genetic structure within this species. Our samples were taken from shingle oaks (*Quercus imbricaria*; section *Lobatae*) in both Iowa City, IA, and St. Louis, MO. *Synergus lignicola* (Osten Sacken) (Clade 6) (Figures S11-S12) was reared from *Callirhytis quercusgemmaria* on red oaks (section *Lobatae*) from Traverse City, MI. These wasps closely match the description of *Synergus davisi* (Beutenmueller 1908) and were reared from the same host gall from which *S. davisi* was previously described, but a recent revision of some Nearctic *Synergus* (Lobato-Vila et al. 2019) synonymized *S. davisi* with *S. lignicola*. Our female *S.lignicola* specimens have an incomplete division in their final antennal segments, such that they can appear to have either 13 or 14 segments.

*Synergus* nr. *lignicola* (Clade 7) (Figures S11-S12) were reared from *Callirhytis quercuspunctata* collected on pin oaks in St. Louis, MO. This twig gall is the same host from which *S. lignicola* was previously reared (another known host is the morphologically similar *Callirhytis cornigera*) (Gillette 1896), but it differs in having 14 (instead of 13) female antennal segments, and a slight incision in the posterior-dorsal edge of its syntergite. At the more conservative partition, ABGD did not separate this species from *S. lignicola* (clade 6), though they were distinguished by both the bPTP method and the more liberal ABGD partition. The difference in morphology and host galls suggest that they are two different species, though a lack of a sympatric collection here allows for the possibility that differences are due to geographic isolation rather than reproductive isolating barriers.

*Synergus oneratus* (Harris) (Clades 8-11; Figures S15-S22) may contain as many as five species among the samples in our collection, each defined by COI sequence similarity, host association, and morphological variation. One species (clades 8a and 8b) was reared from *Acraspis* galls and other leaf galls on various white oaks (section *Quercus*) This species was split into two additional groups by the less conservative ABGD partition but since both groups contained *Synergus* from the same *Acraspis pezomachoides* collection in Peducah, KY, this split does not appear to be supported by ecology or geography. A second species (clade 9) was reared from *Andricus biconius* and *Andricus robustus* in post oaks (*Quercus stellata*; section *Quercus*), and a third species (clade 10) was reared from *Andricus quercusstrobilanus* in bur, white, and swamp white oaks (section *Quercus)* A fourth species (clade 11) was reared from a variety of stem and leaf galls on oaks in section *Quercus*. Previous taxonomic work has already suggested that *S. oneratus* species contains two geographic varieties differing in the amount of black on the mesonotum: *S. oneratus oneratus* (some yellow) in the Midwest and *S. oneratus coloradensis* (entirely black) in the west (Gillette 1896). The discovery of *S. oneratus oneratus* in California (Fullaway 1911; McCracken and Egbert 1922) indicated a more complex distribution. Here, Clades 10 and 11 (Figure S19, S21) appear to better match the description of *S. oneratus oneratus*, while Clade 8 (Figures S15, S17) is closer to *S. oneratus coloradensis*. No females of clade 9 were available for this comparison.

*Synergus* unk. #2 (Clade 12) (Figure S23) was represented by a single male reared from a *Philonix nigra* gall on leaves of bur oak in Konza, KS (clade 12). Because many keys do not address males, the failure to attach a name to this species may not indicate that is undescribed.

*Synergus villosus* Gillette (Clades 13 & 14) (Figure S24-S25) includes two distinct species. One species (clade 13) was reared from *Acraspis villosa*, a fuzzy leaf gall, on bur oak (section *Quercus*) in Urbana, IL. A second species (clade 14) was reared from *Callirhytis quercuslanata* and *Dryocosmus imbricariae*, a fuzzy leaf gall and a stem bullet gall, respectively, both on red oak (section *Lobatae)* in Tiffin, IA. Some caution should be used in definitively separating clades 13 and 14, as collections were from different states and differences could be due to geography. The original description of *S. villosus* (Gillette 1896) was from *A. villosa* galls collected in Iowa and the morphology of these wasps exactly match our Illinois wasp, while the samples from red oak showed them to have entirely yellow mesonota, which differs from the bur oak collection and the original *S. villosus* description (thorax entirely black).

*Synergus magnus* Gillette (Clade 15) (Figure S26) was reared from *Amphibolips quercusjugulans* leaf galls and from unidentified acorn galls on black oaks (*Quercus velutina*) and red oaks, respectively (both section *Lobatae*). This species was previously reared from an *Amphibolips cookii* gall (Gillette 1896).

*Synergus erinacei* Gillette (Clades 16 & 17) (Figures S27-S29) contains either one or two different species. One putative species (clade 16) was reared from *Acraspis erinacei* and *A. pezomachoides* leaf galls on white oaks (section *Quercus*). The second (clade 17) was reared from *Acraspis macrocarpae* on bur oaks (section *Quercus*). The two *S. erinacei* clades are also separated by geography, so a definitive conclusion regarding species status requires additional work.

*Synergus walshii* Gillette (Clades 18 & 19) (Figures S30-S33) contains two putative species. One (clade 18) was reared from galls of two *Phylloteras* species on bur oak and swamp white oak (section *Quercus*), the other (clade 19) from various *Andricus* leaf galls on bur, white, and post oaks (section *Quercus)*. Previous collections of *S. walshii* (Gillette 1896) were from *Andricus quercusflocci* galls, one of the clade 19 hosts.

*Synergus punctatus* Gillette (Clades 20, 22-25) (Figures S34-S35; S37-S41) may contain up to five distinct species. The first species (clade 20) was reared from leaf galls of *Andricus nigricens* on swamp white oak (section *Quercus*). A second species (clade 22) was reared from *Acraspis* and *Philonix* leaf galls on white, bur, and chinquapin oaks (*Quercus muehlenbergii*) (section *Quercus*) and females had some color variation, with thorax and abdomen color ranging from black to a chestnut brown. Two additional putative species – both with a mix of black- and brown-bodied individuals – were reared from galls of *Andricus robustus* (clade 23) and *Acraspis erinacei* (clade 24) galls on post and white oaks, respectively. We are more tentative about definitively splitting clades 23 and 24 due to conflation of genetic and host differences with geographic distance.

*Synergus* unk #3 (Clade 21) (Figure S36) was reared from *Disholcaspis* bullet galls on branches of white, swamp white, and post oaks (section *Quercus*) and differed from *S. punctatus* by having females with mostly chestnut-brown heads, and <1/3 of the posterior edge of their syntergite covered with punctures.

*Synergus campanula* Osten Sacken (Clades 25-27) (Figures S42-S46) contains between two and three species. One species (clade 25) was collected from clustered midrib leaf galls of *Andricus dimorphus* on dwarf chinquapin oak (*Quercus prinoides*; section *Quercus*). The other species (or two) (clades 26 & 27) were reared from a mix of leaf and stem galls from white, swamp white, and post oaks (section *Quercus*).

## DISCUSSION

### Nearctic Synergus diversity and taxonomy

One immediate lesson from this study is that the Nearctic *Synergus* harbor both cryptic and undiscovered species. For five of 11 previously-named species in our collections, we find evidence of additional genetic, morphological, and/or ecological structure indicative of >1 species. Three other putative species apparently fit no previous description. Because this was a geographically-limited survey, it thus seems very likely that many more undiscovered species exist in North America, both as members of morphologically-cryptic assemblages, and as undescribed species. Indeed, when recent authors have focused on the Nearctic, they have consistently added new species (e.g., Lobato-Vila & Pujade-Villar 2017; Lobato-Villa 2018; 2019). The incompleteness of the record is perhaps unsurprising as most taxonomic and phylogenetic work has emphasized the *Synergus* of the western Palearctic (Penzes et al. 2012) while the most comprehensive list of Nearctic *Synergus* species is now >40 years old (Burks 1979).

A taxonomic revision of the Nearctic *Synergus* that incorporates both molecular and ecological data appears warranted, and, though such a revision is beyond the scope of this paper, it might be informed by some of our findings. In particular, one morphological character previously suggested as useful for organizing the Nearctic fauna may have some significance. Gillette (1896) split the genus into three “natural” groups based on whether females of each species have 13, 14, or 15 segments in their flagella. Among our collections, we found only 14- and 15-segmented females (with *S. lignicola* being indeterminately 14-segmented), but all 15-segmented females (clades 12-15) grouped together on the mtCOI tree (Figure 2; Supplemental Figures 47 and 50), suggesting a common ancestry among these samples. While we acknowledge caveats about inferring too much from a single short mitochondrial sequence (Funk and Omland 2003; and more on this below), this may be an early indication that antennal segment number is an informative character. To assist future efforts in this regard, we include supplemental profile and forewing pictures of wasps from each clade (Supplemental figures S1-S46) and have deposited examples of most clades into the collection of the University of Iowa Museum of Natural History.

### Host ranges and axes of specialization

Our primary motivation for this study was to evaluate the extent to which inquilines specialize on aspects of their host environments, which then would stand as an initial, indirect assessment of how changes in host use might result in lineage divergence. Evidence from our study suggests that Nearctic *Synergus* specialize on aspects of host plant identity, gall morphology, and/or gall phenology. One obvious axis of specialization was host tree section. Though we made large collections of galls from several oak species in both section *Quercus* (white oaks) and section *Lobatae* (red oaks), we found no evidence of a *Synergus* species that was reared from both sections. This pattern conforms to previous studies, which have suggested *Synergus* may often be specific to closely related oaks species (Pujade-Villar 2003). Other Cynipidae, including the gall wasp hosts of these *Synergus*, also show fidelity to oaks with similar chemistry (which generally means more closely related oaks; Abrahamson et al. 1998, 2003).

Less specialization was evident at finer levels of plant taxonomic organization. For example, *Synergus* species reared from galls on white oak (*Quercus alba*) were also often reared from galls on swamp white oak (*Quercus bicolor*), bur oak (*Quercus macrocarpa*), or post oak (*Quercus stellata*) – all North American members of section *Quercus*, the white oaks. Specialization on oak section rather than oak species may explain why prior assessments of host plant specialization in the Palearctic *Synergus* – where most samples have been from the same oak section – have not resolved strong genetic structure associated with host plant (e.g., Bihari et al. 2011, Stone et al. 2017). Given that *Synergus* in this study specialize on either red and white oaks, it will be interesting in the future to include *Synergus* associated from oaks in section *Virentes* (the live oaks), another North American group which overlaps geographically with the southern ranges of some of the oaks in this study.

While *Synergus* did not appear to specialize at the oak species level (below the section level), they did specialize on galls of different morphology. Though our haphazard collections give us an incomplete picture of all hosts attacked by each species, most species tended to favor galls that shared similar characteristics (Figure 2, Supplemental Figure 51). Most strikingly, even when we reared *Synergus* from different gall hosts across the course of the year, each generation nevertheless conserved elements of gall morphology. The Clade 1 *S. laeviventris* wasps, for example, emerged in late May from small bud galls of *Andricus pisiformis* collected in early April, and then adults emerged again in the fall from a second small bud gall (*A. quercusfrondosus)* collected in August and September (Figure 4). Adults of this species may then oviposit into pre-winter bud galls of *A. pisiformis* or perhaps attack a third overwintering host that we did not collect in this study. Similarly, we captured three discrete generations of *S. laeviventris* Clade 3 wasps, each attacking oak apple galls containing a central cell surrounded by radiating fibers, but each made by different galler species (Figure 4).

Conservation of host gall morphology suggests that *Synergus* specialization conforms to the “Enemy Hypothesis” (Bailey et al. 2009). That is: galls with similar qualities will exclude certain natural enemies and therefore similar galls will tend to be attacked by the same or similar species (Ward et al. 2019). Brookfield (1972) similarly suggested that inquilines and parasitoids of galls depend more on gall morphology than other factors such as gall wasp species or tree host (note, however, that this last is belied by our data). Because the evolutionary relationships among the Nearctic gall wasps themselves are not yet known, it is not possible to distinguish gall morphology from gall wasp phylogeny. A phylogeny of North American gall wasps will be helpful in this respect.

Phenology has also been previously considered as an important axis of adaptation and specialization for *Synergus* because some developing galls may offer only a limited “window of opportunity” during which an inquiline might successfully oviposit (Penzes et al. 2012). The *S. laeviventris* Clades 1 and 3 wasps described above appear to fit this pattern, with each of their respective two or three generations attacking different galls, presumably during some window of their early development (Figure 4). In contrast to this pattern, the long (July-October) emergence schedule of *S. punctatus* Clade 24 wasps from *A. erinacei* galls (Figure 5) suggests that while these *Synergus* undergo multiple generations per year, they are capable of attacking *A. erinacei* galls for much of the time that these galls occur on leaves. *Synergus punctatus* even appears to undergo a period of diapause during the winter that ends shortly before the next generation of *A. erinacei* appears on leaves again.

**Figure 5.**
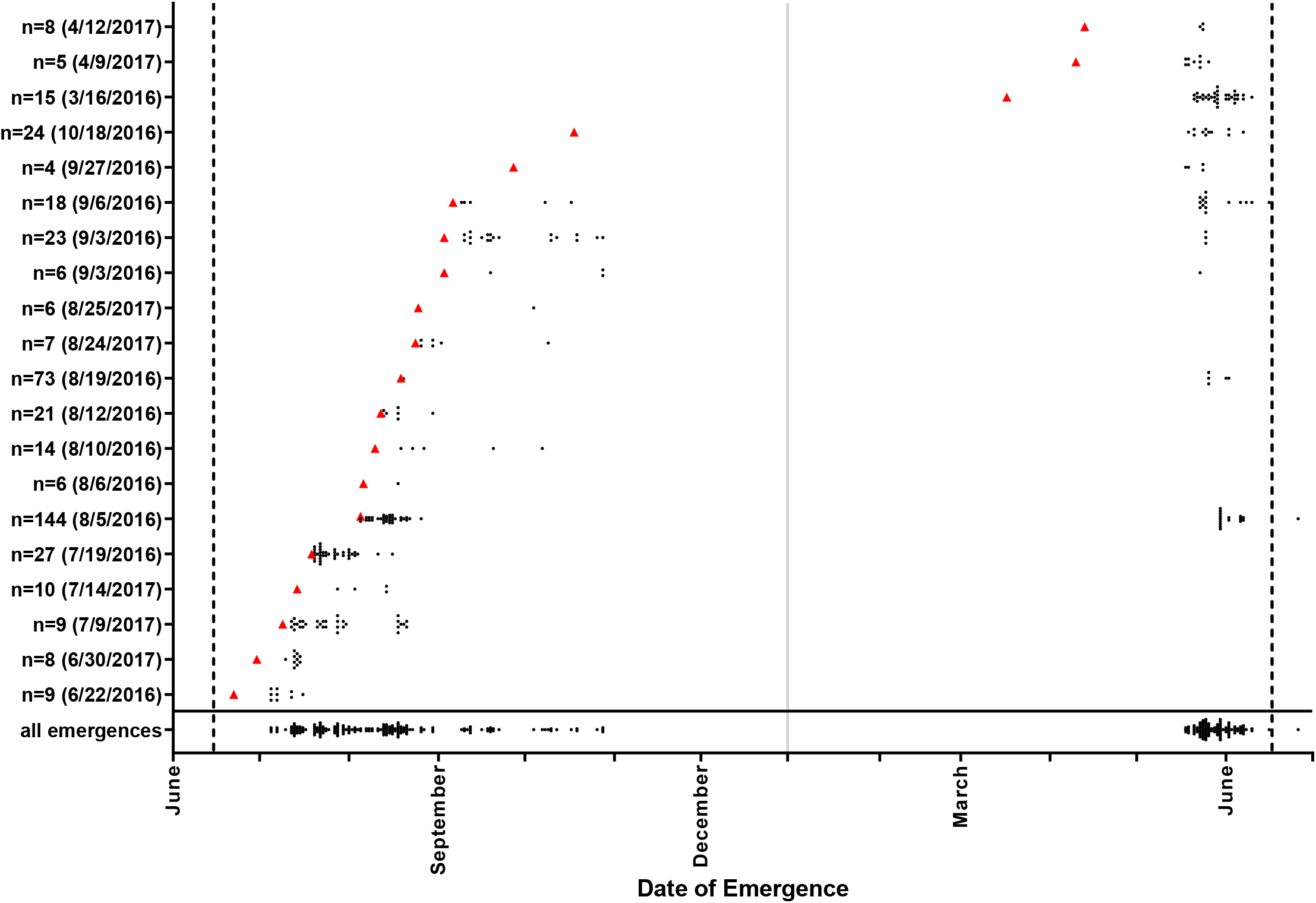
Emergence dates of adult *Synergus punctatus* (clade 24) from 20 separate collections of *Acraspis erinacei* galls, organized by their calendar date of collection. Vertical dashed lines indicate the earliest date when we observed young *A. erinacei* galls on trees during this study. Red triangles denote the date of each collection, with the three collections in year two representing the previous year’s leaves collected from the ground beneath trees.

Characterizing *Synergus* host ranges and their relative level of specialization has important connotations for conservation and pest control. Oak gall wasps have been transported into nonnative ranges (Walker et al. 2002; Prior and Hellmann 2010), with some classified as invasive due to their impact on trees (Buffington et al. 2016; Davis 2017). *Synergus*, because they are often the most abundant enemy of some gall species, may be particularly important in the control of alien gallers (Schönrogge et al. 1996, 2000), and their host breadth could inform likelihood of colonization. For instance, knowledge about host gall characters might allow for biocontrol efforts that employ native *Synergus* rather than introducing non-native parasites or inquilines (though the latter can also be successful: Moriya et al. 1989).

### Implications for Synergus evolution

One leading explanation for why parasitoid wasps are so diverse is that that adaptations to new insect hosts may lead to host-associated differentiation and speciation (Stireman et al. 2005; Feder and Forbes 2010; Kaiser et al. 2015; König et al. 2015). When a parasitoid adopts a new host, trade-offs associated with adaptation to the new host can result in reproductive isolation from individuals attacking the ancestral host (Hood et al. 2015). We know surprisingly little about whether the same ecological differences can drive lineage divergence in insect inquilines: one exception is the tumbling flower beetle *Mordellistena convicta* (Coleoptera: Mordellidae), which feeds on the galls of two lineages of *Eurosta solidaginis* gall-flies (Diptera: Tephritidae) on goldenrods (*Solidaginis*; Compositae). Life history and molecular work has shown that both *E. solidaginis* and *M. convicta* have formed host-associated lineages on two different species of goldenrod, suggesting that speciation has “cascaded” across trophic levels (gall former to inquiline) (Abrahamson et al. 2003; Eubanks et al. 2003; Rhodes et al. 2012). This beetle system offers a compelling hint that ecological changes might lead to lineage divergence in inquilines, but missing is a broader study of ecological and host use patterns within an entire inquiline clade.

What do we learn about the role of host shifts in *Synergus* evolution? Though not a comprehensive phylogenetic history of these species, our mtCOI tree of Nearctic *Synergus* (Figure 2, Supplemental Fig 47, 50) nevertheless demonstrates that transitions between host tree sections (*Quercus* to *Lobatae*), between gall wasp species, and between gall morphologies are all common. This is not a definitive signature that host shifts drive speciation in *Synergus*, but the broad correlation between ecological changes and the evolution of reproductive isolation suggests this as a leading hypothesis. Transitions among host wasps, galls, and tree sections all represent changes that – to differing extents – require the evolution of novel behaviors, life histories, and strategies for circumventing host defenses. The accumulation of such divergent characters in the context of different habitats has been shown to directly (Craig et al. 1993; Forbes et al. 2005; Fujiyama et al. 2013; Doellman et al. 2019; Hood et al. 2019) and indirectly (Nosil and Hohenlohe 2012) lead to the evolution of reproductive isolation in many different insect systems.

An alternative to the above hypothesis is that ecological divergence among *Synergus* occurs after reproductive isolation evolves between lineages. For instance, perhaps *Synergus* speciate in the absence of divergent ecological selection (e.g., Nosil and Flaxman 2011), and then later shift into new hosts or habitats upon secondary contact because partitioning of environments facilitates coexistence of ecologically similar sympatric species (Dufour et al. 2017). One advantage of our renewed exploration of the Nearctic *Synergus* is that it suggests taxa for further study that may allow the disentanglement of these competing hypotheses. For instance, clades where individuals with little or no mtCOI differentiation have been reared from galls of different morphologies (e.g., Clades 3, 9, 27) could be interrogated further. If experimental tests revealed significant host-associated reproductive isolation and/or population genetic analyses suggested evidence of host-associated genetic differentiation, these might support a hypothesis that difference in host use are contributing to divergence. Similarly, as new samples of *Synergus* from across a wider geographic area are added to this dataset, an increasing role for geographic isolation might emerge between sister species.

### Notes on the Relationship between Nearctic and Palearctic Synergus

This first genetic analysis of a large number of Nearctic *Synergus* allows for a preliminary assessment of their relationships to the better-studied Palearctic *Synergus*. We combined our mtCOI sequences to the previously published dataset of Palearctic *Synergus* mtCOI sequences from Aces et al (2010) and generated Bayesian and Maximum likelihood trees following previously described methods. Both methods generated trees with the same topology: the Nearctic *Synergus* formed two distinct clades while the Palearctic *Synergus* formed one monophyletic clade (Figure 6). However, at this time we cannot conclusively comment on the evolutionary relationship amongst these clades due to the tree being based on one gene. Future multi-locus analyses promise to reveal additional insights into these relationships.

**Figure 6.**
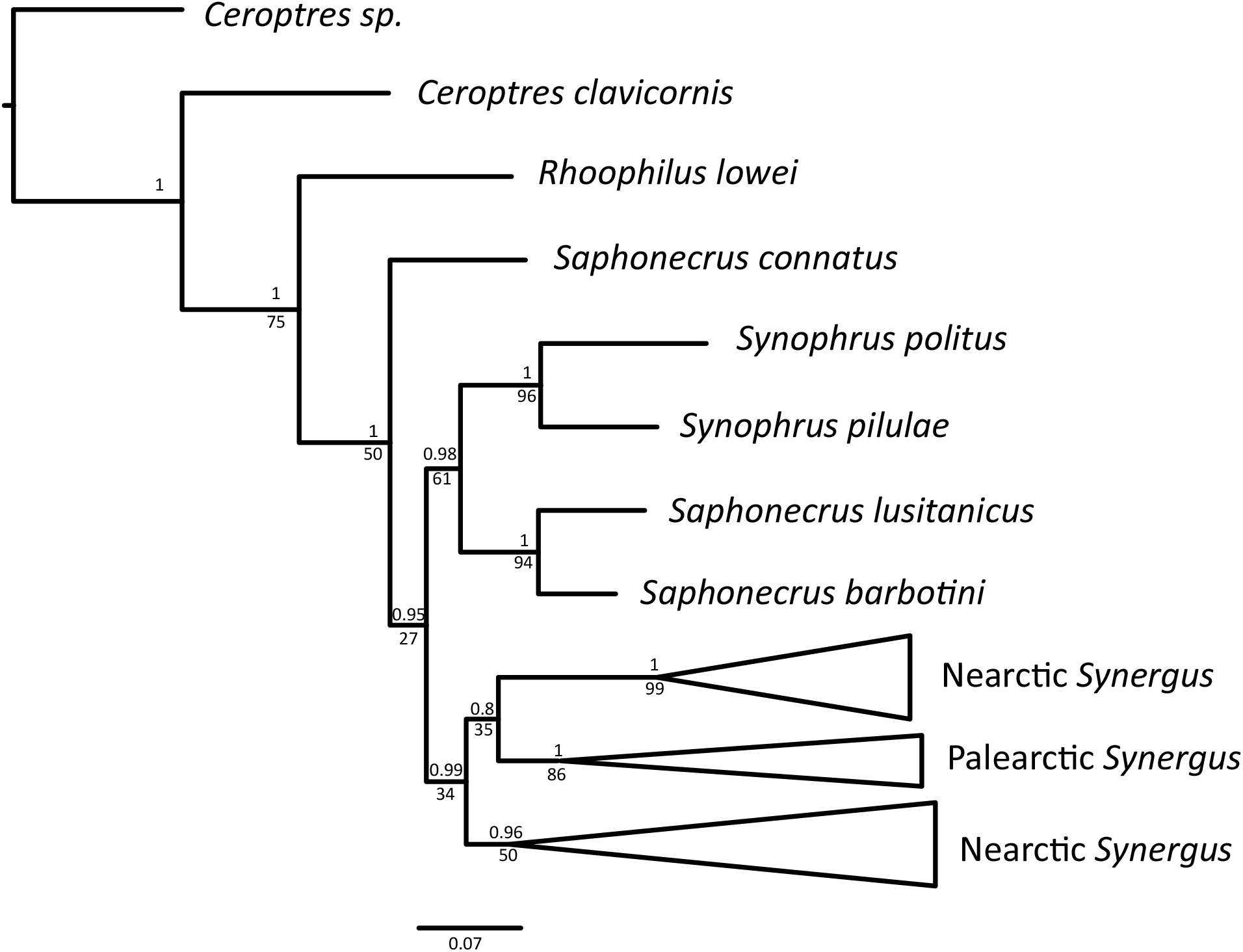
Phylogenetic tree of Nearctic *Synergus* from this study (shown here collapsed into two clades) combined with sequences from Acs et al 2010. Values above the branches represent Bayesian posterior probabilities and the values below the branch are ML bootstrap values. See Supplemental Figures 52 and 5 for full trees.

## Acknowledgements

Funding for collections was provided to A.K.G.W. and S.I.S. from the Center for Global and Regional Environmental Research at the University of Iowa. Robin Bagley, Will Carr, Sarah Duhon, Alaine Hippee, Kyle McElroy, Daniel McGarry, Kevin Neely, Monzer Shakally, Eric Tvedte, Joseph Verry, and Caleb Wilson helped collect galls. Irene Lobato-Vila provided helpful context regarding possible morphological variation within *Synergus lignicola* and its synonymy with *Synergus davisi*.

## Notes

### Competing Interest Statement

The authors have declared no competing interest.

## References

Abe, Y., Ide, T., & Wachi, N. (2011). Discovery of a New Gall-Inducing Species in the Inquiline Tribe Synergini (Hymenoptera: Cynipidae): Inconsistent Implications from Biology and Morphology. Annals of the Entomological Society of America, 104(2), 115–120.

Abrahamson, W. G., Blair, C. P., Eubanks, M. D., & Morehead, S. A. (2003). Sequential radiation of unrelated organisms: the gall fly *Eurosta solidaginis* and the tumbling flower beetle *Mordellistena convicta*. Journal of Evolutionary Biology, 16(5), 781–789.

Abrahamson, W. G., Melika, G., Scrafford, R., & Csóka, G. (1998). Gall-inducing insects provide insights into plant systematic relationships. American Journal of Botany, 85(8), 1159–1165.

Ács, Z., Challis, R. J., Bihari, P., Blaxter, M., Hayward, A., Melika, G.,… & Schönrogge, K. (2010). Phylogeny and DNA barcoding of inquiline oak gallwasps (Hymenoptera: Cynipidae) of the Western Palaearctic. Molecular Phylogenetics and Evolution, 55(1), 210–225.

Ashmead, W. H. (1885). A bibliographical and synonymical catalogue of the North American Cynipidae, with description of new species. Transactions of the American Entomological Society 12(3-4), 291–304.

Askew, R. R. (1961). On the biology of the inhabitants of oak galls of Cynipidae (Hymenoptera) in Britain. Transactions of the Society for British Entomology, 14(11), 237–268.

Bailey, R., Schönrogge, K., Cook, J. M., Melika, G., Csóka, G., Thuróczy, C., & Stone, G. N. (2009). Host Niches and Defensive Extended Phenotypes Structure Parasitoid Wasp Communities. PLoS Biology, 7(8), e1000179.

Beutenmüller, W. (1907). Notes on a few North American Cynipidae, with descriptions of new species. Bulletin of the American Museum of Natural History, 23, 463–466.

Bihari, P., Sipos, B., Melika, G., Fehér, B., Somogyi, K., Stone, G. N., & Pénzes, Z. (2011). Western Palaearctic phylogeography of an inquiline oak gall wasp, *Synergus umbraculus*. Biological Journal of the Linnean Society, 102(4), 750–764.

Bird, J. P., Melika, G., Nicholls, J. A., Stone, G. N., & Buss E. A. (2013). Life history, natural enemies, and management of *Disholcaspis quercusvirens* (Hymenoptera: Cynipidae) on live oak trees. J. Econ. Entomol., 106(4), 1747–1756.

Blair, C. P., Abrahamson, W. G., Jackman, J. A., & Tyrrell, L. (2005). Cryptic speciation and host-race formation in a purportedly generalist tumbling flower beetle. Evolution, 59(2), 304–316.

Brookfield, J. F. (1972). The Inhabitants (Hymenoptera: Cynipidae, Chalcidoidea) Of the Cynipidous Galls of *Quercus Borealis* In Nova Scotia. The Canadian Entomologist, 104(7), 1123–1133.

Buffington, M. L., Melika, G., Davis, M., & Elkinton, J. S. (2016). The Description of *Zapatella davisae*, New Species, (Hymenoptera: Cynipidae) a Pest Gallwasp of Black Oak *(Quercus velutina)* in New England, USA. Proceedings of the Entomological Society of Washington, 118(1), 14–27.

Burks, B. D. (1979). Superfamily Cynipoidea. In K.V. Krombein, P.D. Hurd, and D.R. Smith, (Eds.), Catalog of Hymenoptera in America north of Mexico, volume 1: Symphyta and Apocrita (Parasitica) (pp. 1045–1107). Smithsonian Institution Press, Washington District of Columbia, United States of America.

Chen, H., Rangasamy, M., Tan, S.Y., Wang, H., & Siegfried, B.D. (2010). Evaluation of five methods for total DNA extraction from Western Corn Rootworm Beetles. PLoS one, 5, e11963.

Condon, M. A., Scheffer, S. J., Lewis, M. L., & Swensen, S. M. (2008). Hidden neotropical diversity: greater than the sum of its parts. Science, 320(5878), 928–931.

Condon, M. A., Scheffer, S. J., Lewis, M. L., Wharton, R., Adams, D. C., & Forbes, A. A. (2014). Lethal interactions between parasites and prey increase niche diversity in a tropical community. Science, 343(6176), 1240–1244.

Craig, T. P., J. K. Itami, W. G. Abrahamson, and J. D Horner. (1993). Behavioral evidence for host-race formation in *Eurosta solidaginis*. Evolution, 47, 1696–1710.

Csóka, G., Stone G. N., & Melika G. (2005). Biology, ecology and evolution of gall-inducing Cynipidae. In A. Raman, C.W. Schaefer, and T. M. Withers, (eds.), Biology, ecology and evolution of gall-inducing arthropods. (pp. 569–636) Science Publishers, Inc. Enfield, New Hampshire, USA

Darriba, D., Taboada, G. L., Doallo, R., & Posada, D. (2012). jModelTest 2: more models, new heuristics and parallel computing. Nature methods, 9(8), 772. doi:10.1038/nmeth.2109

Davis, M. (2017). Biology, Molecular Systematics, Population Dynamics and Control of a Stem Gall Wasp, Zapatella Davisae (Hymenoptera: Cynipidae) (Doctoral dissertation, University of Massachusetts Libraries).

Desneux, N., Starý, P., Delebecque, C. J., Gariepy, T. D., Barta, R. J., Hoelmer, K. A., & Heimpel, G. E. (2009). Cryptic species of parasitoids attacking the soybean aphid (Hemiptera: Aphididae) in Asia: *Binodoxys communis* and *Binodoxys koreanus* (Hymenoptera: Braconidae: Aphidiinae). Annals of the Entomological Society of America, 102(6), 925–936.

Doellman, M. M., Egan, S. P., Ragland, G. J., Meyers, P. J., Hood, G. R., Powell, T. H., … & Feder, J. L. (2019). Standing geographic variation in eclosion time and the genomics of host race formation in *Rhagoletis pomonella* fruit flies. Ecology and Evolution, 9(1), 393–409.

Dufour, C. M., Herrel, A., & Losos, J. B. (2017). Ecological character displacement between a native and an introduced species: the invasion of *Anolis cristatellus* in Dominica. Biological Journal of the Linnean Society, 123(1), 43–54.

Egan, S. P., Hood, G. R., Martinson, E. O., & Ott, J. R. (2018). Cynipid gall wasps. Current Biology, 28(24), R1370–R1374.

Eubanks, M. D., Blair, C. P., & Abrahamson, W. G. (2003). One host shift leads to another? Evidence of host-race formation in a predaceous gall-boring beetle. Evolution, 57(1), 168–172.

Feder J. L., Forbes A. A. (2010). Sequential speciation and the diversity of parasitic insects. Ecological Entomology, 35, 67–76.

Forbes A. A., Bagley R. K., Beer M., Hippee A. C., & Widmayer H. A. (2018). Quantifying the unquantifiable: why Hymenoptera - not Coleoptera - is the most speciose animal order. BMC Ecology, 18:21.

Forbes, A. A., Fisher, J., Feder, J. L. (2005). Habitat avoidance: Overlooking an important aspect of host-specific mating and sympatric speciation? Evolution. 59(7), 1552–1559.

Forbes A. A., Hood G. R., Hall M. C., Lund J., Izen R., Egan S. P., & Ott J. R. (2015). Parasitoids, hyperparasitoids, and inquilines associated with the sexual and asexual generations of the gall former, *Belonocnema treatae* (Hymenoptera: Cynipidae). The Annals of the Entomological Society of America, 109, 49–63.

Forbes, A. A., Satar, S., Hamerlinck, G., Nelson, A. E., & Smith, J. J. (2012). DNA barcodes and targeted sampling methods identify a new species and cryptic patterns of host specialization among North American Coptera (Hymenoptera: Diapriidae). Annals of the Entomological Society of America, 105(4), 608–612.

Forbes A. A., Stelinski L. L., Powell T. H. Q., Smith J. J., Feder J. L. (2009). Sequential sympatric speciation across trophic levels. Science, 323, 776–779.

Fujiyama, N., H. Ueno, S. Kahono, S. Hartini, Matsubayashi, K. W., Kobayashi, N., & Katakura, H. (2013). Distribution and differentiation of *Henosepilachna diekei* (Coleoptera: Coccinellidae) on two host-plant species across Java, Indonesia. Ann Entomol Soc Am, 106, 741–752.

Fullaway, D.T. (1911). Monograph of the gall making Cynipidae (Cynipinae) of California. Ann. ent. Soc. Am. 4, 331–381.

Funk, D. J. & Omland, K. E. (2003). Species-level paraphyly and polyphyly: frequency, causes, and consequences, with insights from animal mitochondrial DNA. Annual Review of Ecology, Evolution, and Systematics, 34(1), 397–423.

Goulet H., Huber J. T. (eds). (1993). Hymenoptera of the world: An identification guide to families. Agriculture Canada, Canada.

Gillette, C. P. (1896). A monograph of the genus *Synergus*. Transactions of the American Entomological Society, 23, 85–100

Hamerlinck, G., Hulbert, D., Hood, G. R., Smith, J. J., & Forbes, A. A. (2016). Histories of host shifts and cospeciation among free-living parasitoids of *Rhagoletis* flies. Journal of Evolutionary Biology, 29, 1766–1779.

Hipp, A. L., Manos, P. S., González-Rodríguez, A., Hahn, M., Kaproth, M., McVay, J. D., … Cavender-Bares, J. (2018). Sympatric parallel diversification of major oak clades in the Americas and the origins of Mexican species diversity. New Phytologist, 217, 439–452.

Hood G. R., Forbes A. A., Powell T. H. Q., Egan S. P., Hamerlinck G., Smith J. J., & Feder J. L. (2015). Sequential Divergence and the Multiplicative Origin of Community Diversity. Proceedings of the National Academy of Sciences, U.S.A., 112, E5980–E5989.

Hood, G. R., Zhang, L., Hu, E. G., Ott, J. R., & Egan, S. P. (2019). Cascading reproductive isolation: Plant phenology drives temporal isolation among populations of a host-specific herbivore. Evolution, 73(3), 554–568.

Ide, T., Kusumi, J., Miura, K., & Abe, Y. (2018). Gall Inducers Arose from Inquilines: Phylogenetic Position of a Gall-Inducing Species and Its Relatives in the Inquiline Tribe Synergini (Hymenoptera: Cynipidae). Annals of the Entomological Society of America, 111(1), 6–12.

Joseph, M. B., Gentles, M., & Pearse, I. S. (2011). The parasitoid community of *Andricus quercuscalifornicus* and its association with gall size, phenology, and location. Biodiversity and Conservation, 20(1), 203–216.

Joyce, A.L., J. S. Bernal, S.B. Vinson, R.E. Hunt, F. Schulthess, & R.F. Medina. (2010). Geographic variation in male courtship acoustics and genetic divergence of populations of the *Cotesia flavipes* species complex. Entomol Exp Appl., 137, 153–164.

Kaiser, L., Le Ru, B. P., Kaoula, F., Paillusson, C., Capdevielle-Dulac, C., Obonyo, J. O., … & Silvain, J. F. (2015). Ongoing ecological speciation in *Cotesia sesamiae*, a biological control agent of cereal stem borers. Evolutionary applications, 8(8), 807–820.

Kester, K.M., Eldei G.M., & Brown, B.L. (2015). Genetic differentiation of two host–foodplant complex sources of *Cotesia congregata* (Hymenoptera: Braconidae). Ann Entomol Soc Am., 108, 1014–2015.

König, K., Krimmer, E., Brose, S., Gantert, C., Buschlüter, I., König, C., … Steidle, J. L. (2015). Does early learning drive ecological divergence during speciation processes in parasitoid wasps? Proceedings of the Royal Society B: Biological Sciences, 282(1799), 20141850.

Lewis, O. T., Memmott, J., Lasalle, J., Lyal, C. H., Whitefoord, C., & Godfray, H. C. J. (2002). Structure of a diverse tropical forest insect–parasitoid community. Journal of Animal Ecology, 71(5), 855–873.

Lobato-Vila, I., Cibrián-Tovar, D., Barrera-Ruíz, U. M., Equihua-Martínez, A., Estrada-Venegas, E. G., Buffington, M. L., & Pujade-Villar, J. (2019). Review of the Synergus Hartig Species (Hymenoptera: Cynipidae: Synergini) Associated with Tuberous and Other Tumor-Like Galls on Oaks from the New World with the Description of Three New Species from Mexico. Proceedings of the Entomological Society of Washington, 121(2), 193.

Lobato-Vila, I., Cibrián-Tovar, D., BarreraRuíz, U. M., & Pujade-Villar, J. (2018). Study of the inquiline oak gall wasp fauna emerged from agamic galls of *Andricus quercuslaurinus* Melika and Pujade-Villar, 2009 from Mexico. Southwestern Entomologist 43(3), 591–610.

Lobato-Vila, I. & Pujade-Villar, J. (2017). Description of five new species of inquiline oak gall wasps of the genus Synergus Hartig (Hymenoptera, Cynipidae: Synergini) with partially smooth mesopleurae from Mexico. Zoological Studies, 56(36), 1–28

Mayr, G. (1872). Die Einmiethler der mitteleuropäischen Eichengallen. Verh. zool. bot. Ges. Wien, 22, 669–726.

Melika, G. (2006). Gall wasps of Ukraine. Cynipidae. Vestnik Zoologii 21 (Supplement 1), 1–300.

McCracken, M. I., & Egbert, D. B. (1922). California Gall-making Cynipidae: With Descriptions of New Species (Vol. 3, No. 1–4). Stanford University Press.

Moriya, S., Inoue, K., Otake, A., Shiga, M., & Mabuchi, M. (1989). Decline of the Chestnut Gall Wasp Population, *Dryocosmus kuriphilus* YASUMATSU (Hymenoptera : Cynipidae) after the Establishment of *Torymus sinensis* KAMIJO (Hymenoptera : Torymidae). Applied Entomology and Zoology, 24(2), 231–233.

Muesebeck, C. F. W., Krombein, K. V., & Townes, H. K. (1951). Hymenoptera of America north of Mexico. US Government Printing Office.

Nazemi, J., Talebi, A. A., Sadeghi, S. E., Melika, G. & Lozan, A. (2008). Species richness of oak gall wasps (Hymenoptera: Cynipidae) and identification of associated inquilines and parasitoids on two oak species in western Iran. North-Western Journal of Zoology, 4(2).

Nosil, P., & Flaxman, S. M. (2011). Conditions for mutation-order speciation. Proceedings of the Royal Society B: Biological Sciences, 278(1704), 399–407.

Nosil, P., & Hohenlohe, P. A. (2012). Dimensionality of sexual isolation during reinforcement and ecological speciation in *Timema cristinae* stick insects. Evolutionary Ecology Research, 14(4), 467–485.

Noyes, J. (2012). An inordinate fondness of beetles, but seemingly even more fond of microhymenoptera! Hamuli 3: 5–8.

Pénzes, Z., Tang, C. T., Bihari, P., Bozsó, M., Schwéger, S., & Melika, G. (2012). Oak associated inquilines (Hymenoptera, Cynipidae, Synergini). TISCIA Monograph series 11, 1–76.

Pénzes, Z., C.-T. Tang, G. N. Stone, J. A. Nicholls, S. Schwéger, M. Bozsó, and G. Melika. 2018. Current status of the oak gallwasp (Hymenoptera: Cynipidae: Cynipini) fauna of the Eastern Palaearctic and Oriental Regions. Zootaxa. 4433: 245–289.

Price, P. W. (1980). Evolutionary Biology of Parasites. Princeton University Press, Princeton, NJ, USA.

Prior, K. M., & Hellmann, J. J. (2010). Impact of an invasive oak gall wasp on a native butterfly: a test of plant-mediated competition. Ecology, 91(11), 3284–3293.

Puillandre, N., Lambert, A., Brouillet, S., & Achaz, G. (2011). ABGD, Automatic Barcode Gap Discovery for primary species delimitation. Molecular Ecology, 21(8), 1864–1877.

Pujade-Villar, J., Melika, G., Ros-Farré, P., Acs, Z., & Csóka, G. (2003). Cynipid inquiline wasps of Hungary, with taxonomic notes on the western Palaearctic fauna (Hymenoptera: Cynipidae, Cynipinae, Synergini). Folia Entomologica Hungarica, 64, 121–170.

Rhodes, B. C., Blair, C. P., Takahashi, M. K., & Abrahamson, W. G. (2012). The role of olfactory cues in the sequential radiation of a gall-boring beetle, *Mordellistena convicta*. Ecological Entomology, 37(6), 500–507.

Rodríguez, M. Á., & Hawkins, B. A. (2000). Diversity, function and stability in parasitoid communities. Ecology letters, 3(1), 35–40.

Roininen, H., Price, P. W. and Tahvanainen, J. (1996). Bottom-up and top-down influences in the trophic system of a willow, a galling sawfly, parasitoids and inquilines. Oikos, 77, 44–50.

Ronquist, F., Teslenko, M., van der Mark, P., Ayres, D. L., Darling, A., Höhna, S., … Huelsenbeck, J. P. (2012). MrBayes 3.2: efficient Bayesian phylogenetic inference and model choice across a large model space. Systematic biology, 61(3), 539–542. doi:10.1093/sysbio/sys029

Roskam, J. C. (1979). Biosystematics of insects living in female birch catkins. II. Inquiline and predaceous gall midges belonging to various genera. Netherlands Journal of Zoology, 29(3), 283–351.

Sanver, D. & Hawkins, B. A. (2000). Galls as habitats: the inquiline communities of insect galls. Basic and Applied Ecology, 1(1), 3–11.

Schönrogge, K., Stone, G. N. & Crawley, M. J. (1996). Alien herbivores and native parasitoids: rapid developments and structure of the parasitoid and inquiline complex in an invading gall wasp *Andricus quercuscalicis* (Hymenoptera: Cynipidae). Ecological Entomology, 21(1), 71–80.

Schönrogge, K., Walker, P. & Crawley, M. J. (2000). Parasitoid and inquiline attack in the galls of four alien, cynipid gall wasps: host switches and the effect on parasitoid sex ratios. Ecological Entomology, 25(2), 208–219.

Smith, M. A., Woodley, N. E., Janzen, D. H., Hallwachs, W., & Hebert, P. D. (2006). DNA barcodes reveal cryptic host-specificity within the presumed polyphagous members of a genus of parasitoid flies (Diptera: Tachinidae). Proceedings of the National Academy of Sciences, 103(10), 3657–3662.

Smith, M. A., Rodriguez, J. J., Whitfield, J. B., Deans, A. R., Janzen, D. H., Hallwachs, W., & Hebert, P. D. (2008). Extreme diversity of tropical parasitoid wasps exposed by iterative integration of natural history, DNA barcoding, morphology, and collections. Proceedings of the National Academy of Sciences, 105(34), 12359–12364.

Smith, M. A., Eveleigh, E. S., McCann, K. S., Merilo, M. T., McCarthy, P. C., & Van Rooyen, K. I. (2011). Barcoding a quantified food web: crypsis, concepts, ecology and hypotheses. PLoS One, 6(7), e14424.

Stamatakis, A. (2014). RAxML version 8: a tool for phylogenetic analysis and post-analysis of large phylogenies. Bioinformatics, 30(9), 1312–1313.

Stireman, J. O., Nason, J. D., Heard, S. B., & Seehawer, J. M. (2005). Cascading host-associated genetic differentiation in parasitoids of phytophagous insects. Proc Biol Sci, 273(1586), 523–530.

Stone, G. N., Schönrogge, K., Atkinson, R. J., Bellido, D., Pujade-Villar, J. (2002). The population biology of oak gall wasps (Hymenoptera: Cynipidae). Annu. Rev. Entomol., 47, 633–668.

Stone, G. N., Schönrogge, K., Crawley, M. J., & Fraser, S. (1995). Geographic and between-generation variation in the parasitoid communities associated with an invading gallwasp, *Andricus quercuscalicis* (Hymenoptera: Cynipidae). Oecologia, 104(2), 207–217.

Stone, G. N., White, S. C., Csóka, G., Melika, G., Mutun, S., Pénzes, Z., … Nicholls, J. A. (2017). Tournament ABC analysis of the western Palaearctic population history of an oak gall wasp, *Synergus umbraculus*. Molecular Ecology, 26(23), 6685–6703.

Wahl D.B. Genera Ichneumonorum Nearcticae. 2015 (and updates). Version: 2015.08.17. http://www.amentinst.org/GIN/.

Walker, P., Leather, S. R., & Crawley, M. J. (2002). Differential rates of invasion in three related alien oak gall wasps (Cynipidae: Hymenoptera). Diversity and Distributions, 8(6), 335–349.

Ward, A. K. G., Khodor, O. S., Egan, S. P., Weinersmith, K. L., & Forbes, A. A. (2019). A keeper of many crypts: a behaviour-manipulating parasite attacks a taxonomically diverse array of oak gall wasp species. Biology Letters, 15(9), 20190428.

Weld L. H. 1959 Cynipid galls of the Eastern United States. Ann Arbor, MI: Privately printed.

Wiebes-Rijks, A. A. (1979). A character analysis of the species *Synergus* Hartig, section II (Mayr, 1872) (Hymenoptera, Cynipidae). Zoologische Mededelingen, 53(28), 297–321

Weinersmith KL, Forbes AA, Ward AKG, Brandão-Dias PFP, Zhang YM, Egan SP. 2020. Arthropod Community Associated with the Asexual Generation of *Bassettia pallida* Ashmead (Hymenoptera: Cynipidae). Annals of the Entomological Society of America. In Press.

Zhang, J., Kapli, P., Pavlidis, P., & Stamatakis, A. (2013). A general species delimitation method with applications to phylogenetic placements. Bioinformatics, 29(22), 2869–2876.

