## Supplementary Figures 1-46 for "Diversity, host ranges, and potential drivers of speciation among the inquiline enemies of oak gall wasps"

Figure S1. *Synergus laeviventris* (clade 1) – 1081-006-003 - female
from *Andricus flavohirtus* on *Quercus alba*, Cape Giradeau, MO

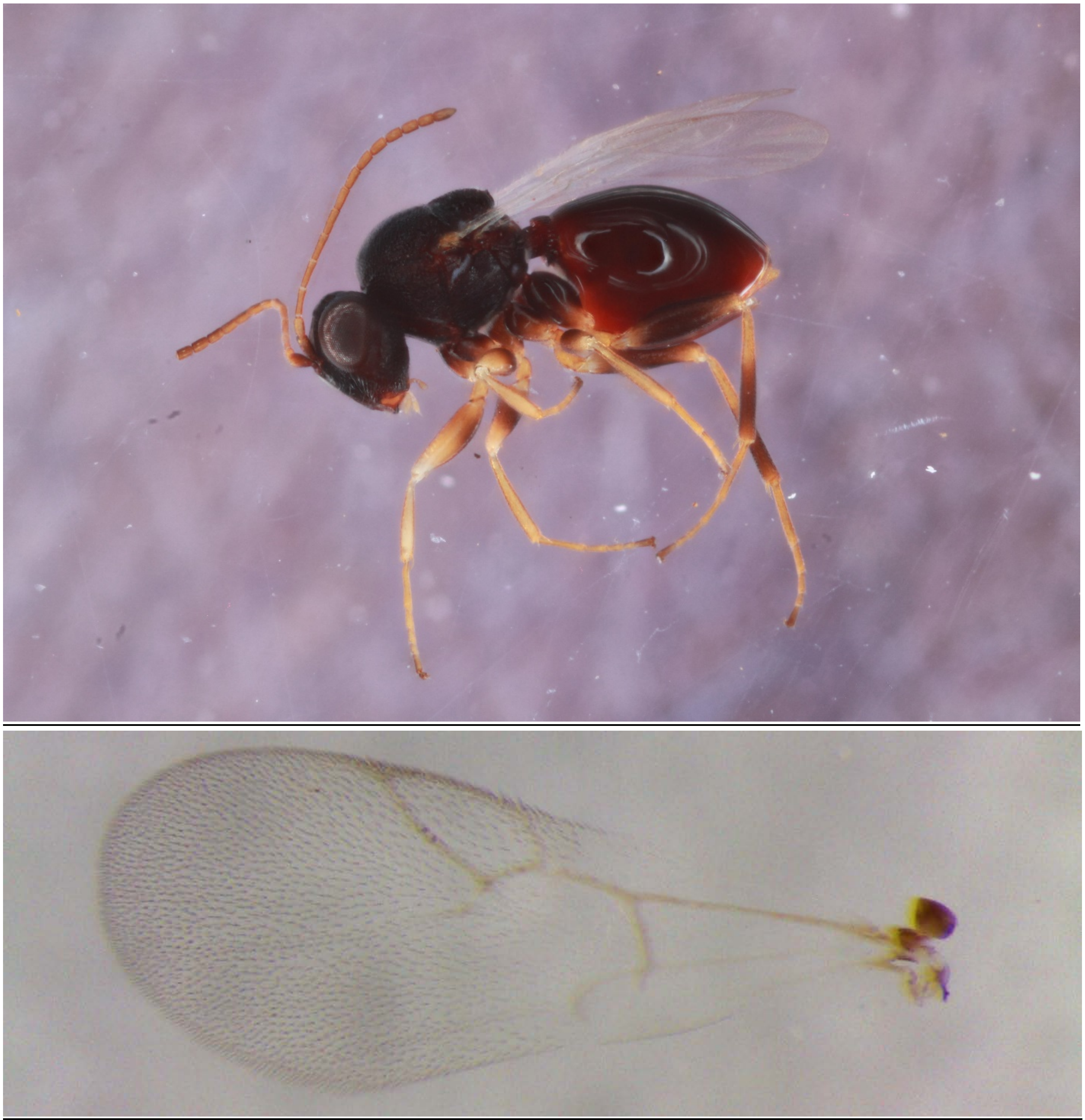

Figure S2. *Synergus laeviventris* (clade 1) – 1081-006-004C - male
from *Andricus flavohirtus* on *Quercus alba*, Cape Giradeau, MO

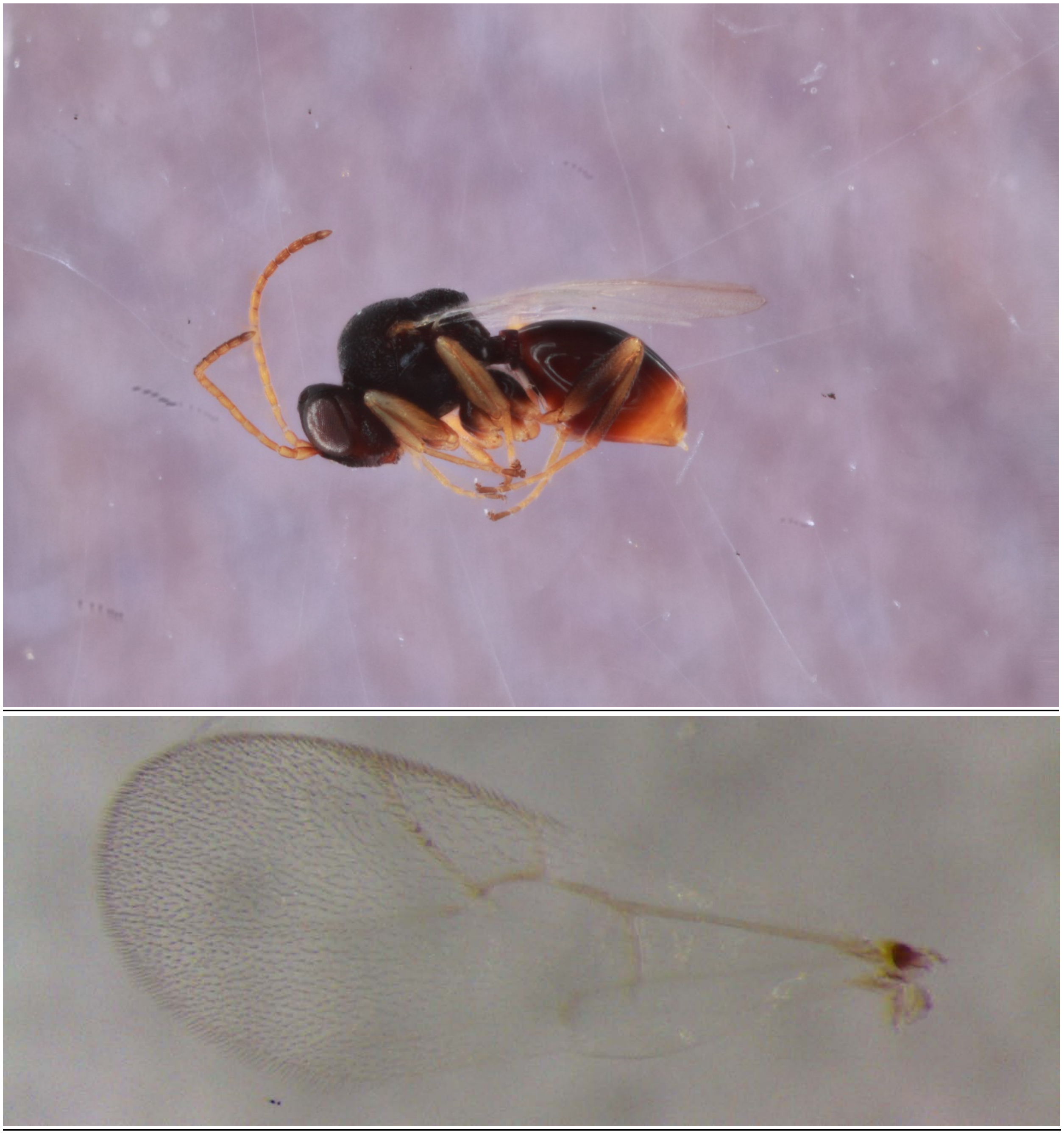

Figure S3. *Synergus laeviventris* (clade 2) – 658-019-002 - female from *Disholcaspis quercusglobulus* on *Quercus alba*, Tiffin, IA

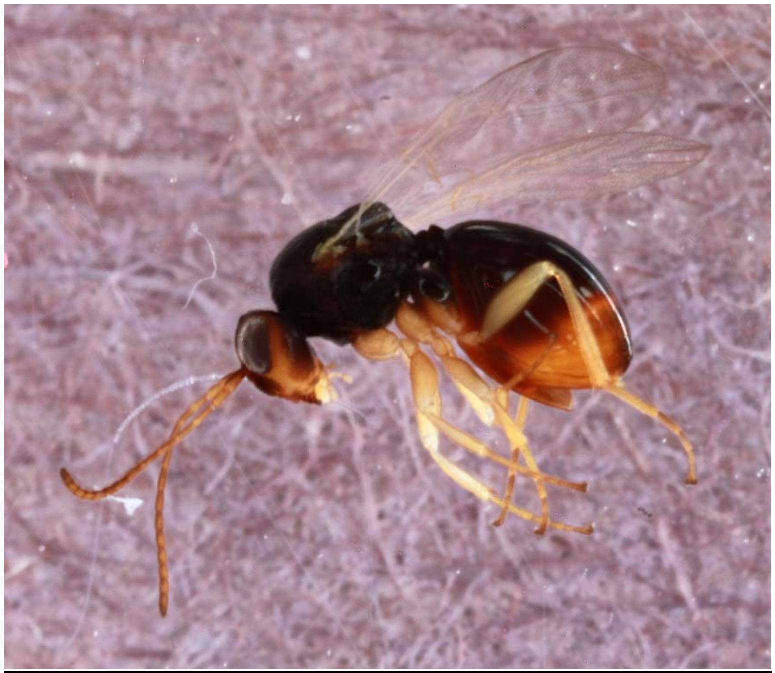

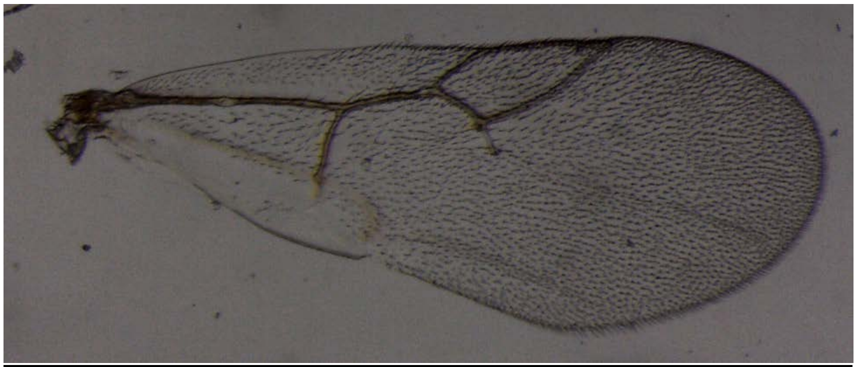

Figure S4. *Synergus laeviventris* (clade 2) – 806-019-11A - male
from *Disholcaspis quercusglobulus* on *Quercus alba*, Tiffin, IA

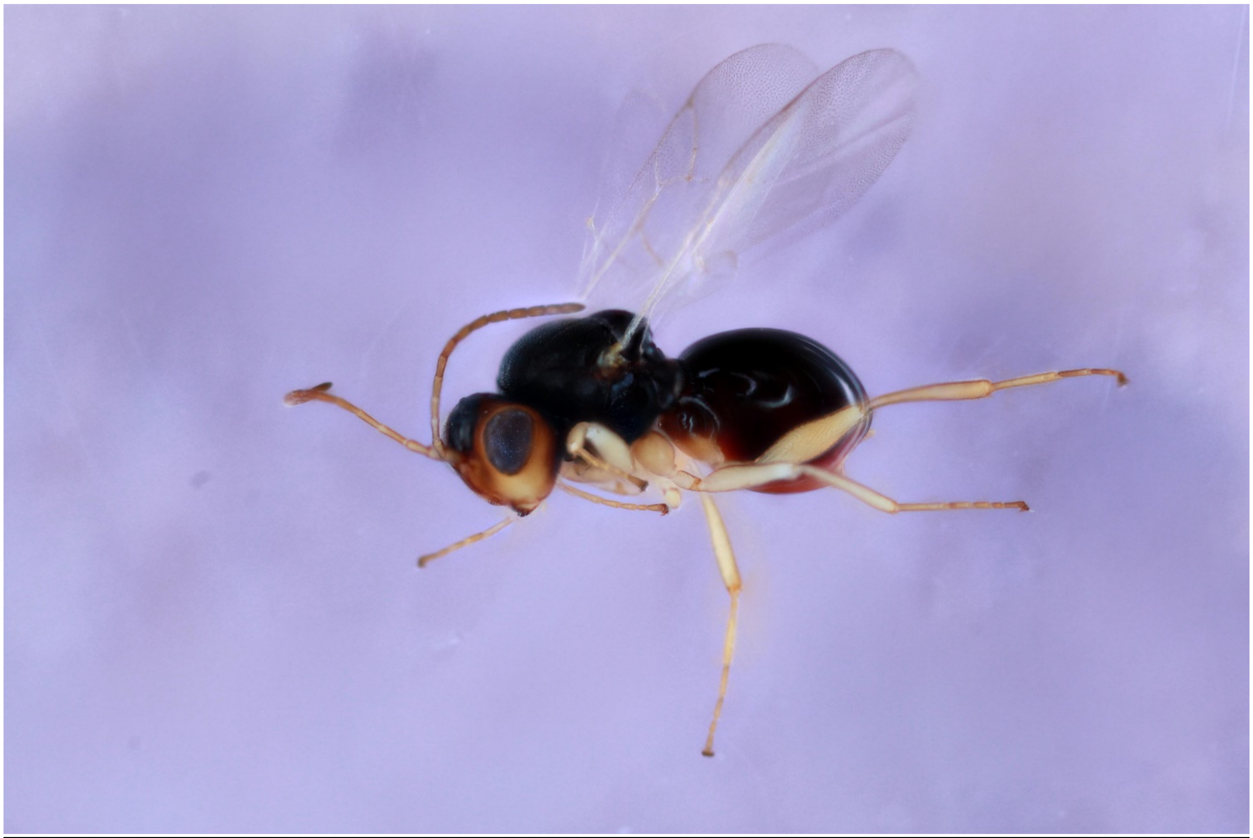

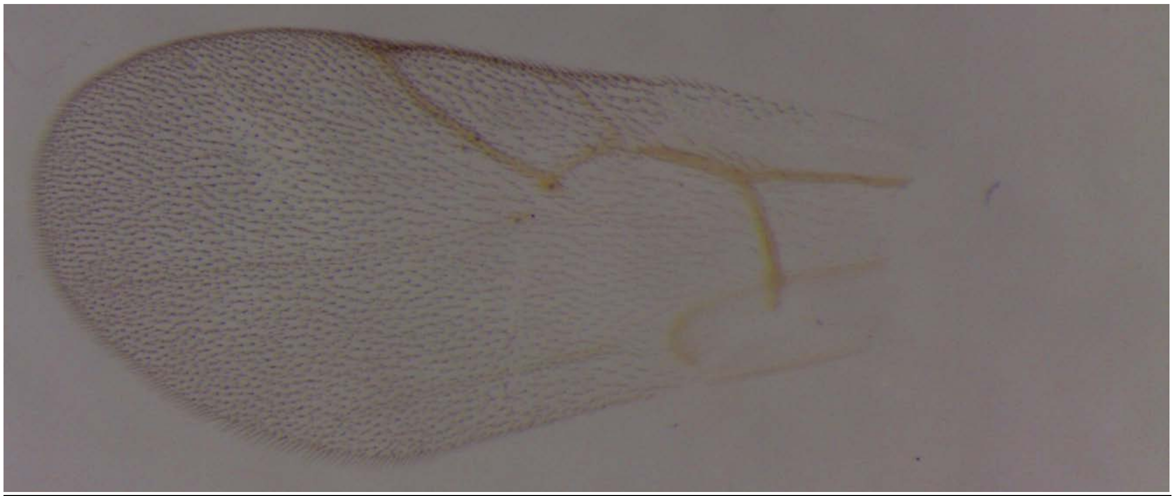

Figure S5. *Synergus laeviventris* (clade 3) – 791-045-6A - female
from *Andricus quercusostensackenii* on *Quercus palustris*, Iowa City, IA

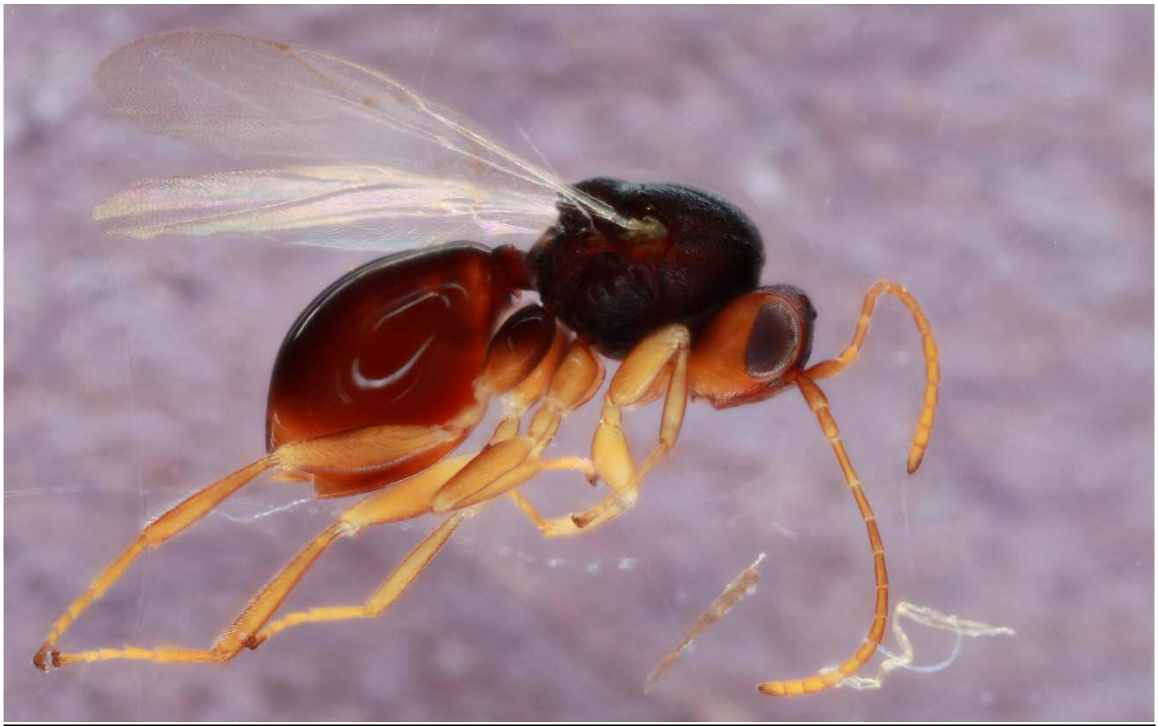

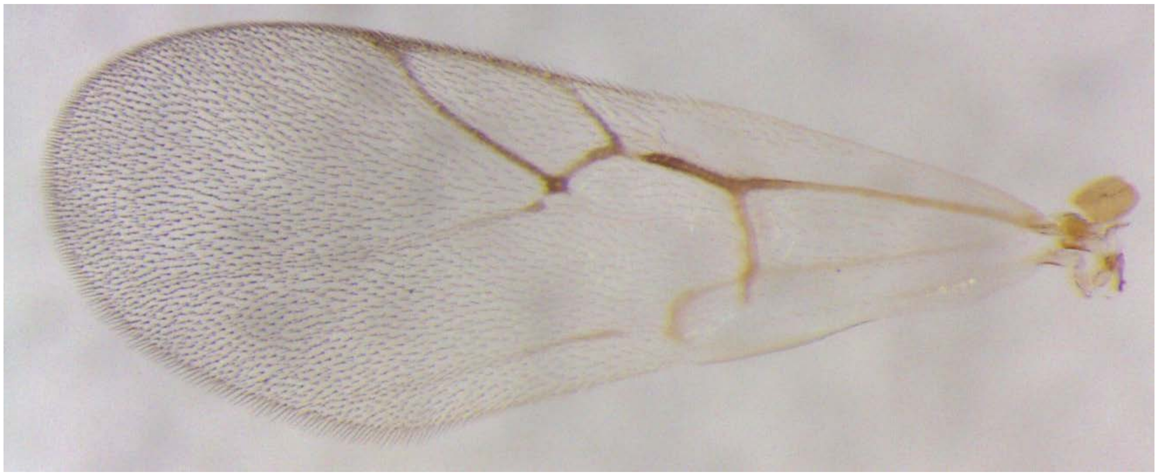

Figure S6. *Synergus laeviventris* (clade 3) – 814-086-005 - male
from *Amphibolips quercusinanis* on *Quercus palustris*, Tiffin, IA

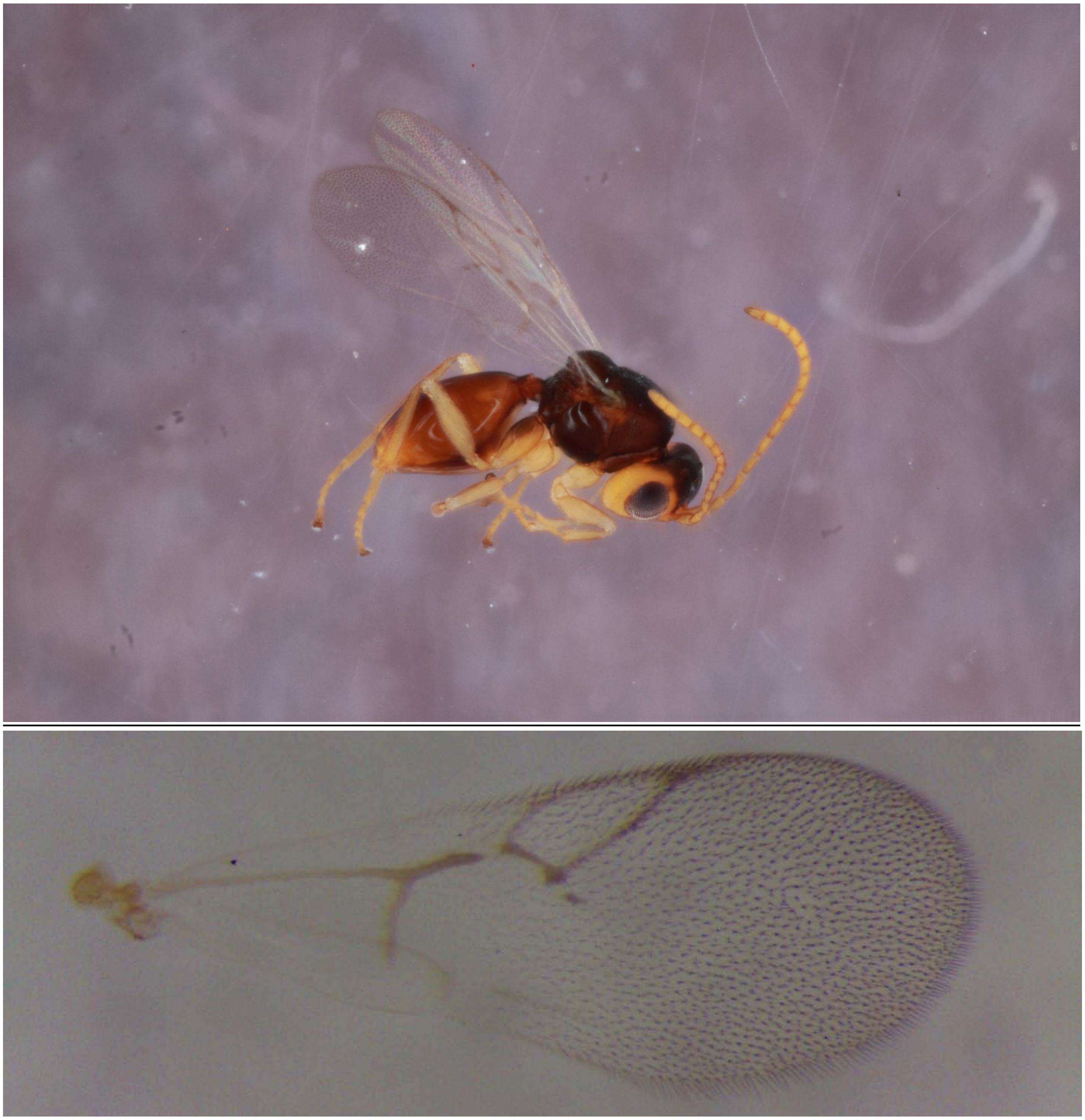

Figure S7. Synergus unk #1 (clade 4) – 698-999-10C - female
Host: unidentified stem gall on *Quercus macrocarpa*, Iowa City, IA

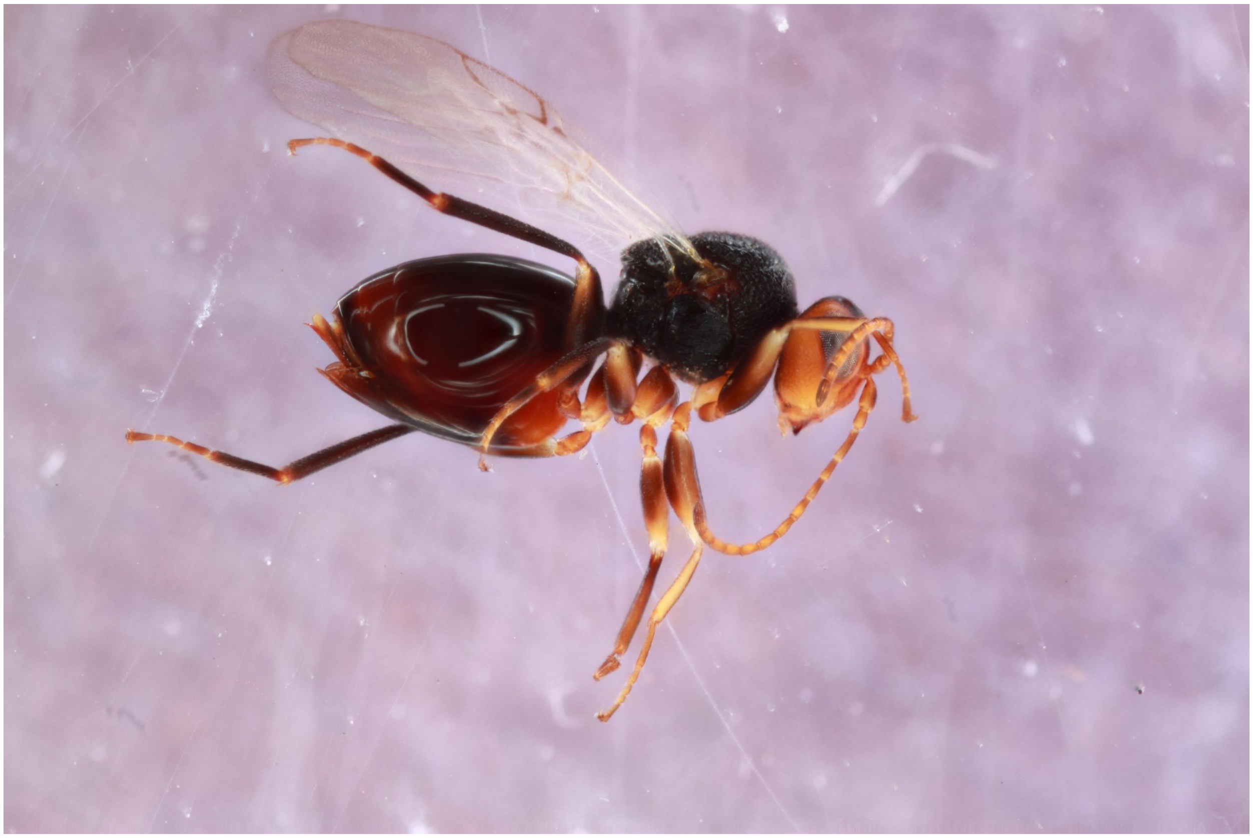

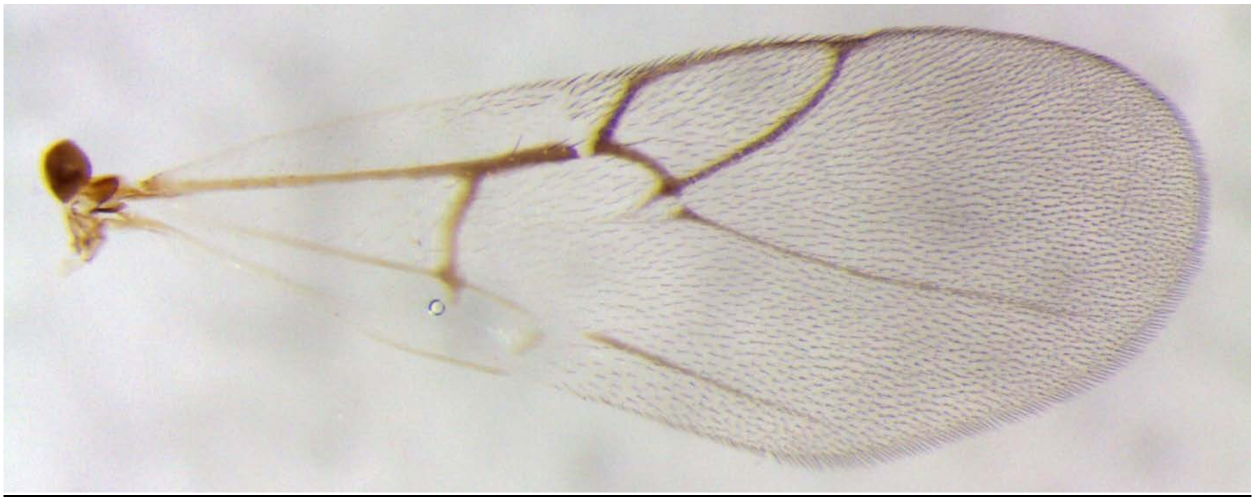

<u>Figure S8. Synergus unk #1 (clade 4) – 698-999-5A - male</u>
Host: unidentified stem gall on *Quercus macrocarpa*, Iowa City, IA

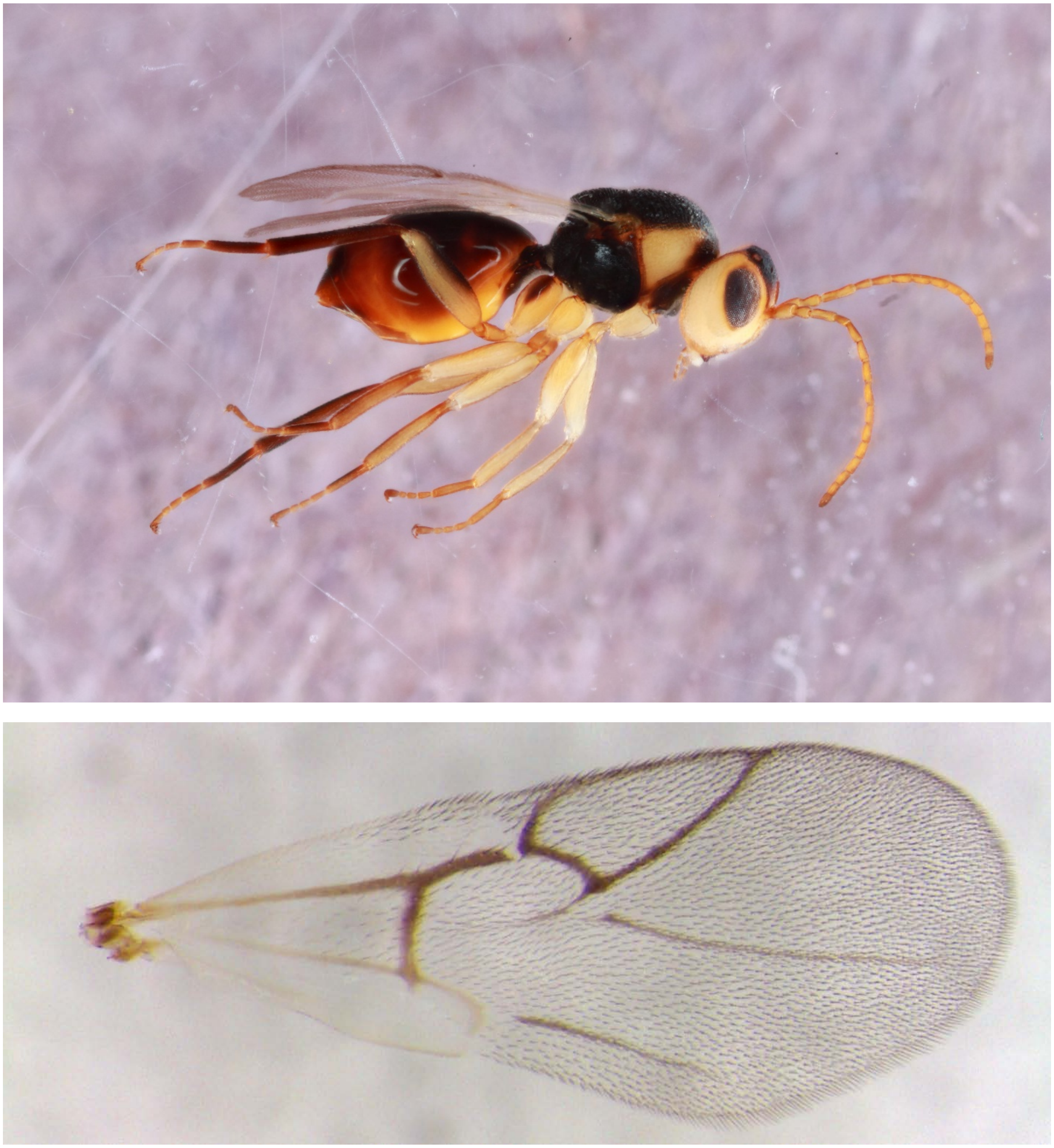

Figure S9. *Synergus coniferae* (Clade 5) – 293-041-4A – female
from *Callirhytis ventricosa* on *Quercus imbricaria*, Iowa City, IA

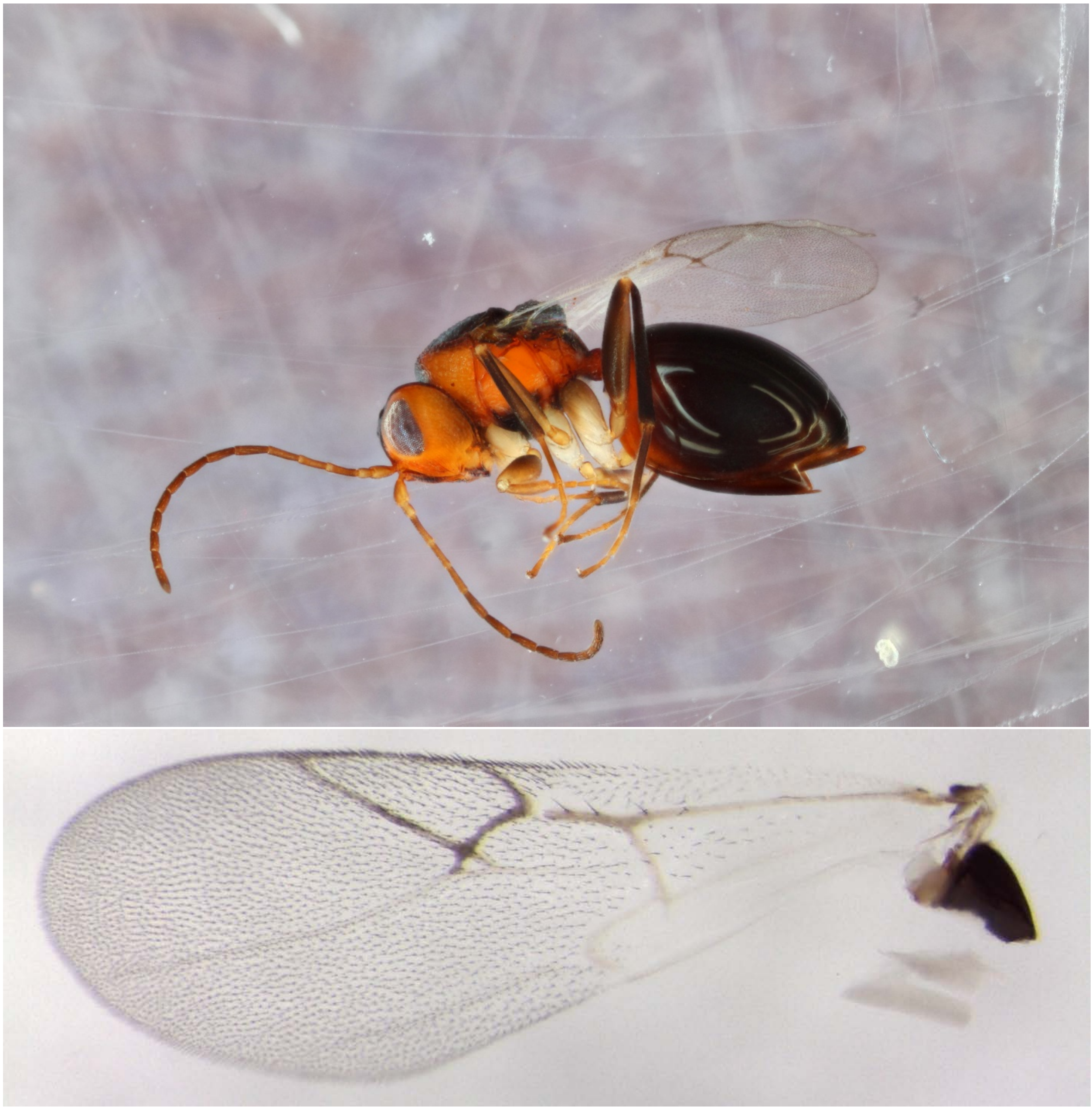

Figure S10. *Synergus coniferae* (Clade 5) – 293-041-4B – male
from *Callirhytis ventricosa* on *Quercus imbricaria*, Iowa City, IA (no wing picture)

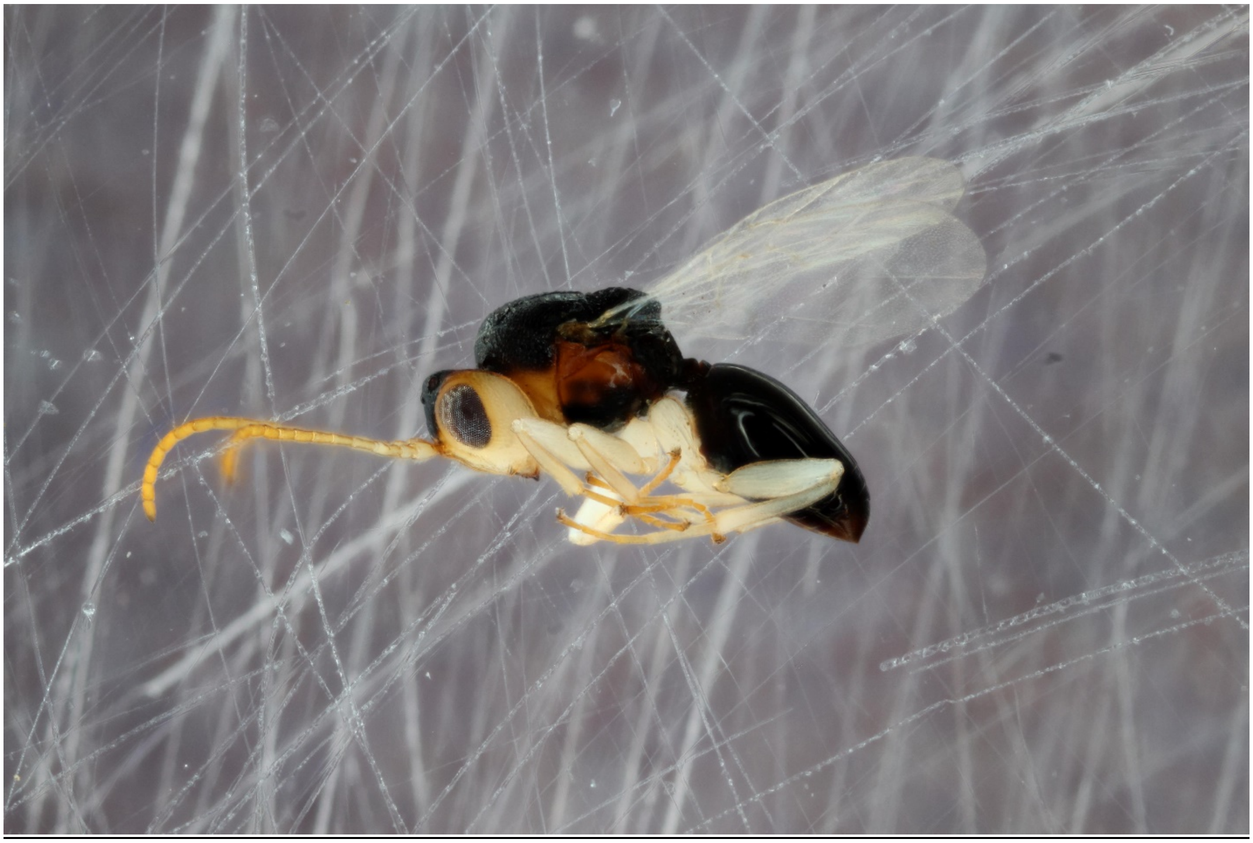

Figure S11. *Synergus lignicola* (= *Synergus davisi*) (Clade 6) – 1076-101-7A – female from *Callirhytis quercusgemmaria* on *Quercus rubra*, Traverse City, MI

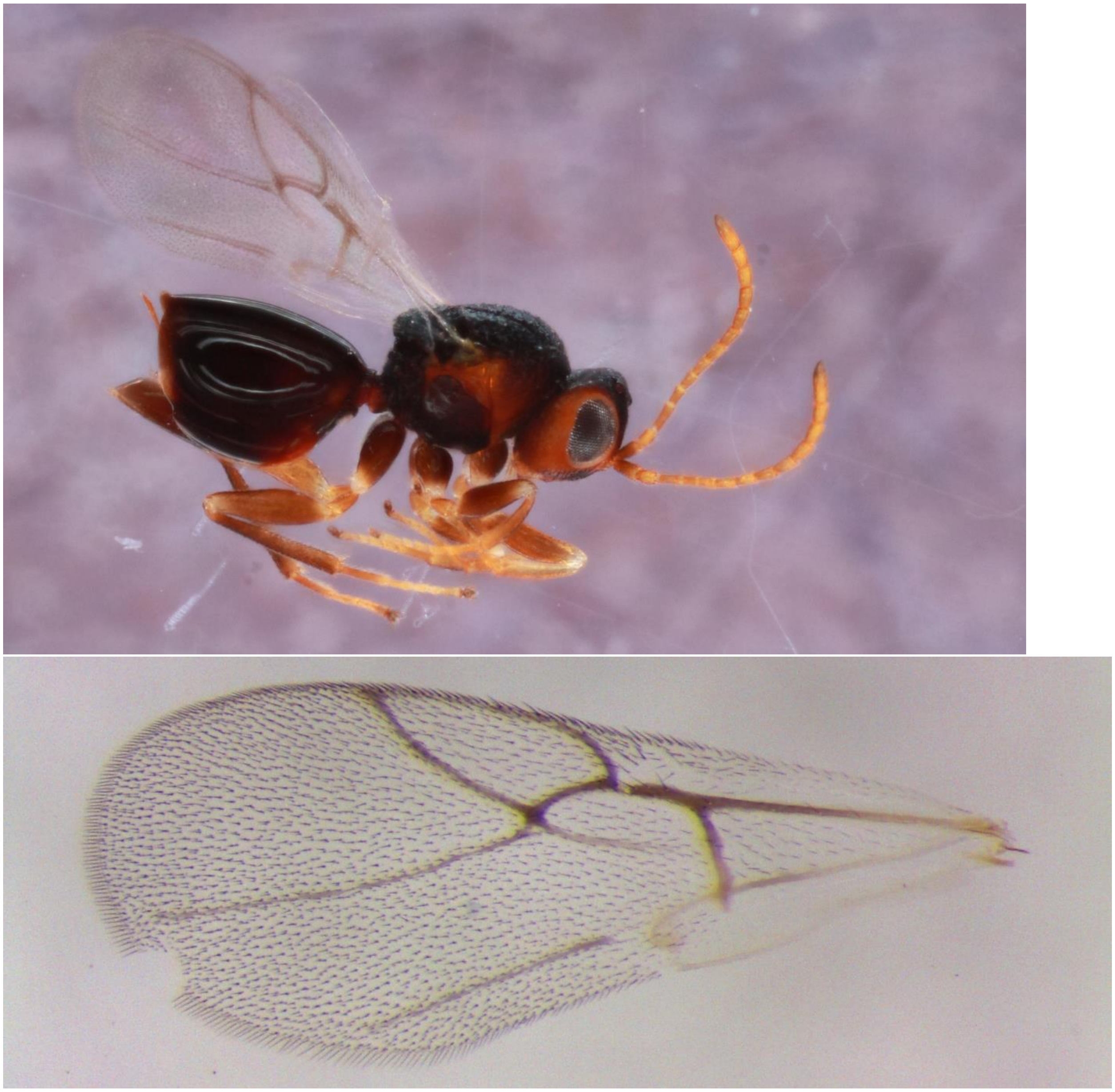

Figure S12. *Synergus lignicola* (= *Synergus davisi*) (Clade 6) – 1076-101-7A – male from *Callirhytis quercusgemmaria* on *Quercus rubra*, Traverse City, MI

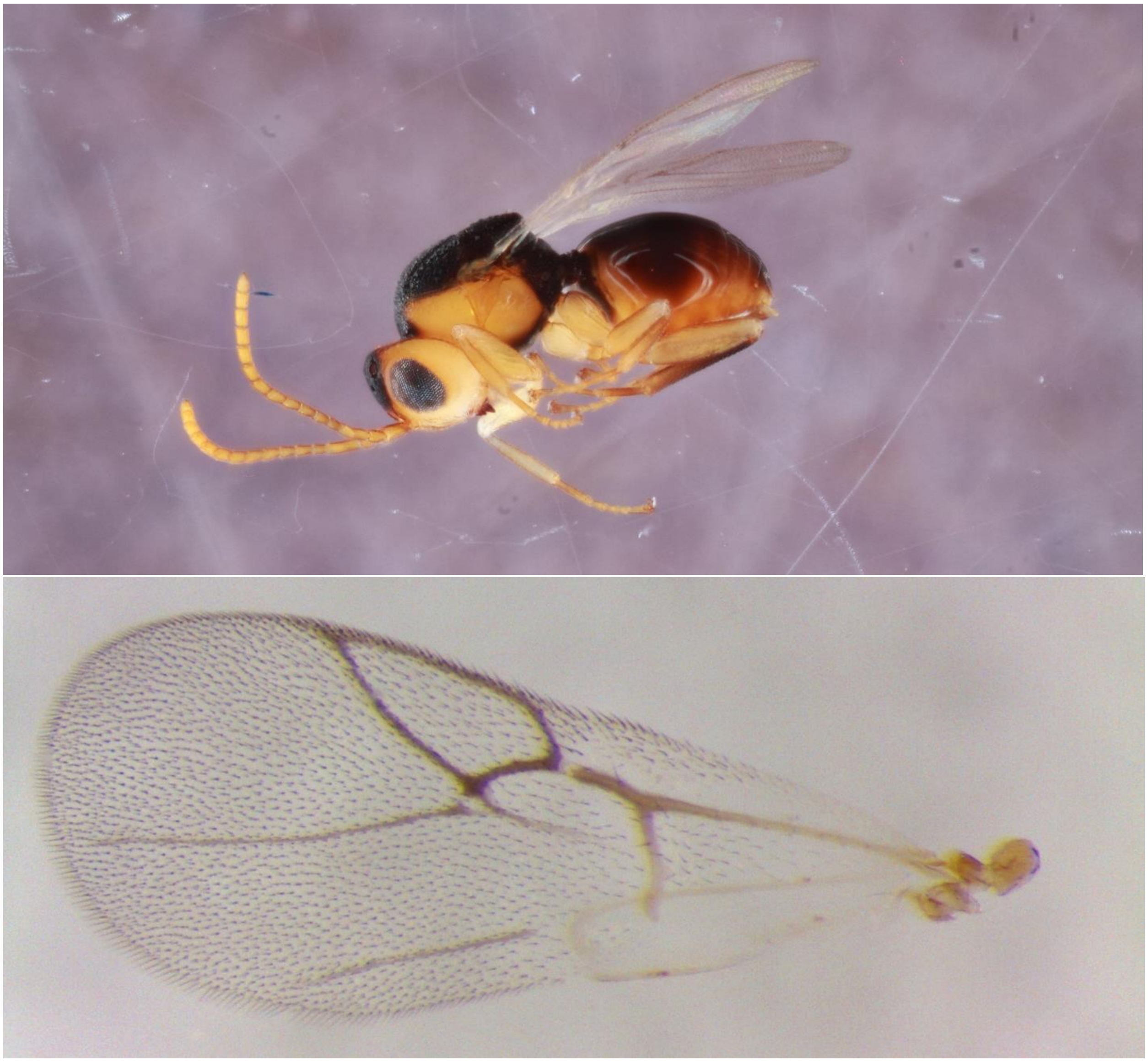

Figure S13. *Synergus* nr. *lignicola* (Clade 7) – 614-069-004A – female from *Callirhytis quercuspunctata* on *Quercus palustris*, St. Peters, MO

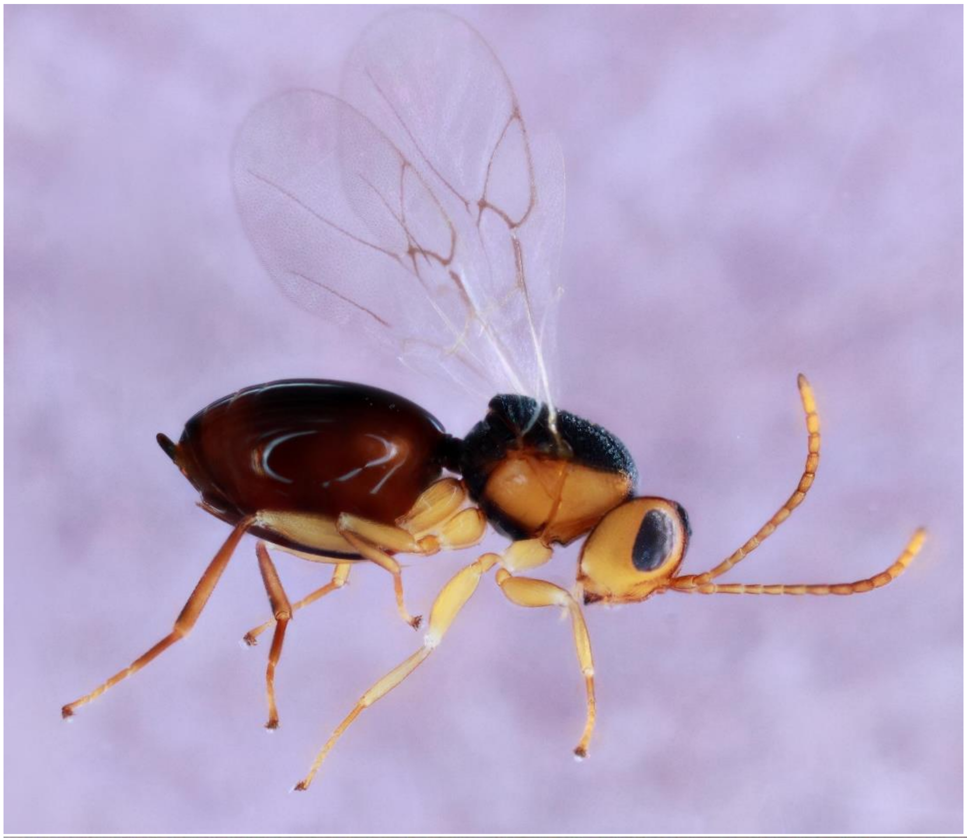

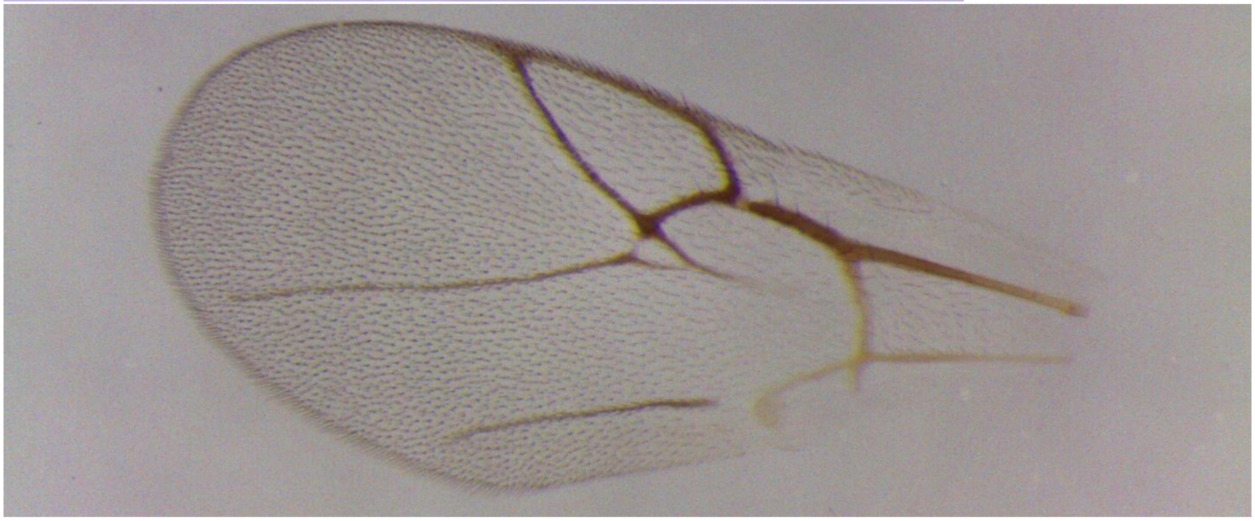

Figure S14. *Synergus* nr. *lignicola* (Clade 7) – 670-069-001 – male from *Callirhytis quercuspunctata* on *Quercus palustris*, St. Louis, MO

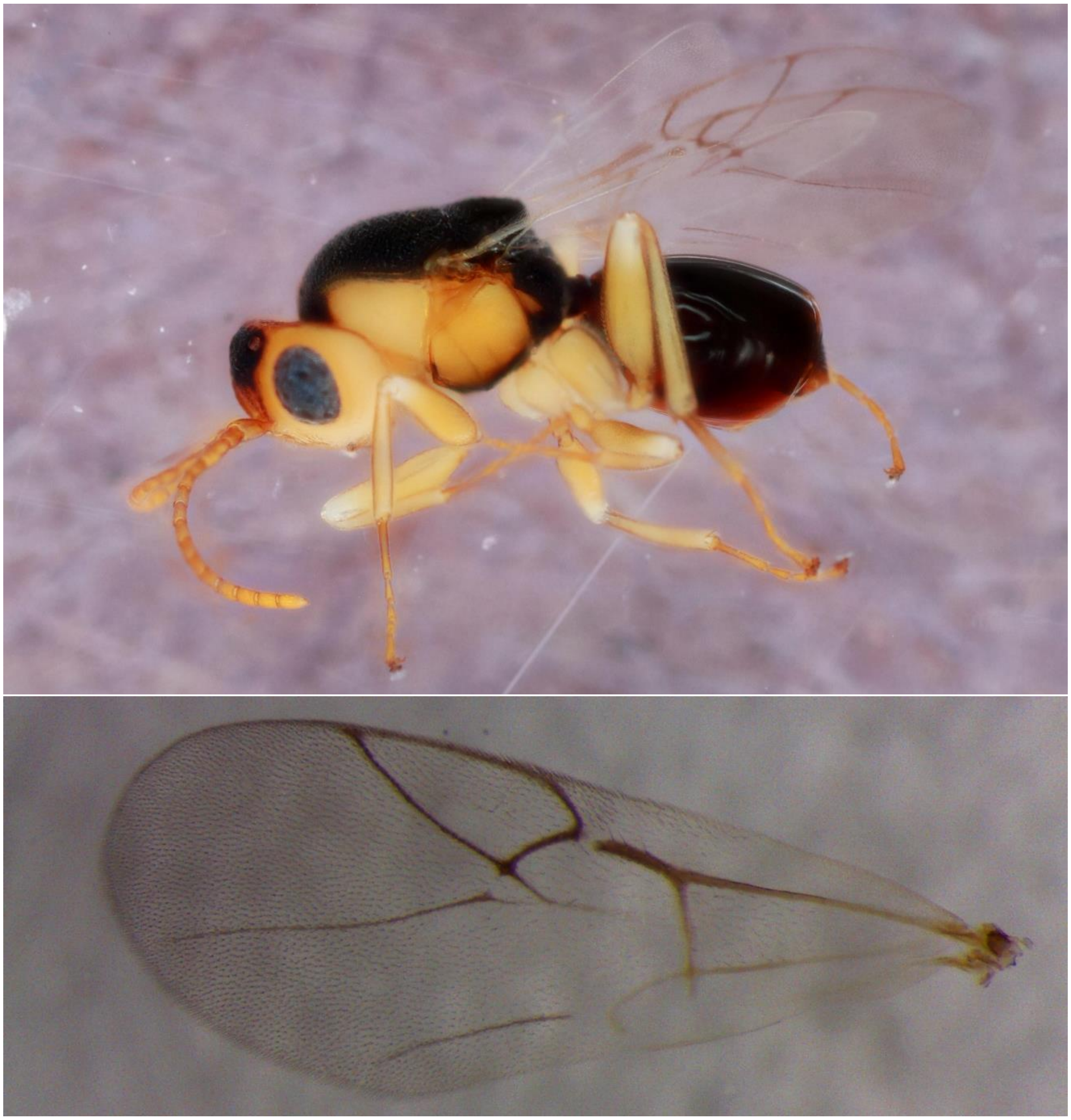

Figure S15. *Synergus oneratus* (Clade 8A) – 442-002-16B – female from *Acraspis erinacei* on *Quercus alba*, Tiffin, IA

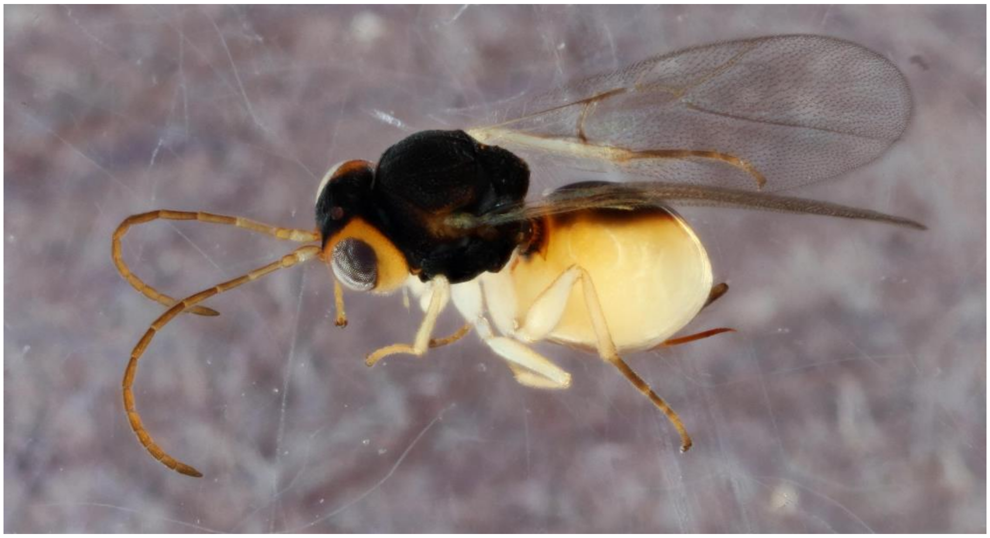

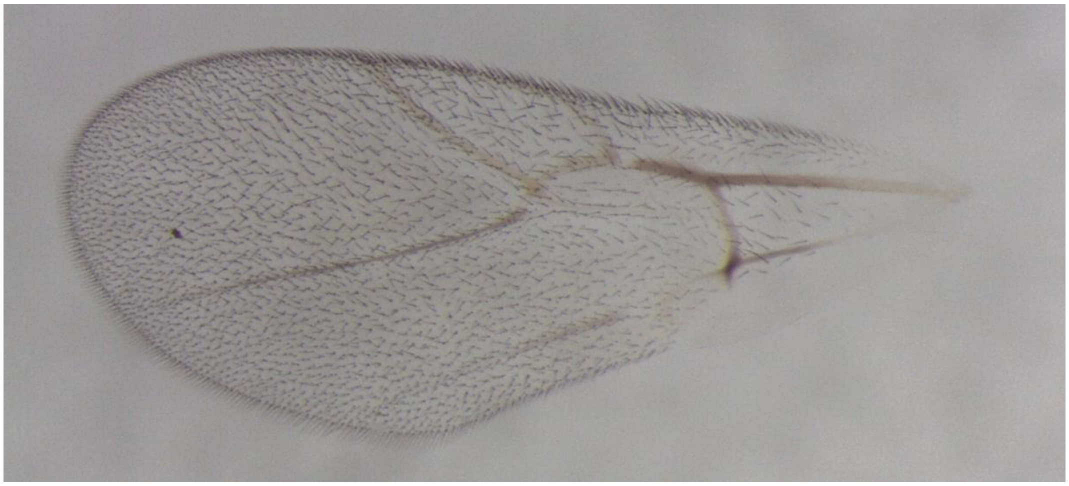

Figure S16. *Synergus oneratus* (Clade 8A) – 1105-003-005A – male (no wing picture) from *Acraspis pezomachoides* on *Quercus alba*, Paducah, KY

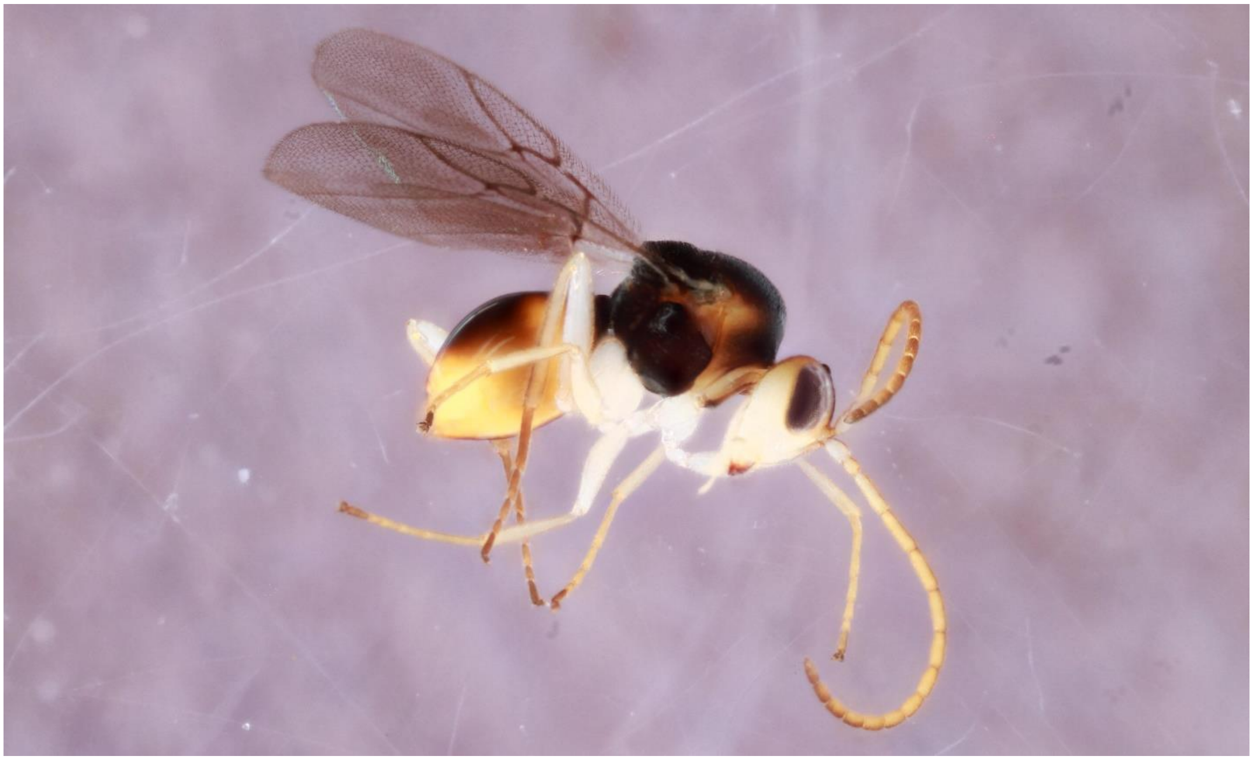

Figure S17. *Synergus oneratus* (Clade 8B) – 1201-003-002 – female
from *Acraspis pezomachoides* on *Quercus alba*, Columbus, OH

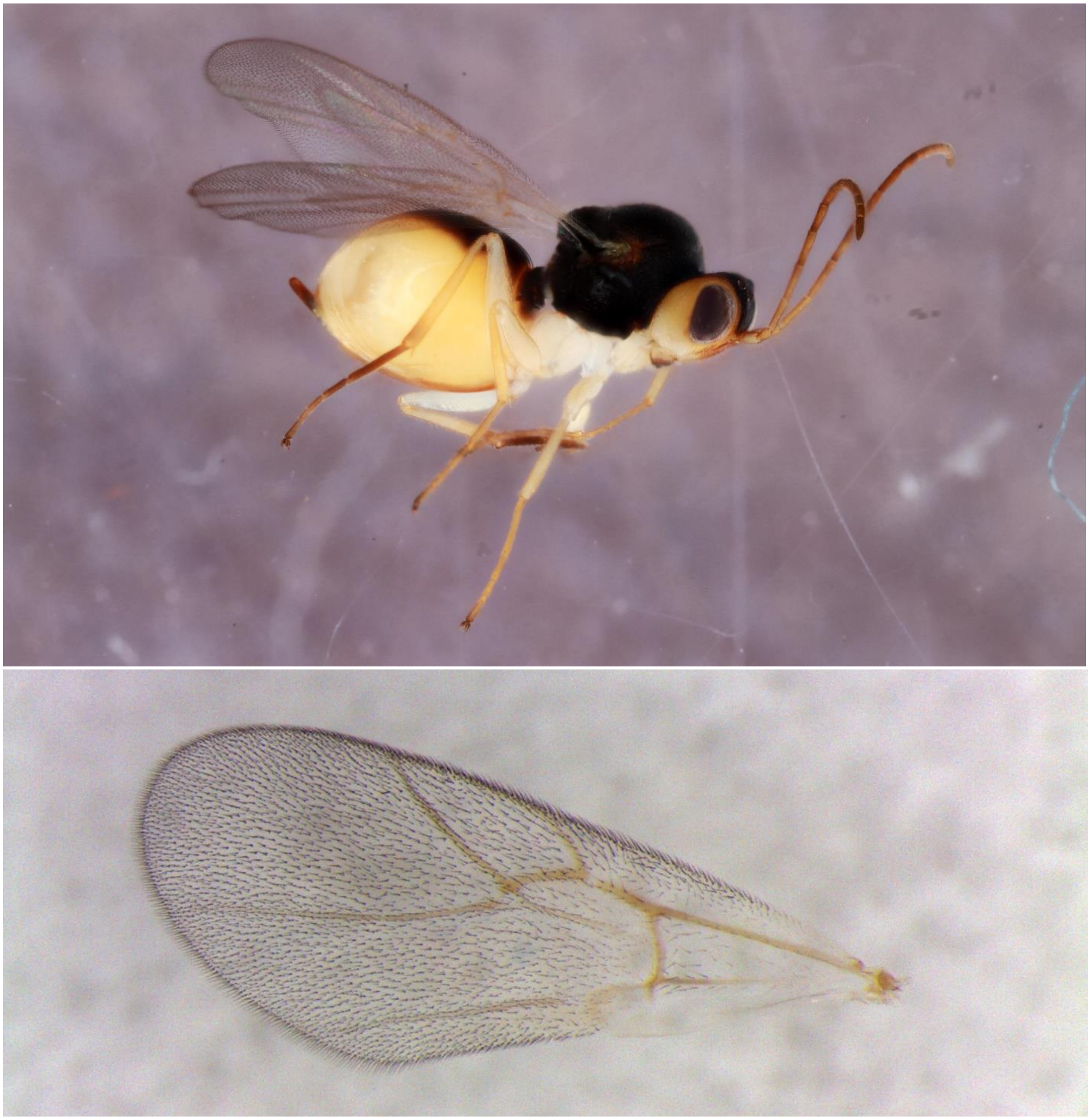

Figure S18. *Synergus oneratus* (Clade 9) – 539-067-004 – male
from *Andricus robustus* on *Quercus stellata*, St. Louis, MO

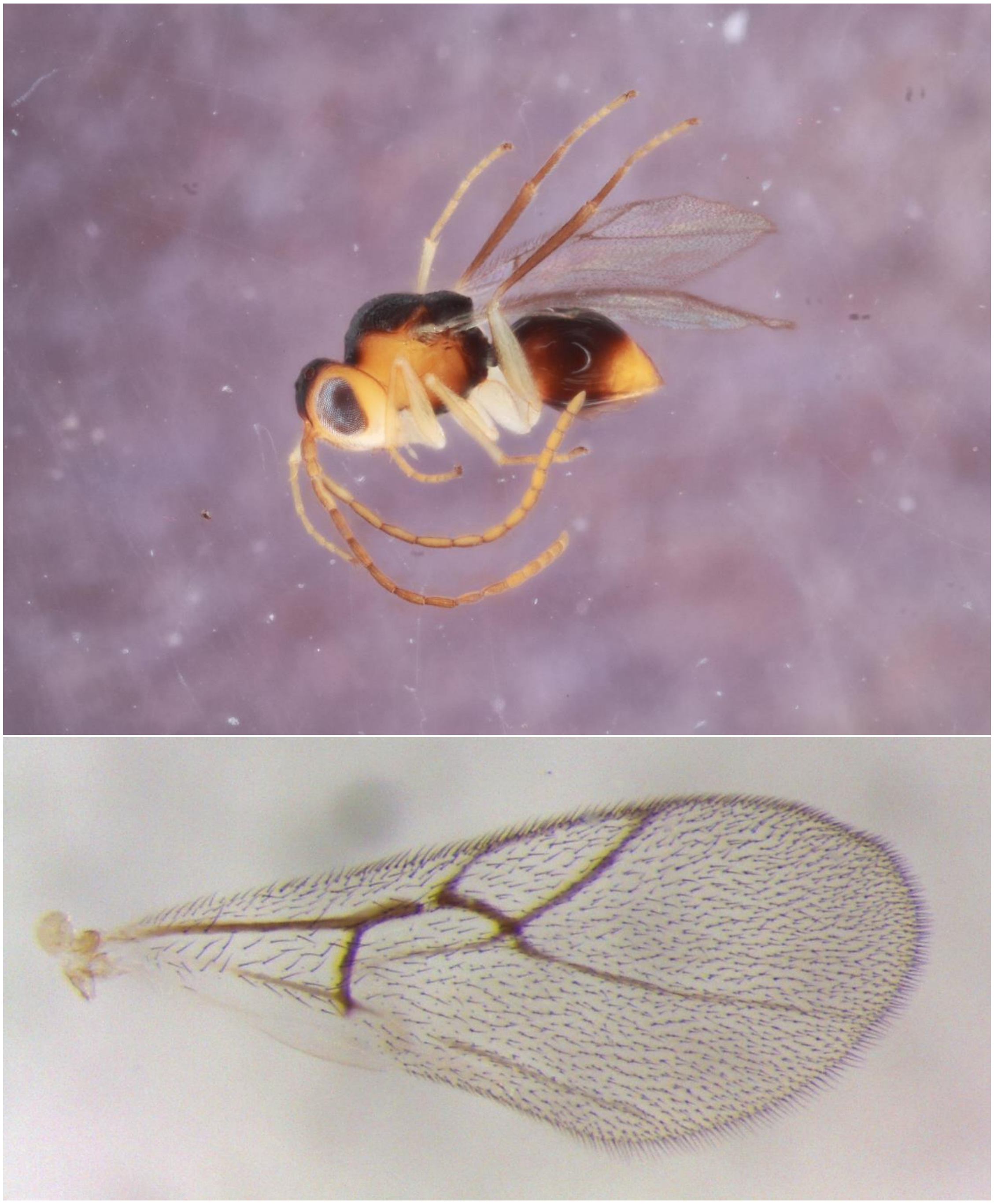

Figure S19. *Synergus oneratus* (Clade 10) – 500-011-006A – female from *Andricus quercusstrobilanus* on *Quercus bicolor*, Iowa City, IA

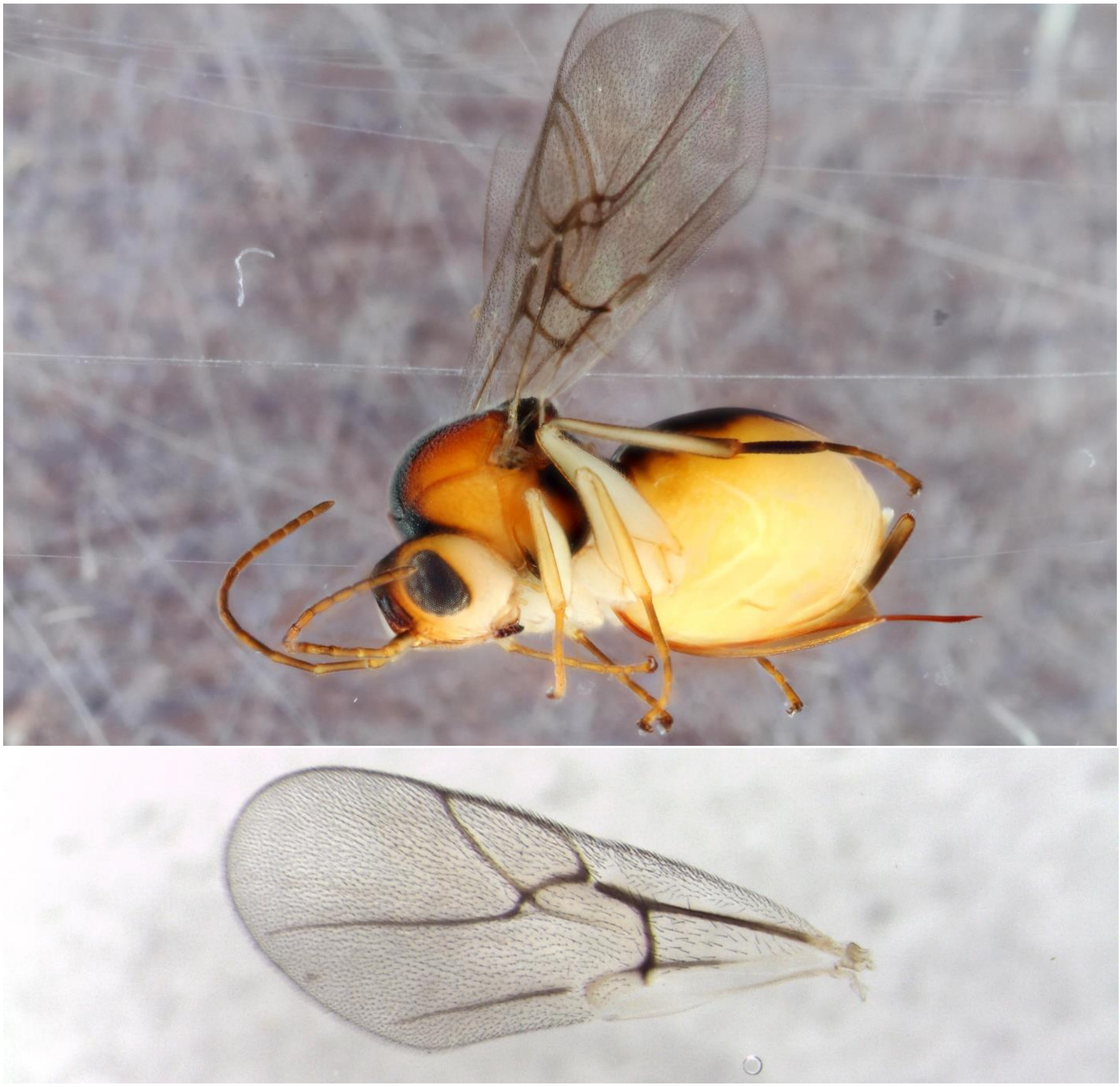

Figure S20. *Synergus oneratus* (Clade 10) – 500-011-002 –male
from *Andricus quercusstrobilanus* on *Quercus bicolor*, Iowa City, IA

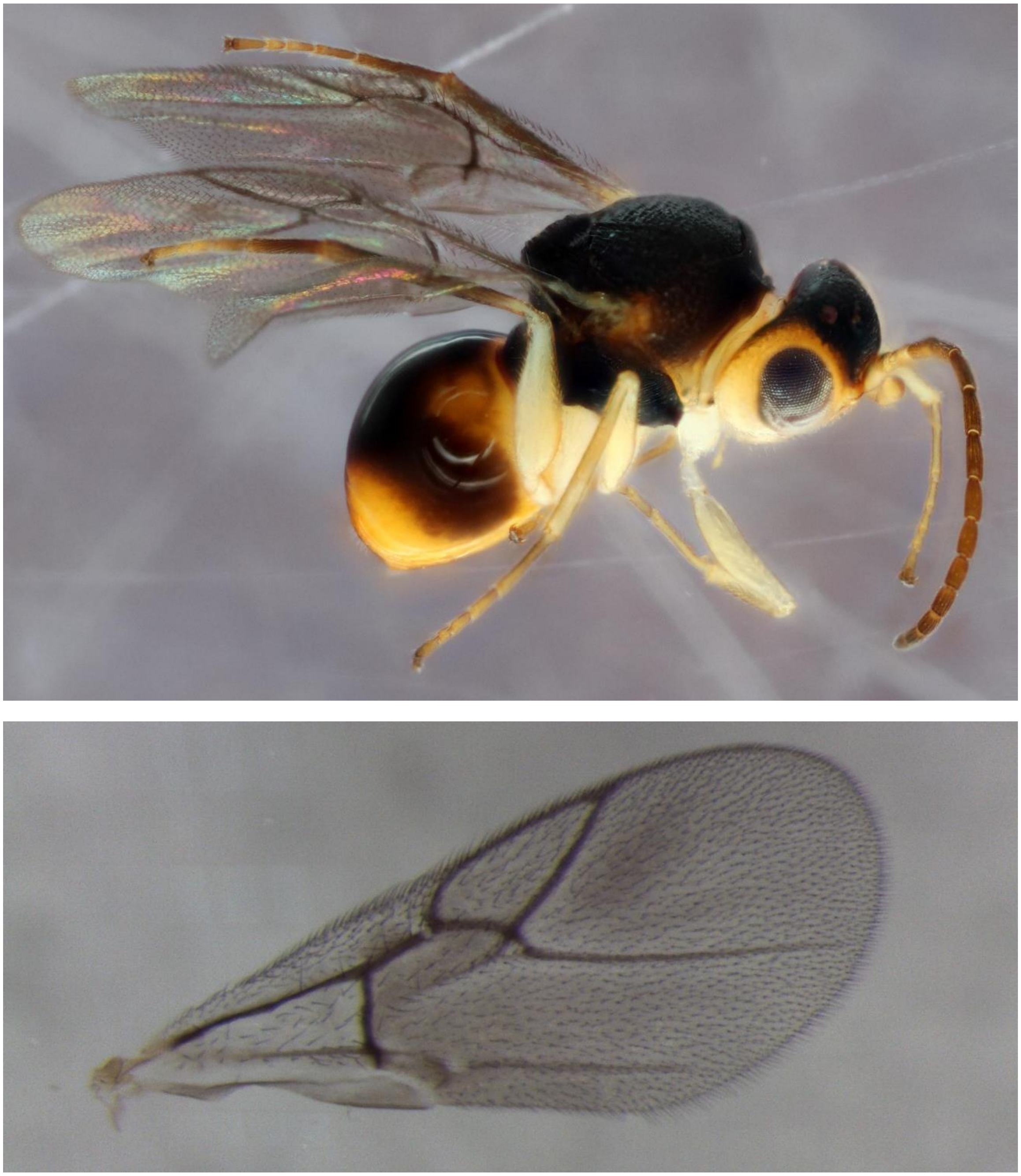

Figure S21. *Synergus oneratus* (Clade 11) – 335-019-001 – female from *Disholcaspis quercusglobulus* on *Quercus alba*, Iowa City, IA

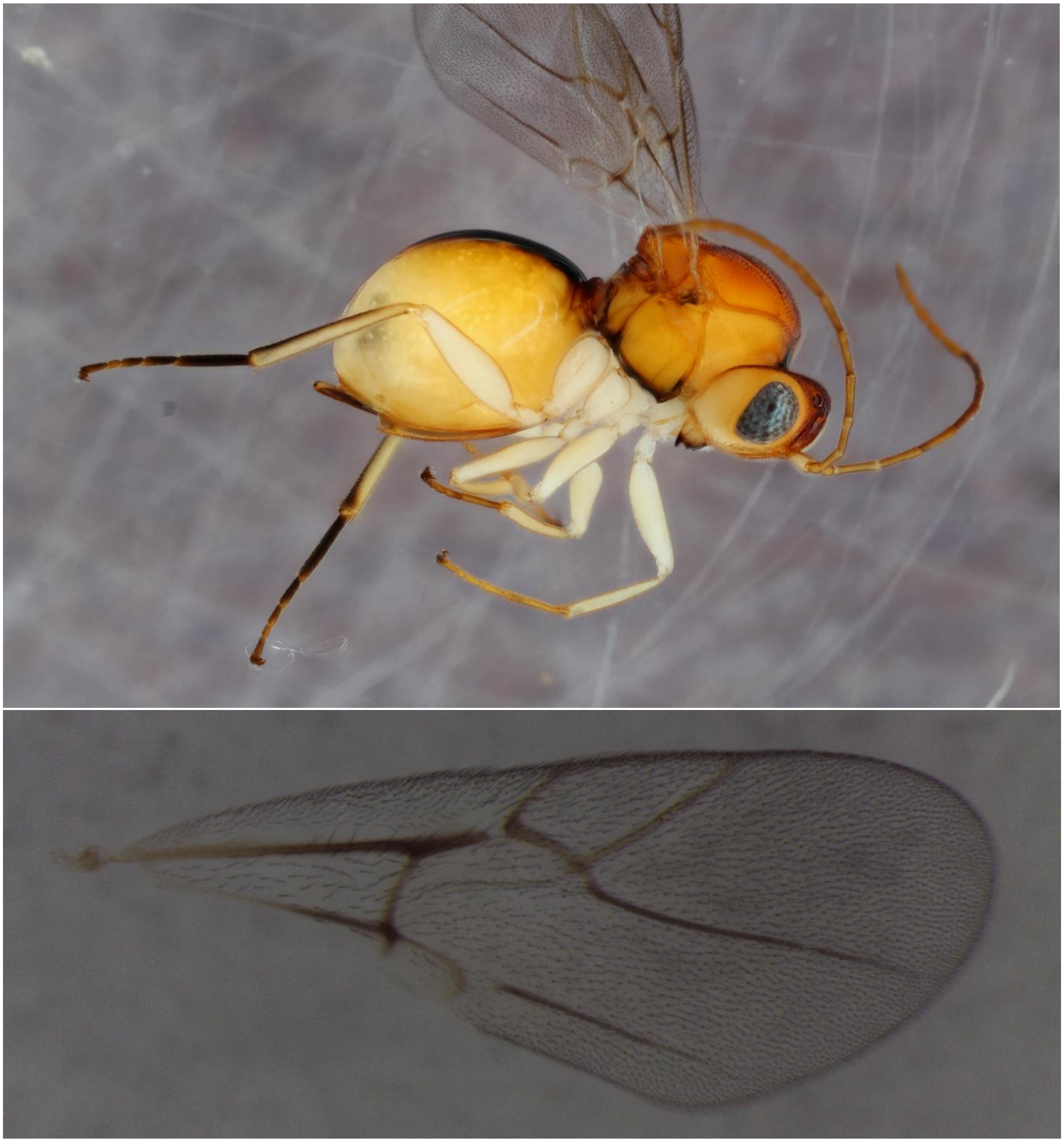

Figure S22. *Synergus oneratus* (Clade 11) – 1113-107-001 – male from unknown twig gall on *Quercus alba*, Nashville, TN

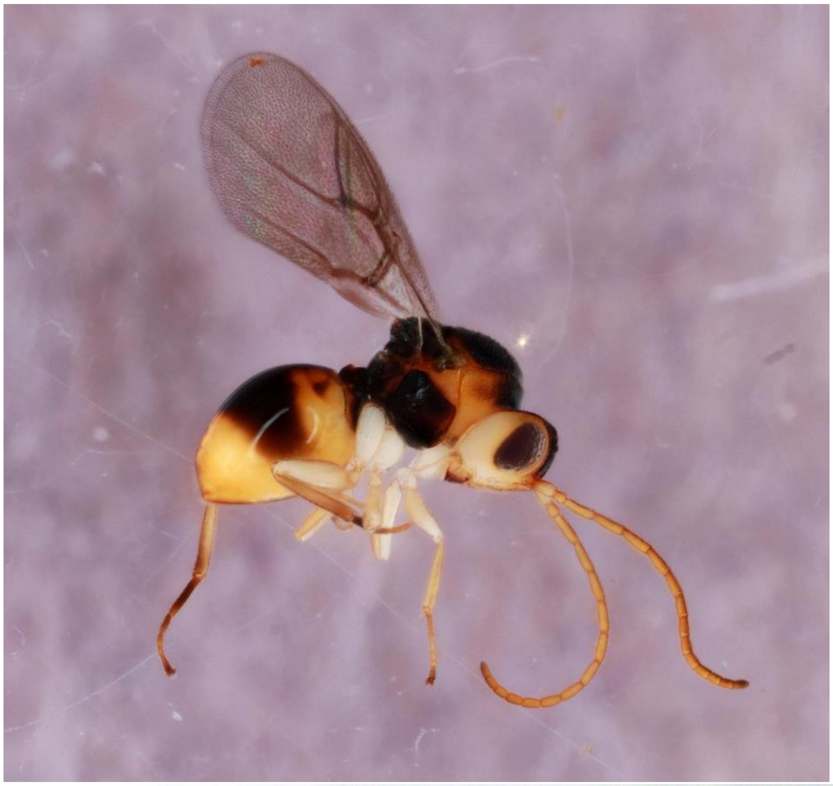

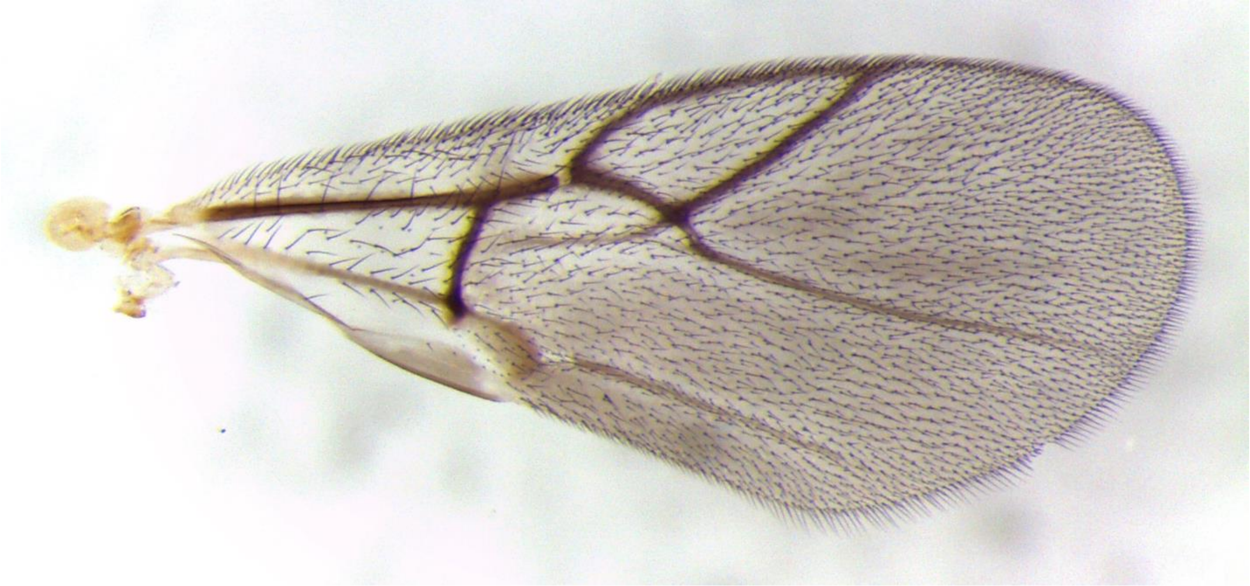

Figure S23. *Synergus* unk #2 (Clade 12) – 636-023-005 –male
from *Philonix nigra* on *Quercus macrocarpa*, Konza, KS

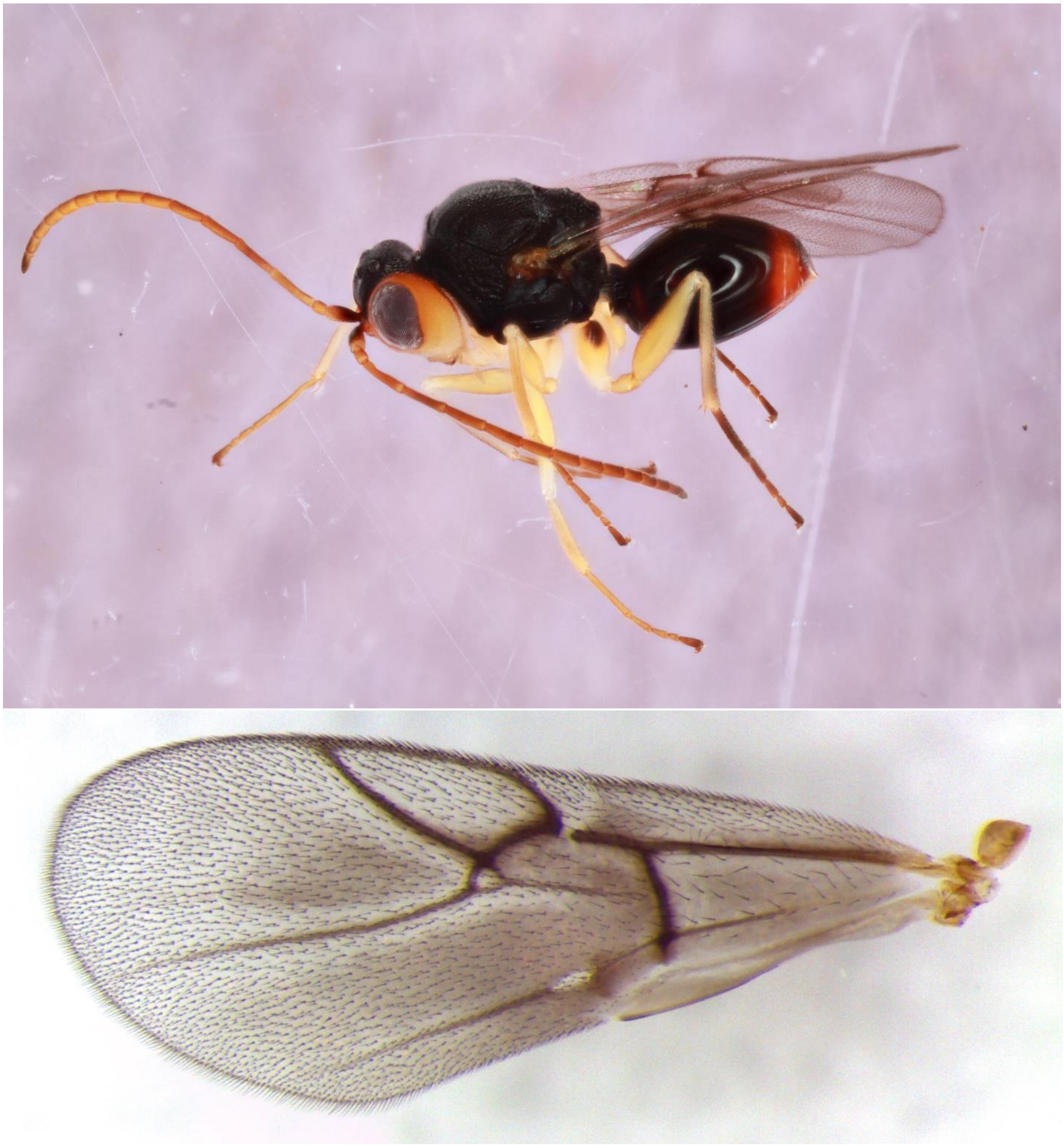

Figure S24. *Synergus villosus* (Clade 13) – 665-059-008B – female
from *Dryocosmus imbricariae* on *Quercus rubra*, Tiffin, IA

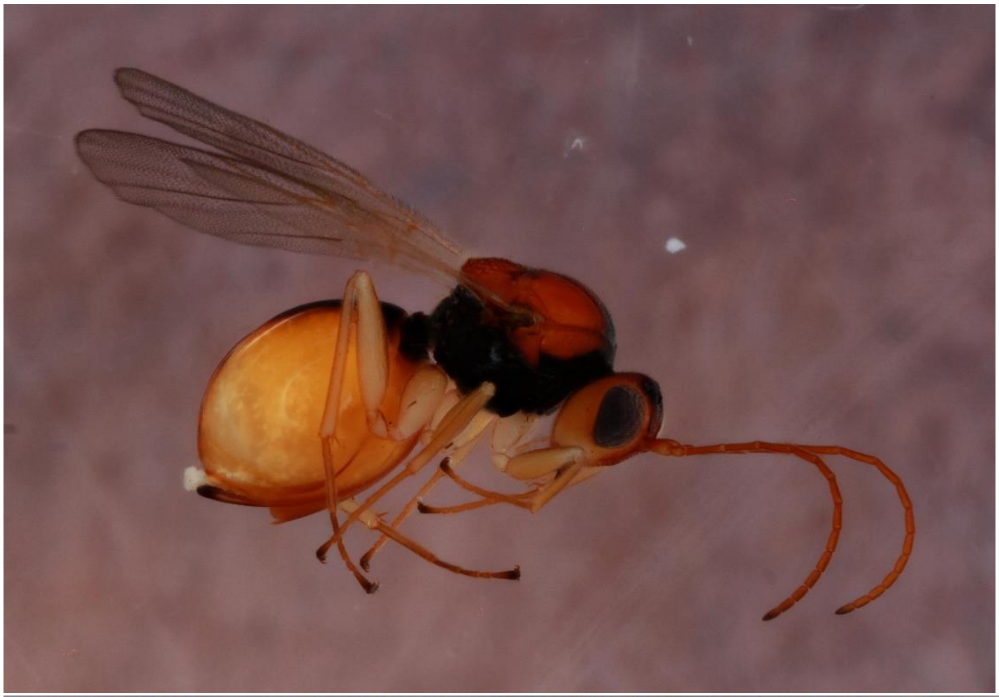

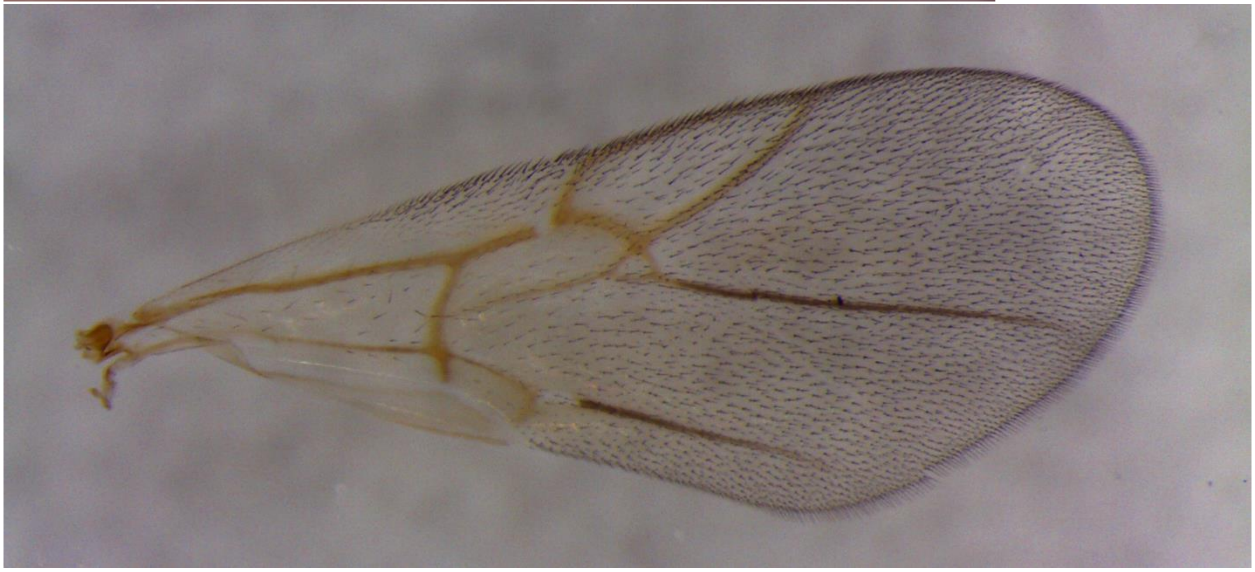

Figure S25. *Synergus villosus* (Clade 14) – 1206-004-002 – female
from *Acraspis villosa* on *Quercus macrocarpa*, Urbana, IL

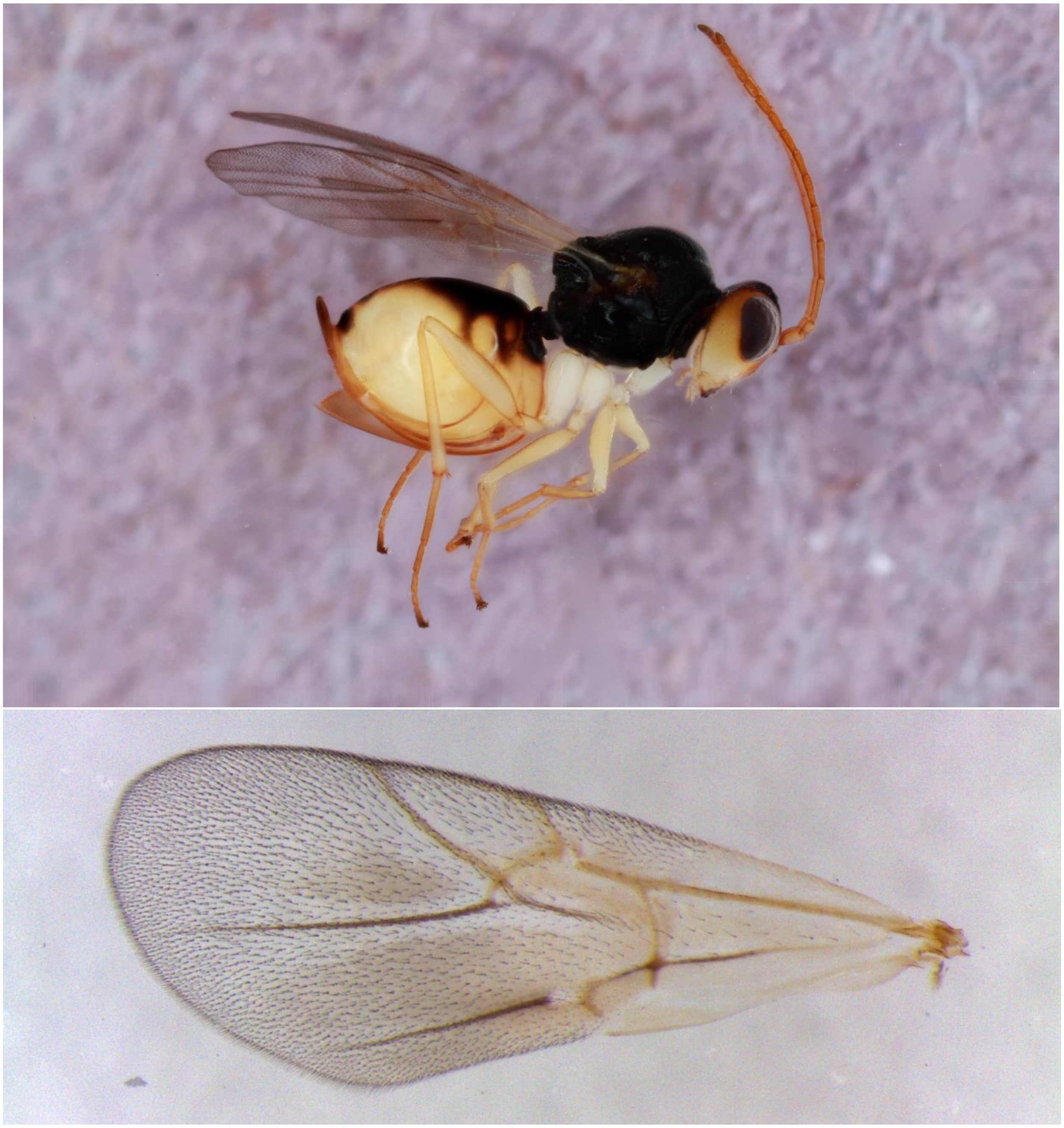

Figure S26. *Synergus magnus* (Clade 15) – 1137-110-002 – female
from *Amphibolips quercusjugulans* on *Quercus velutina*, Paris, TN

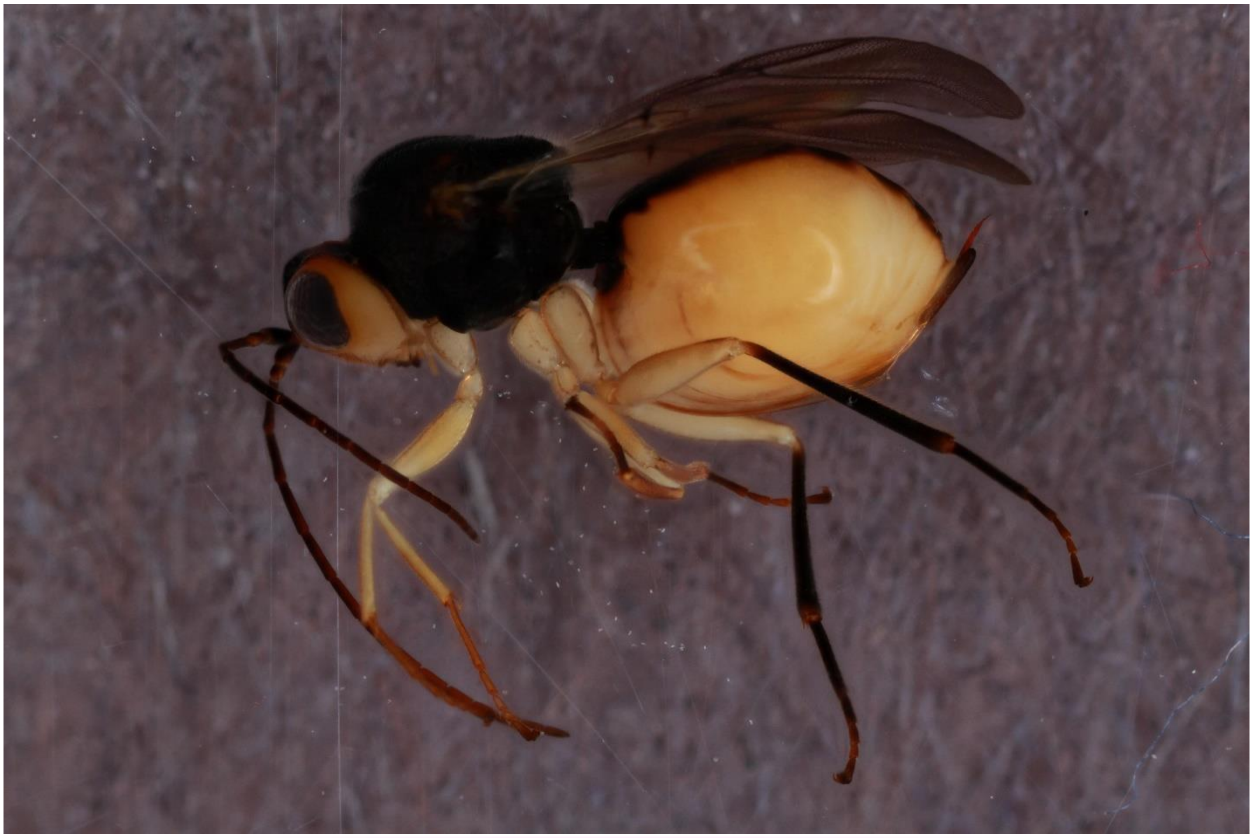

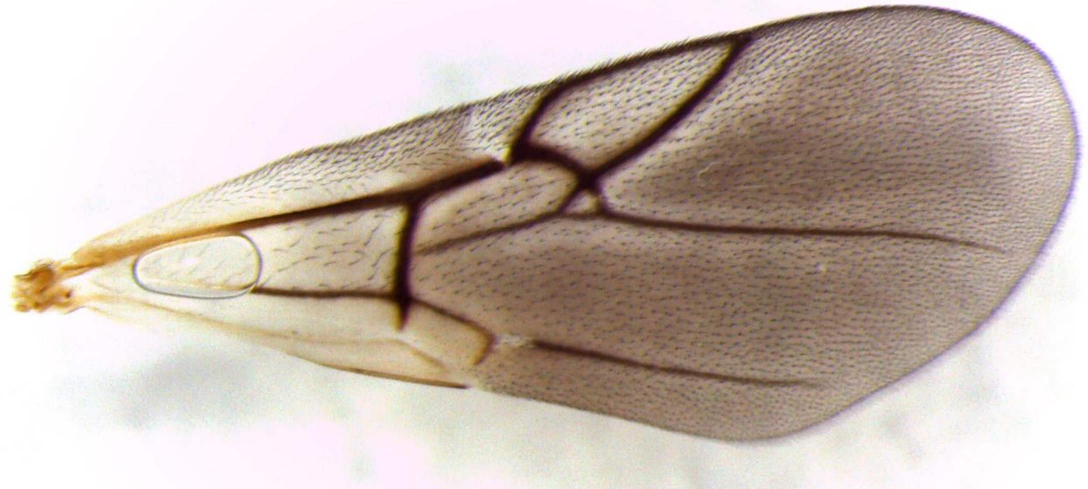

Figure S27. *Synergus erinacei* (Clade 16) – 1105-003-010A – female
from *Acraspis pezomachoides* on *Quercus alba*, Pedukah, KY

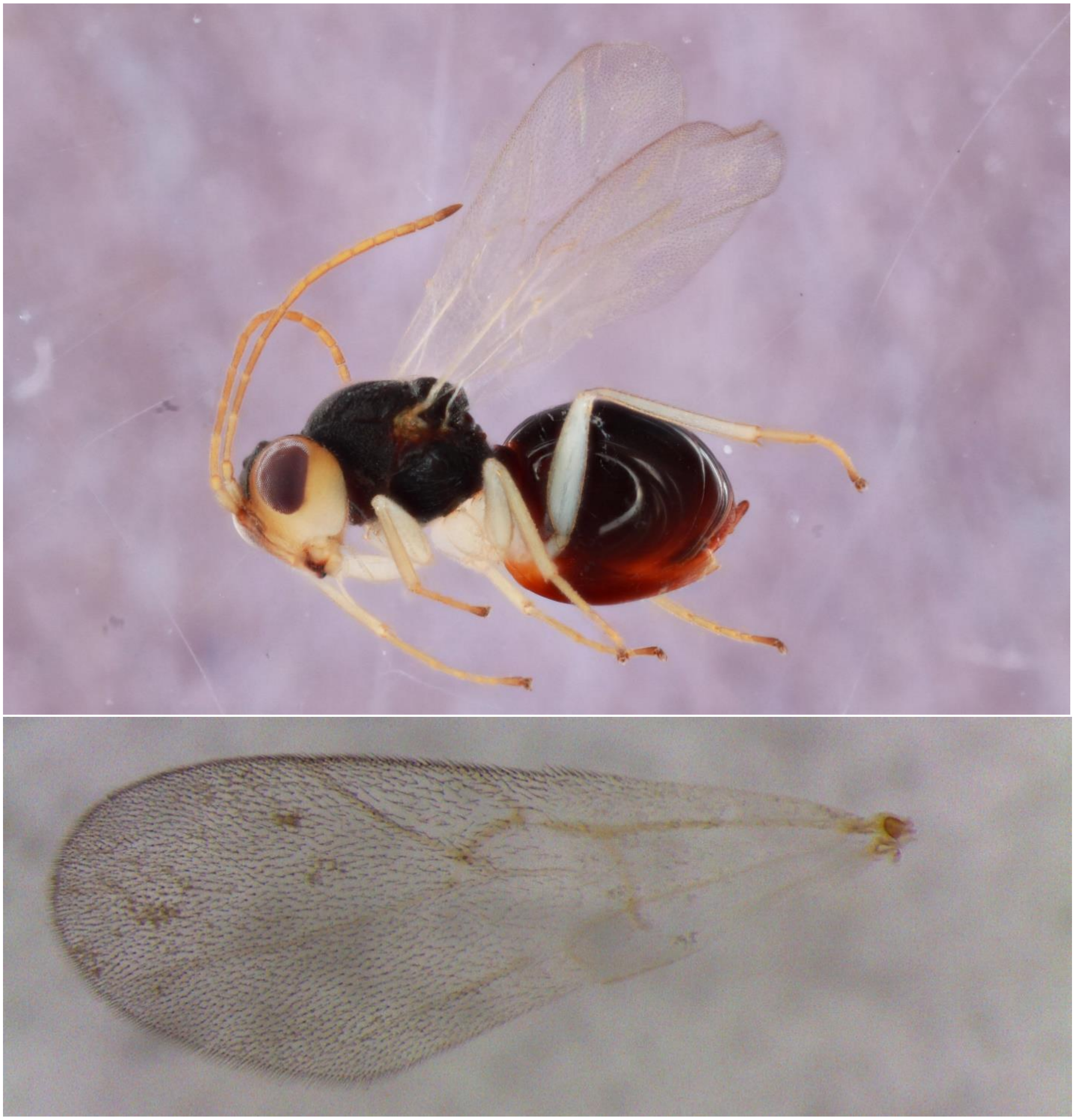

Figure S28. *Synergus erinacei* (Clade 16) – 602-003-016 – male
from *Acraspis pezomachoides* on *Quercus alba*, Urbana, IL

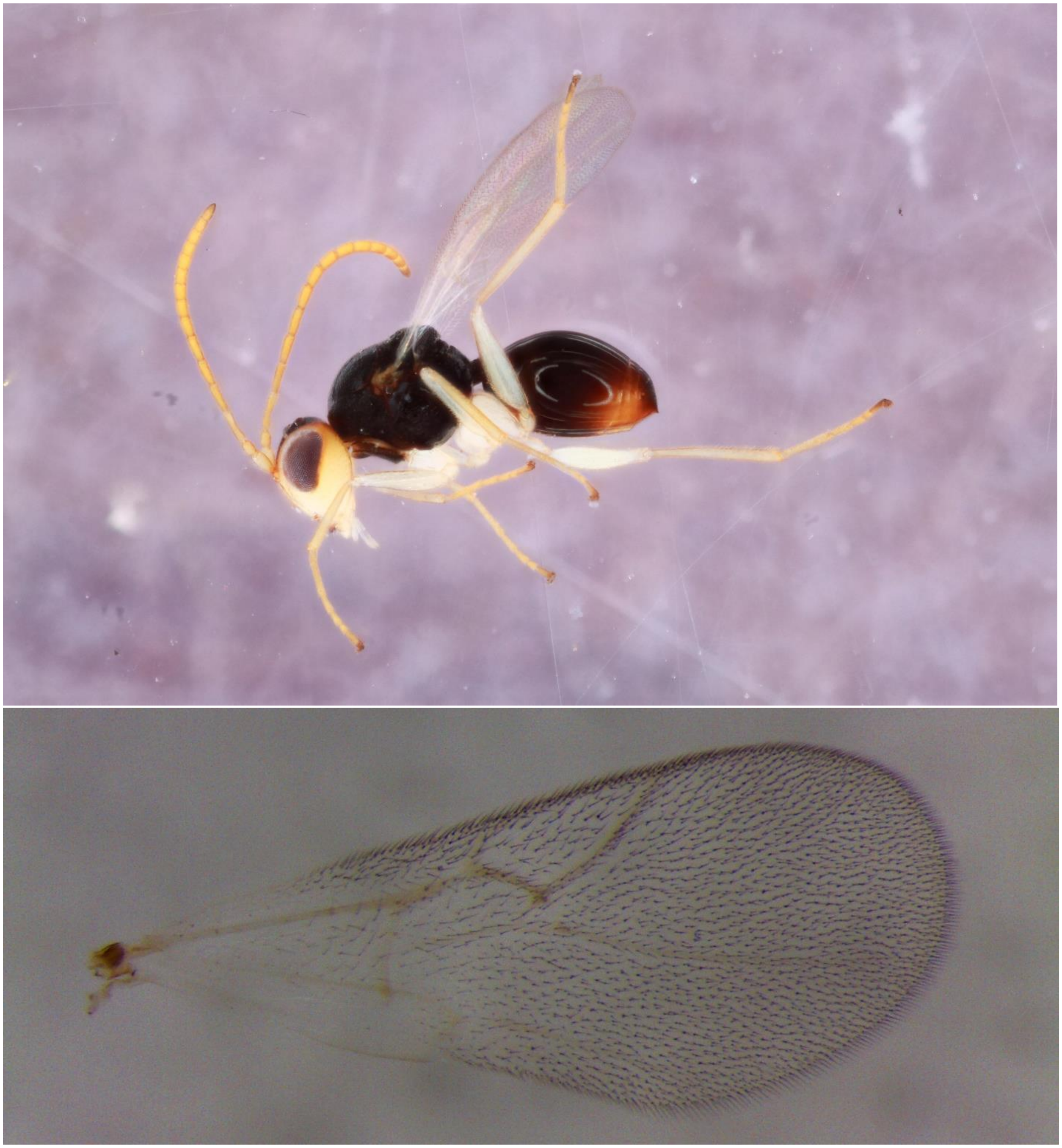

Figure S29. *Synergus erinacei* (Clade 17) – 338-001-002 – female
from *Acraspis macrocarpae* on *Quercus macrocarpa*, Iowa City, IA

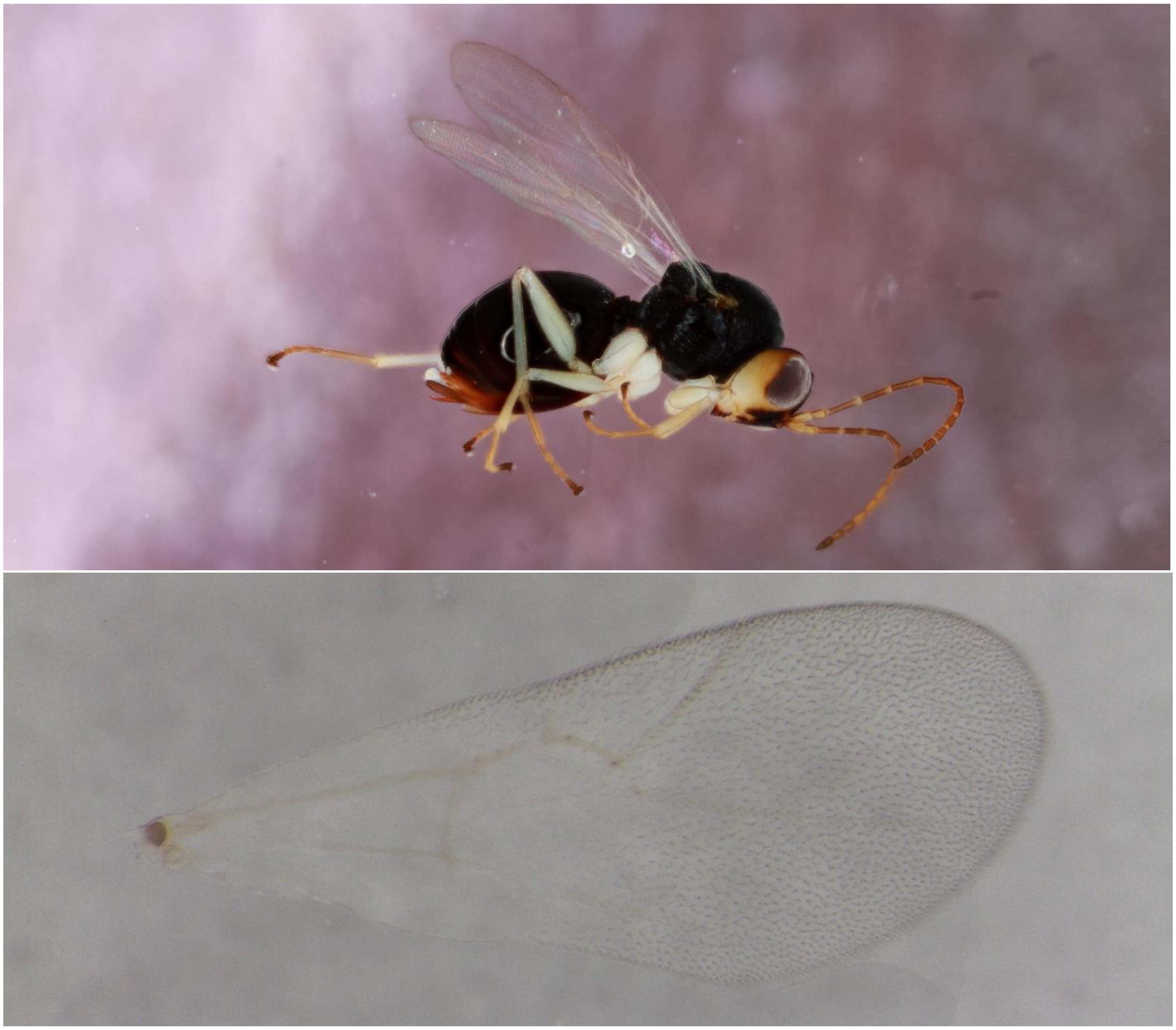

Figure S30. *Synergus walshii* (Clade 18) – 362-024-001A – female
from *Phylloteras poculum* on *Quercus bicolor*, Iowa City, IA

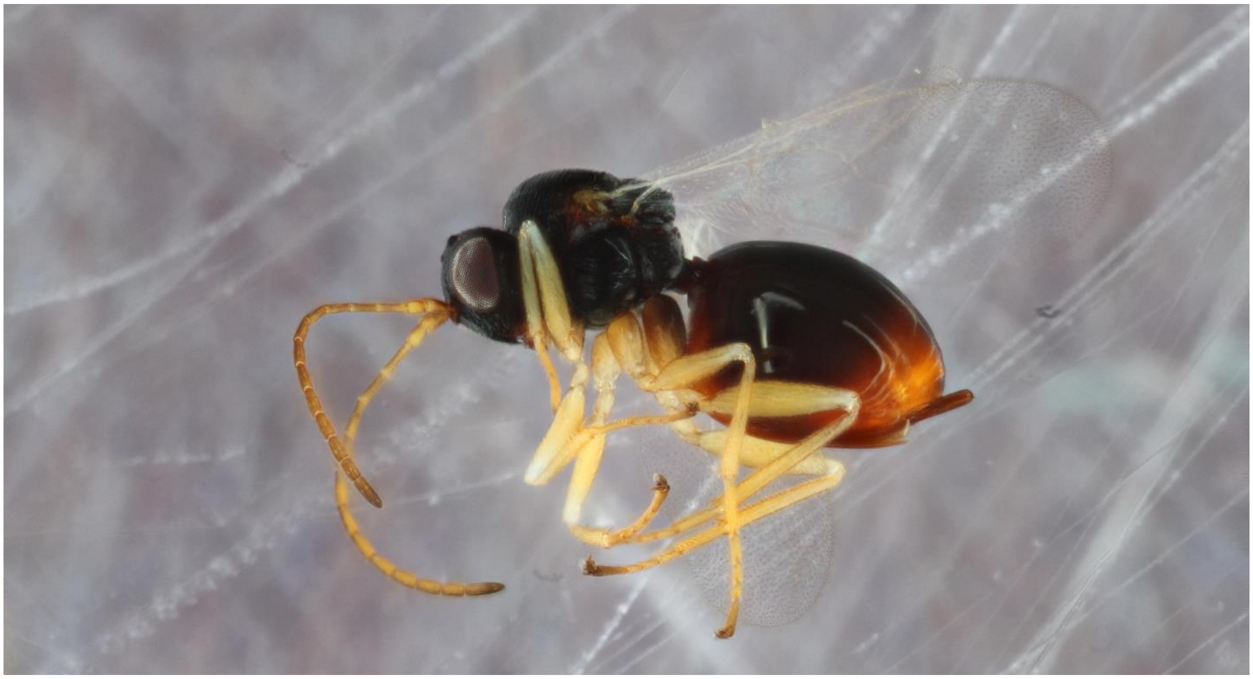

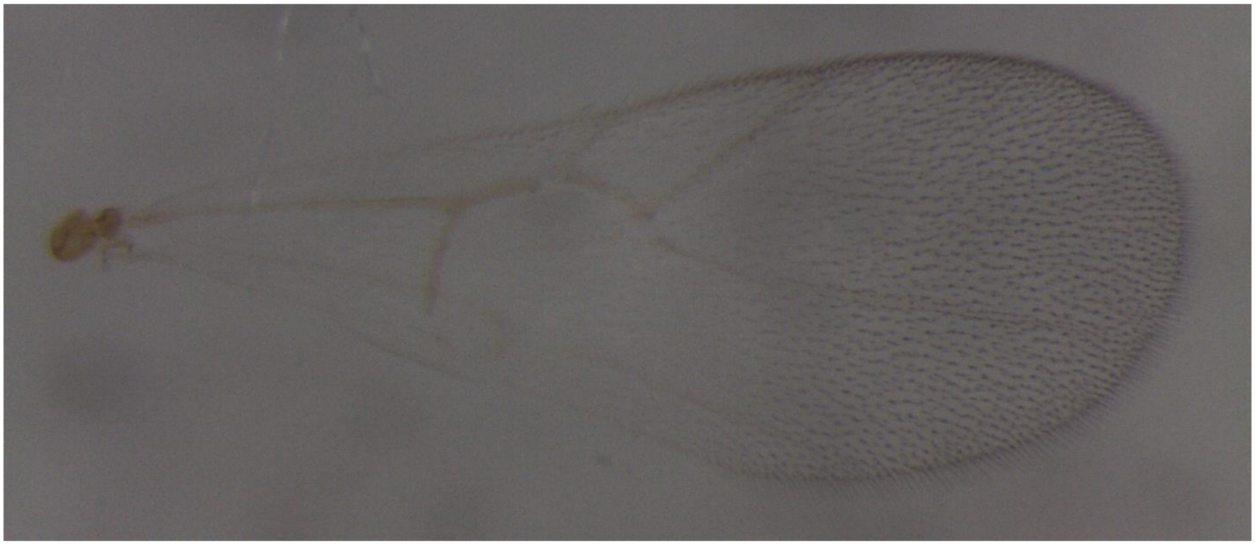

Figure S31. *Synergus walshii* (Clade 18) – 115-1-3 – male
from *Phylloteras volutellae* on *Quercus macrocarpa*, Spirit Lake, IA

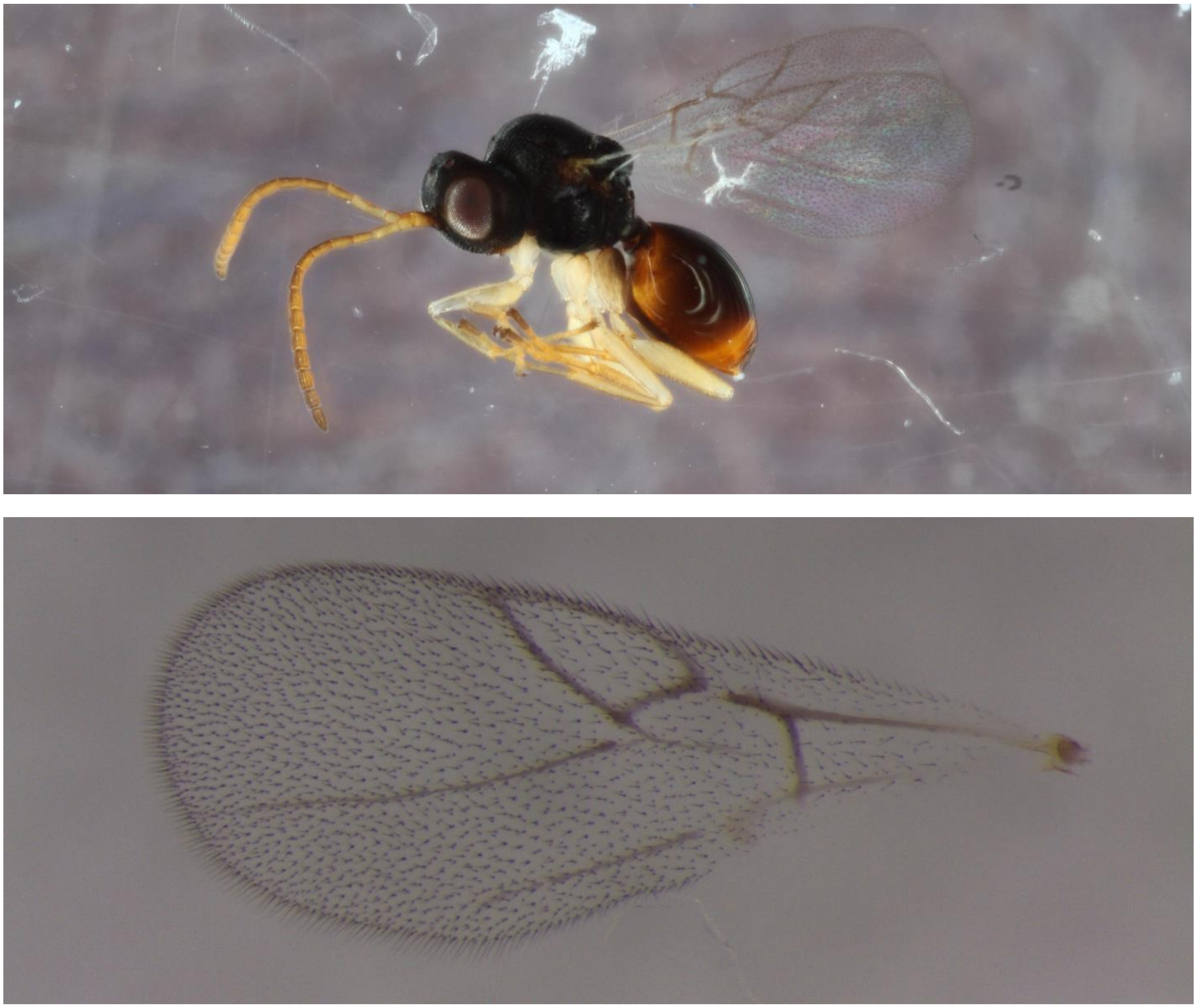

Figure S32. *Synergus walshii* (Clade 19) – 1153-081-003 – female
from *Andricus pattoni* on *Quercus stellata*, Bloomsdale, MO

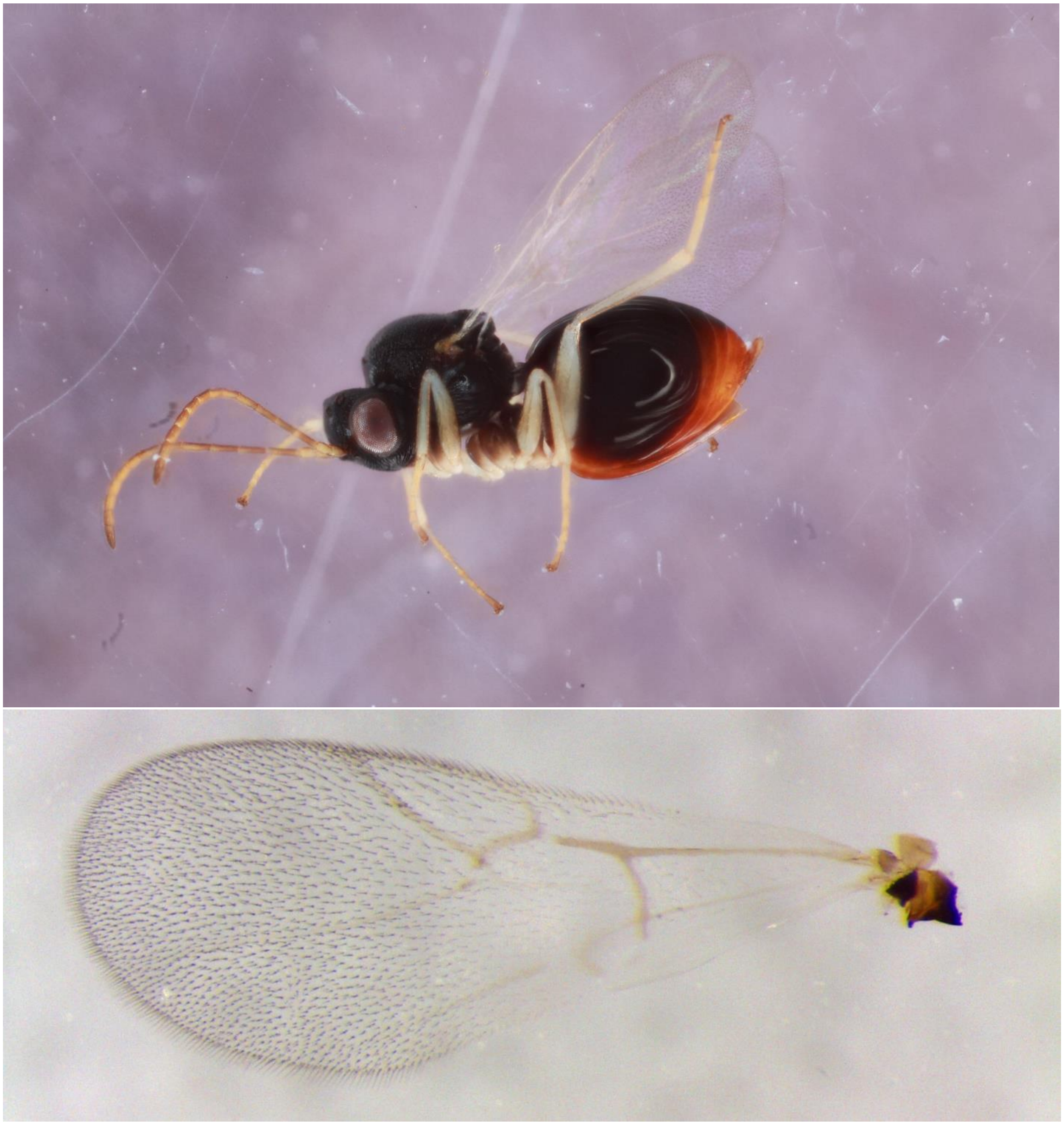

Figure S33. *Synergus walshii* (Clade 19) – 703-061-002A – male
from *Andricus quercusflocci* on *Quercus alba*, Iowa City, IA

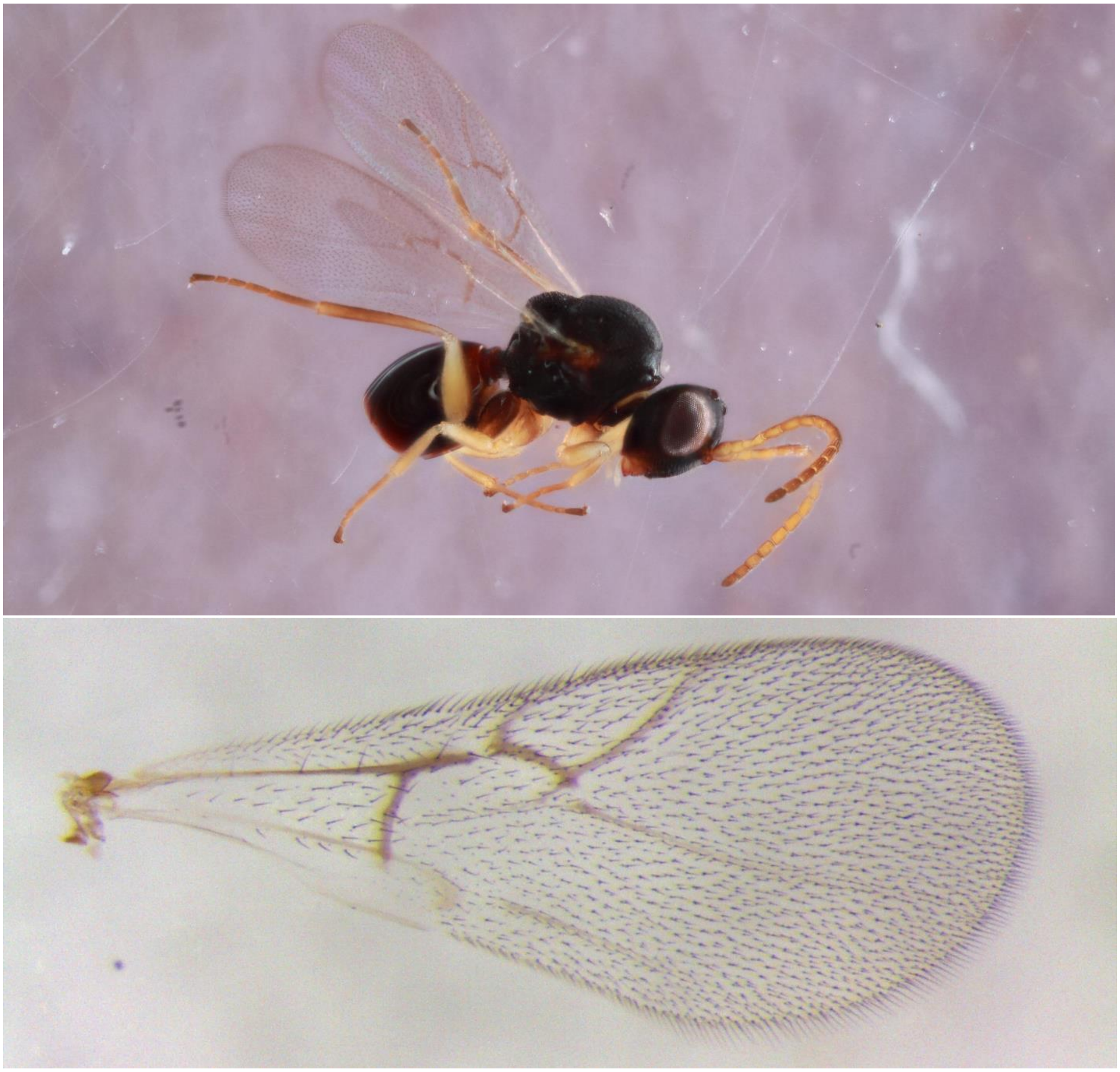

Figure S34. *Synergus punctatus* (Clade 20) – 642-009-023 – female
from *Andricus nigricens* on *Quercus bicolor*, Iowa City, IA

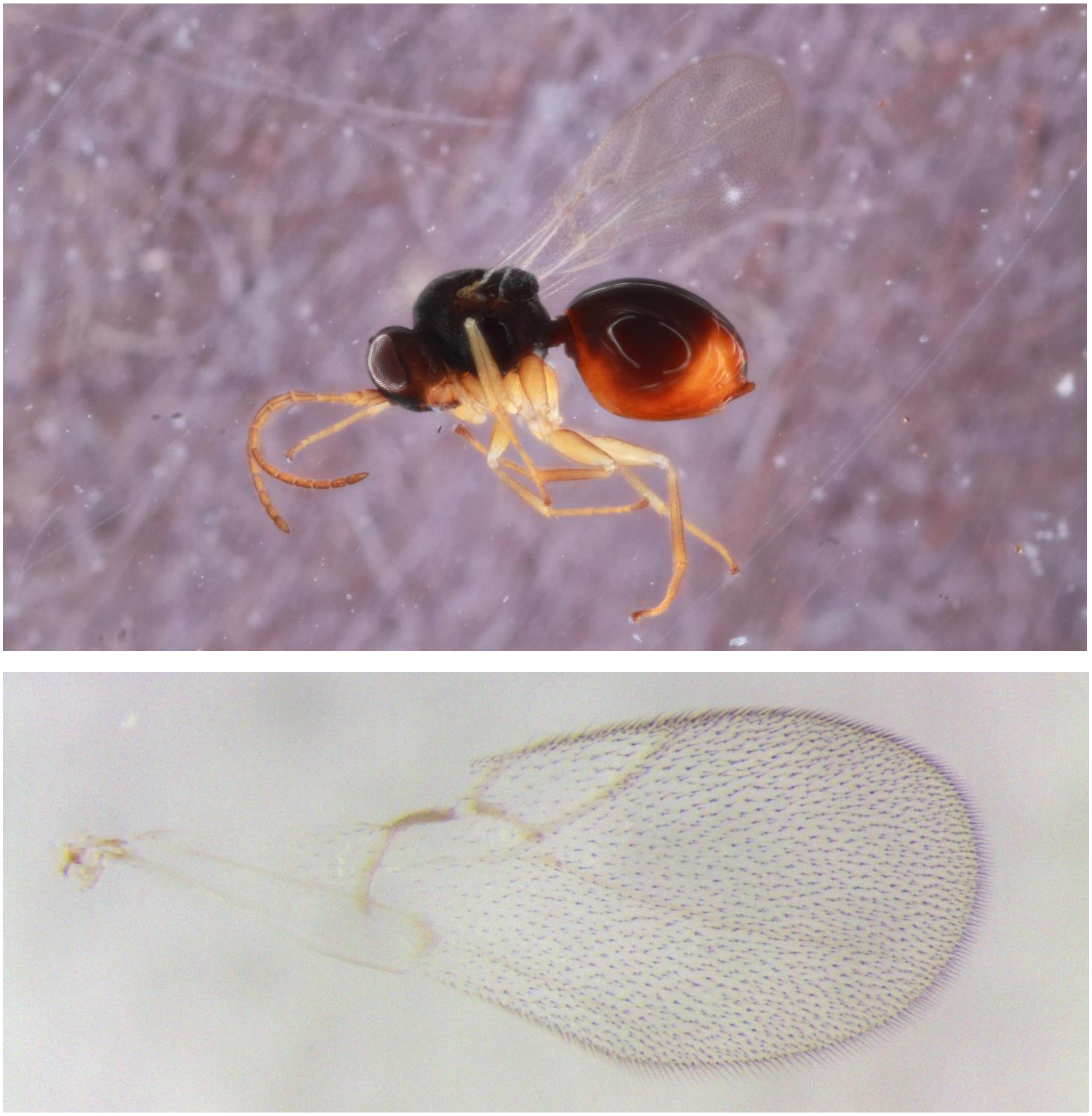

Figure S35. *Synergus punctatus* (Clade 20) – 642-009-015 –male
from *Andricus nigricens* on *Quercus bicolor*, Iowa City, IA

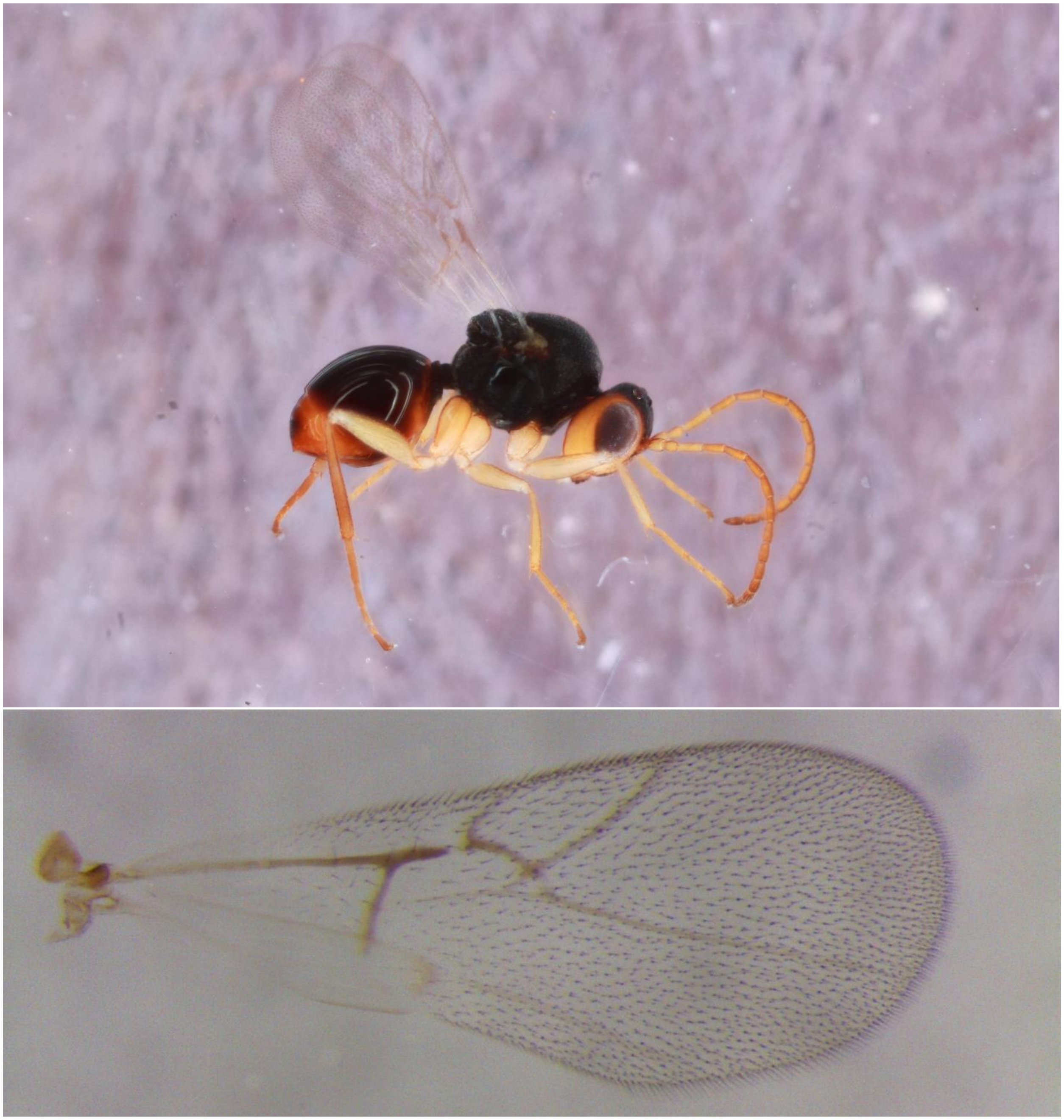

Figure S36. *Synergus* unk. #3 (Clade 21) – 555-019-042F – female
from *Disholcaspis quercusglobulus* on *Quercus stellata*, Paducah, KY

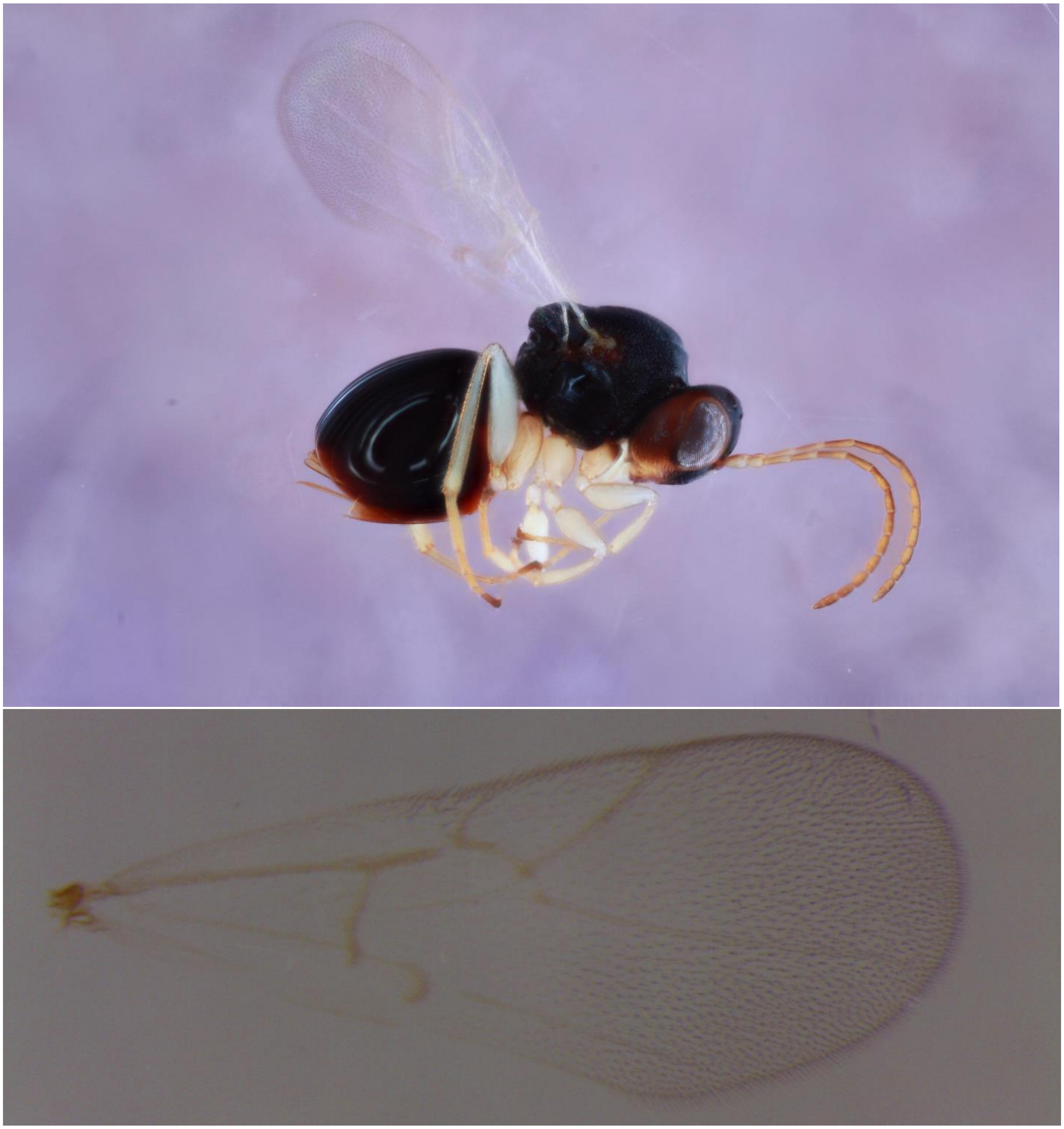

Figure S37. *Synergus punctatus* (Clade 22) – 1025-023-002B – female
from *Philonix nigra* on *Quercus macrocarpa*, Iowa City, IA

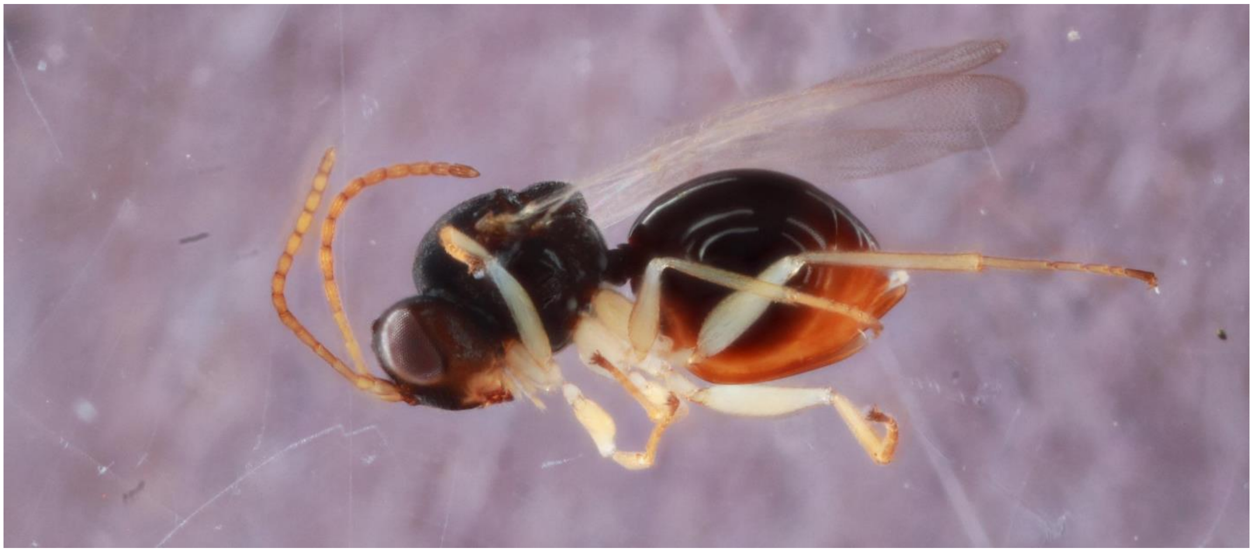

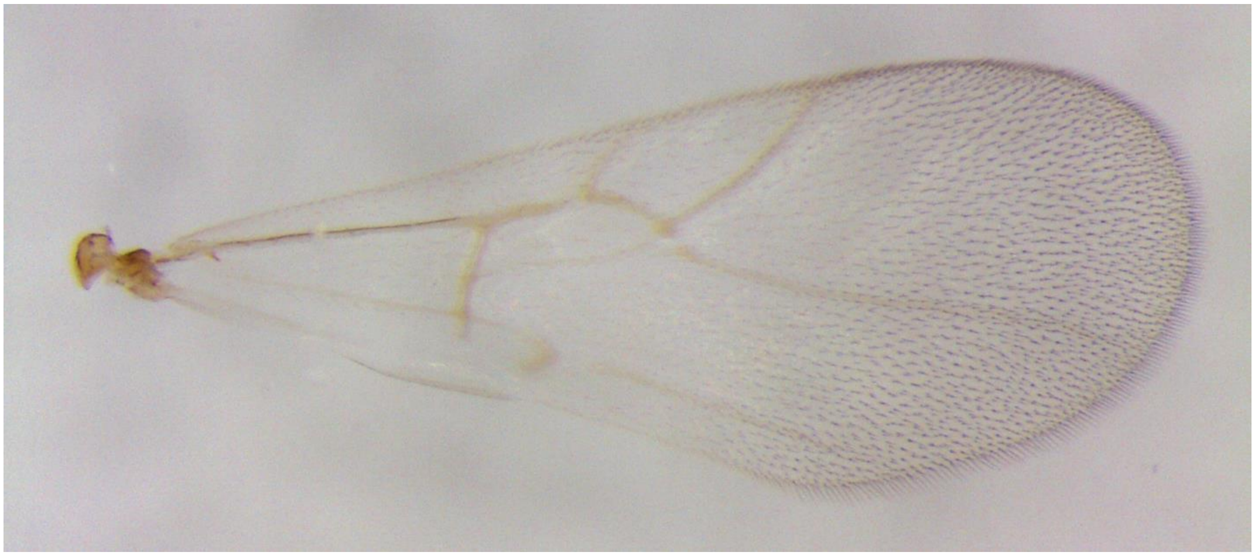

Figure S38. *Synergus punctatus* (Clade 22) – 636-023-005 – male
from *Philonix nigra* on *Quercus macrocarpa*, Konza, KS

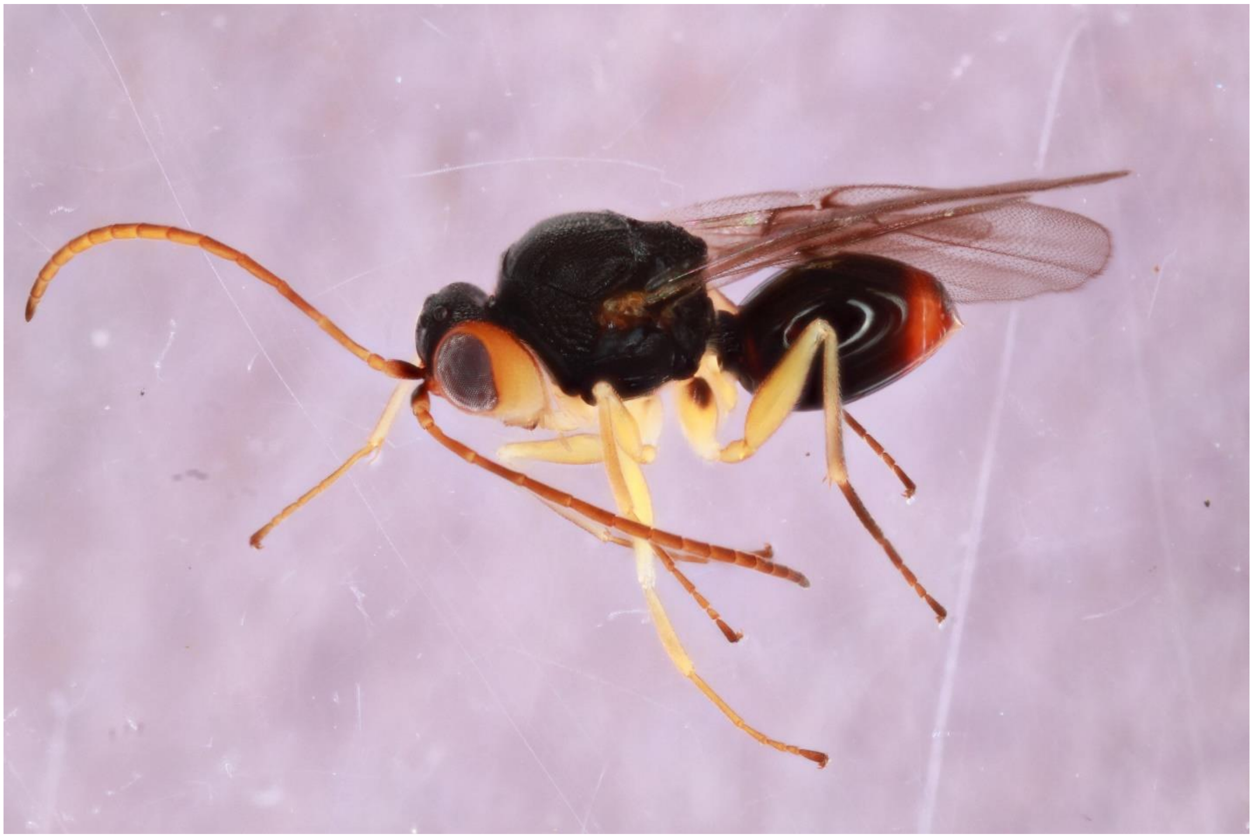

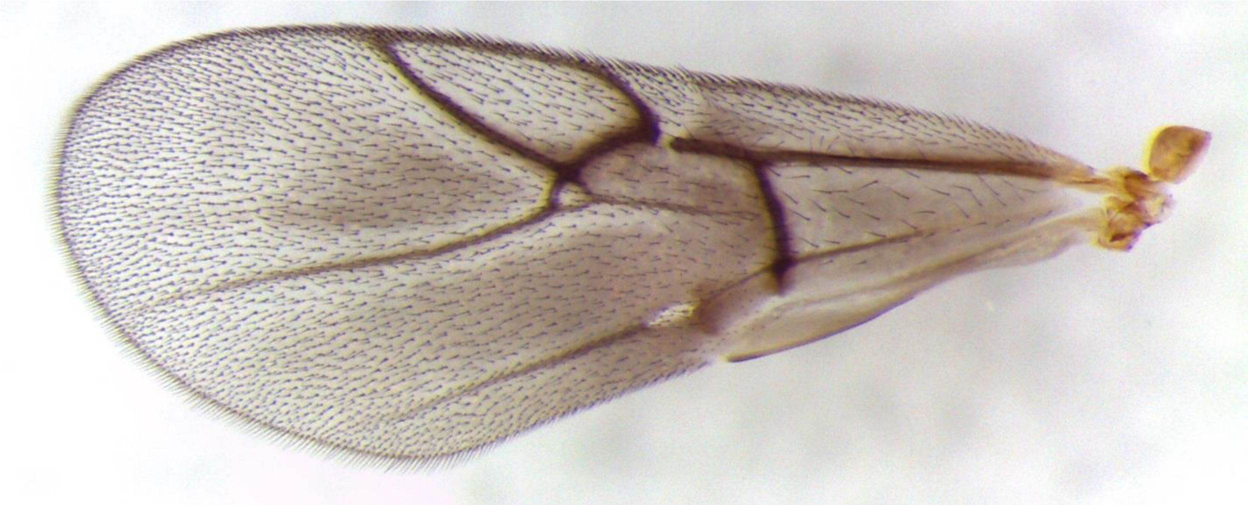

Figure S39. *Synergus punctatus* (Clade 23) – 539-067-002A – female
from *Andricus robustus* on *Quercus stellata*, St. Louis, MO

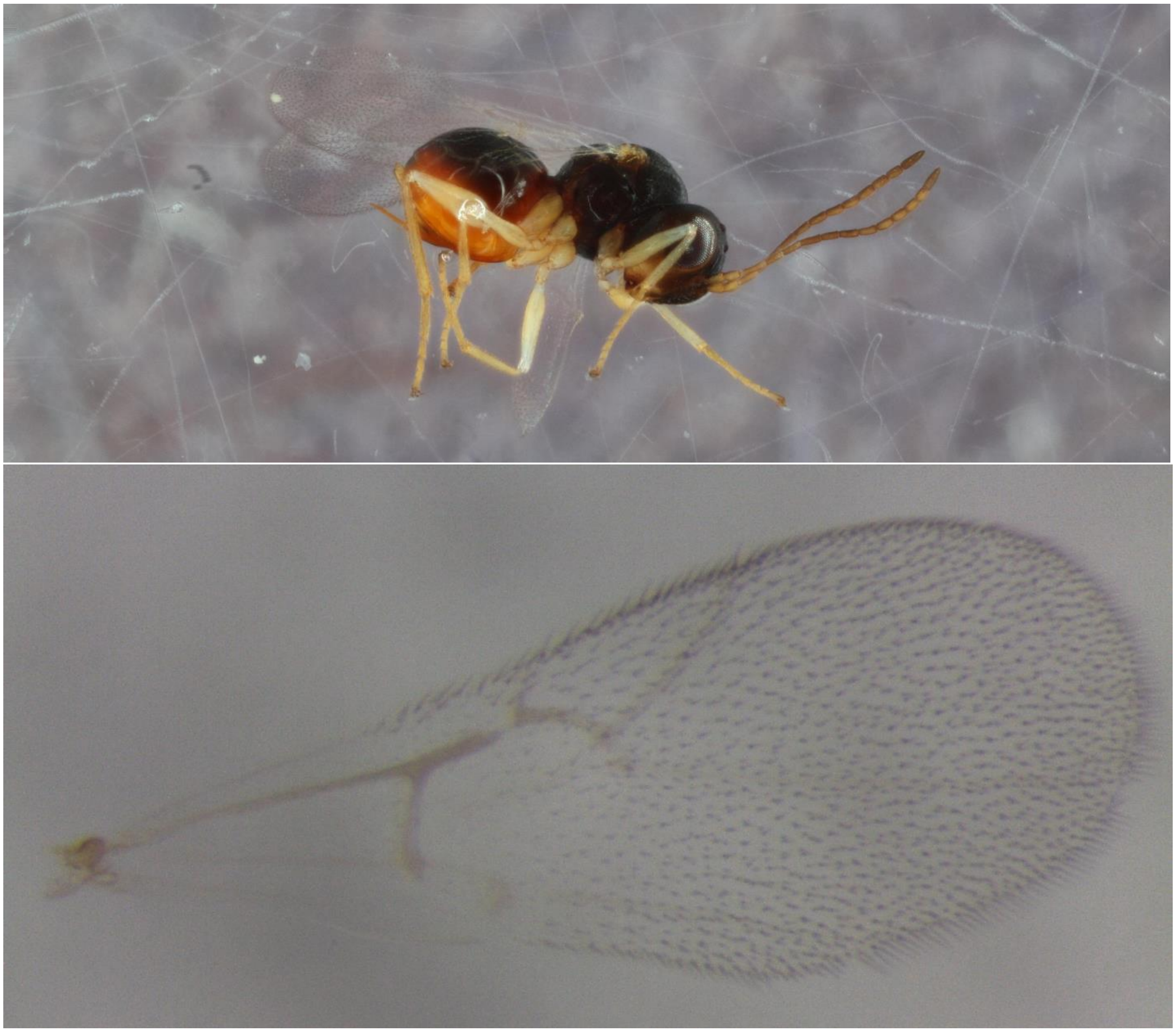

Figure S40. *Synergus punctatus* (Clade 23) – 539-067-003B –male (no wing picture) from *Andricus robustus* on *Quercus stellata*, St. Louis, MO

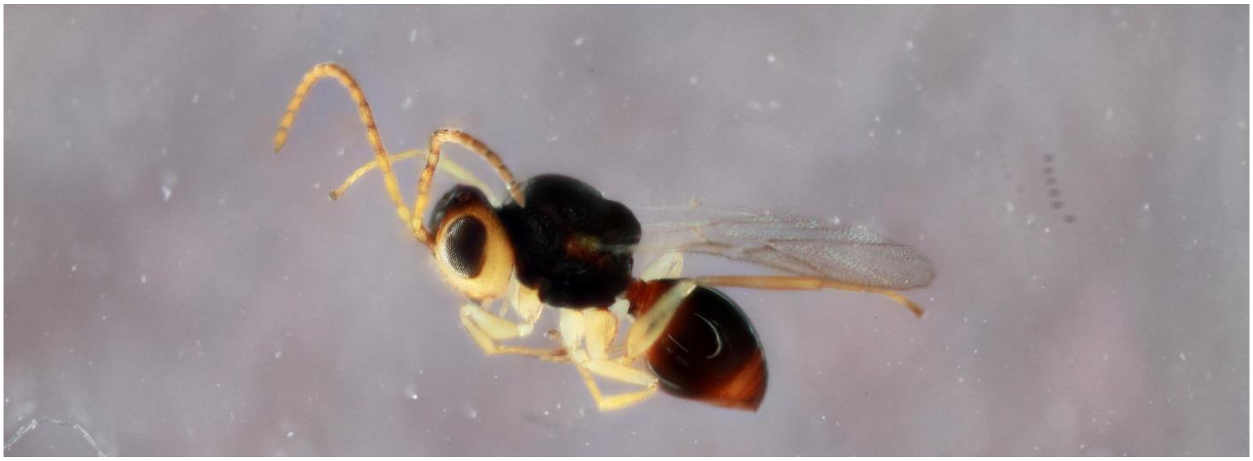

Figure S41. *Synergus punctatus* (Clade 24) – 547-002-003 – female (no wing picture) from *Acraspis erinacei* on *Quercus alba*, Paducah, KY

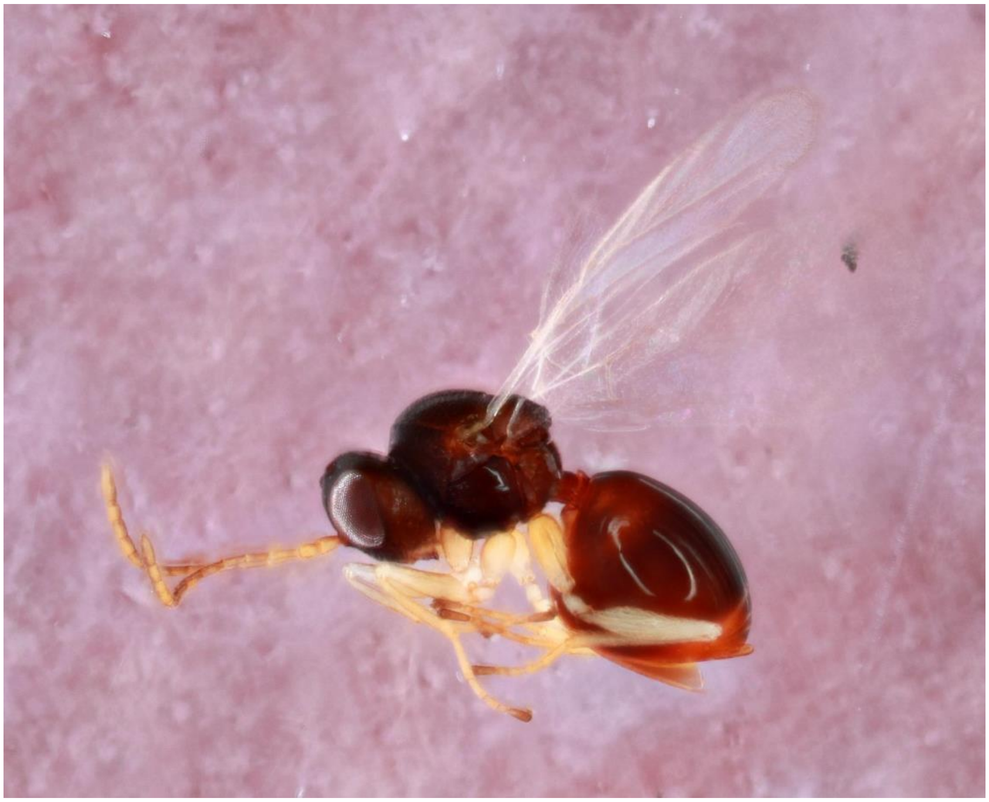

Figure S42. *Synergus campanula* (Clade 25) – 631-005-003B – female
from *Andricus dimorphus* on *Quercus prinoides*, Konza, KS

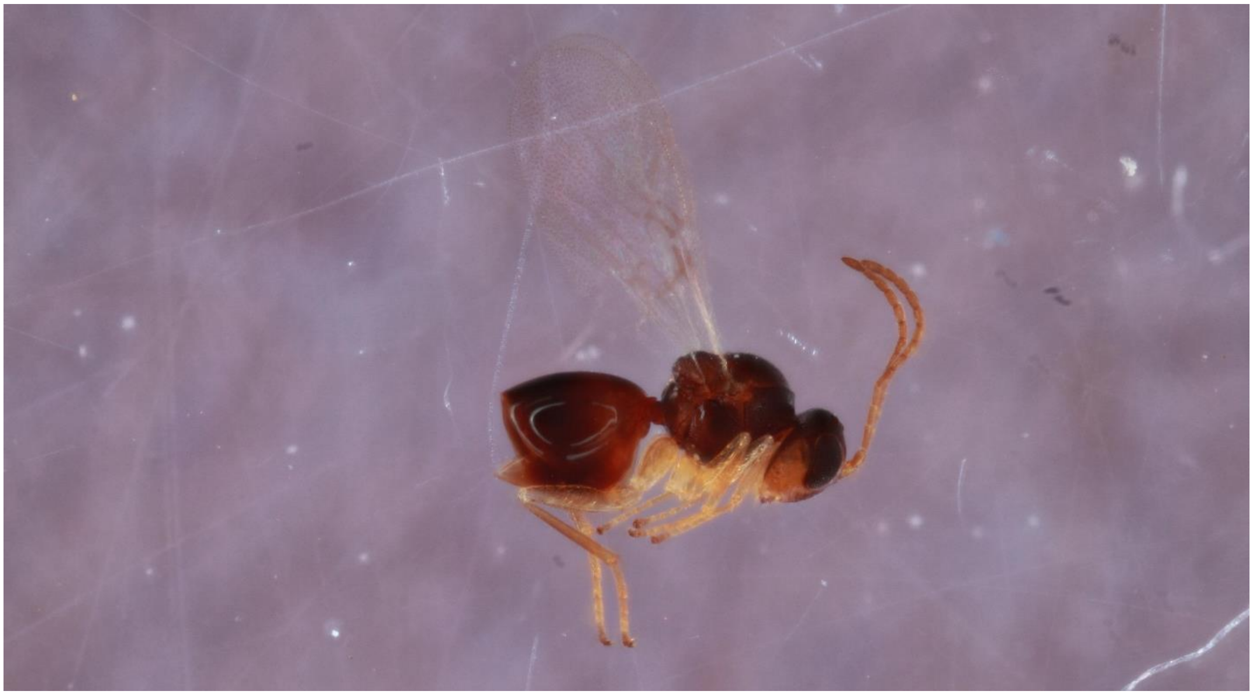

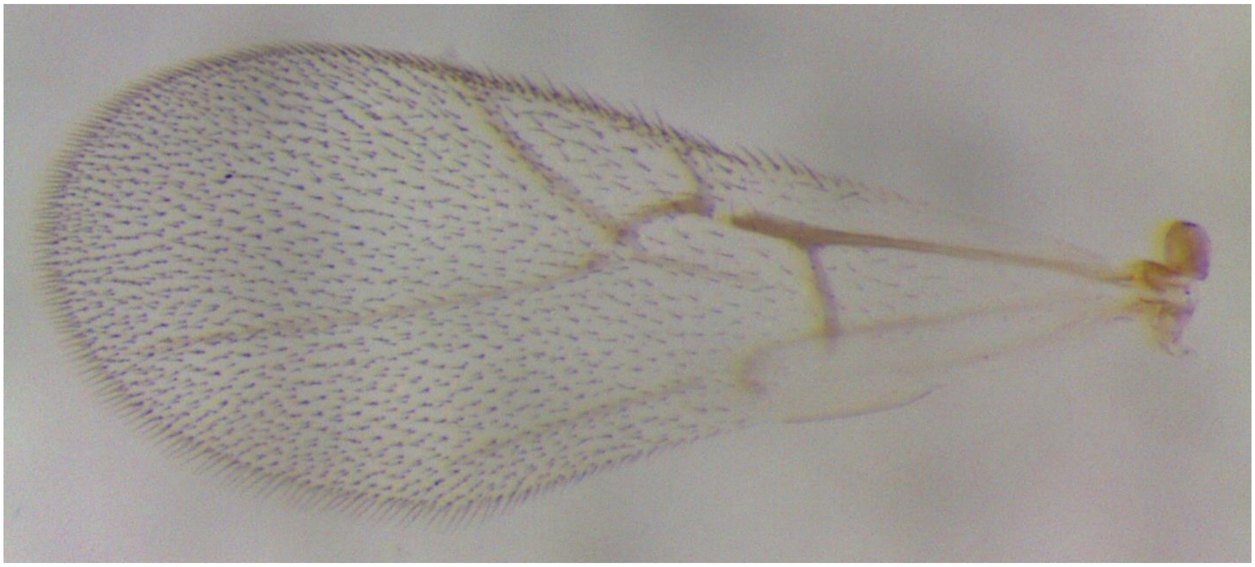

Figure S43. *Synergus campanula* (Clade 26) – 555-019-042F – female
from *Disholcaspis quercusglobulus* on *Quercus stellata*, Paducah, KY

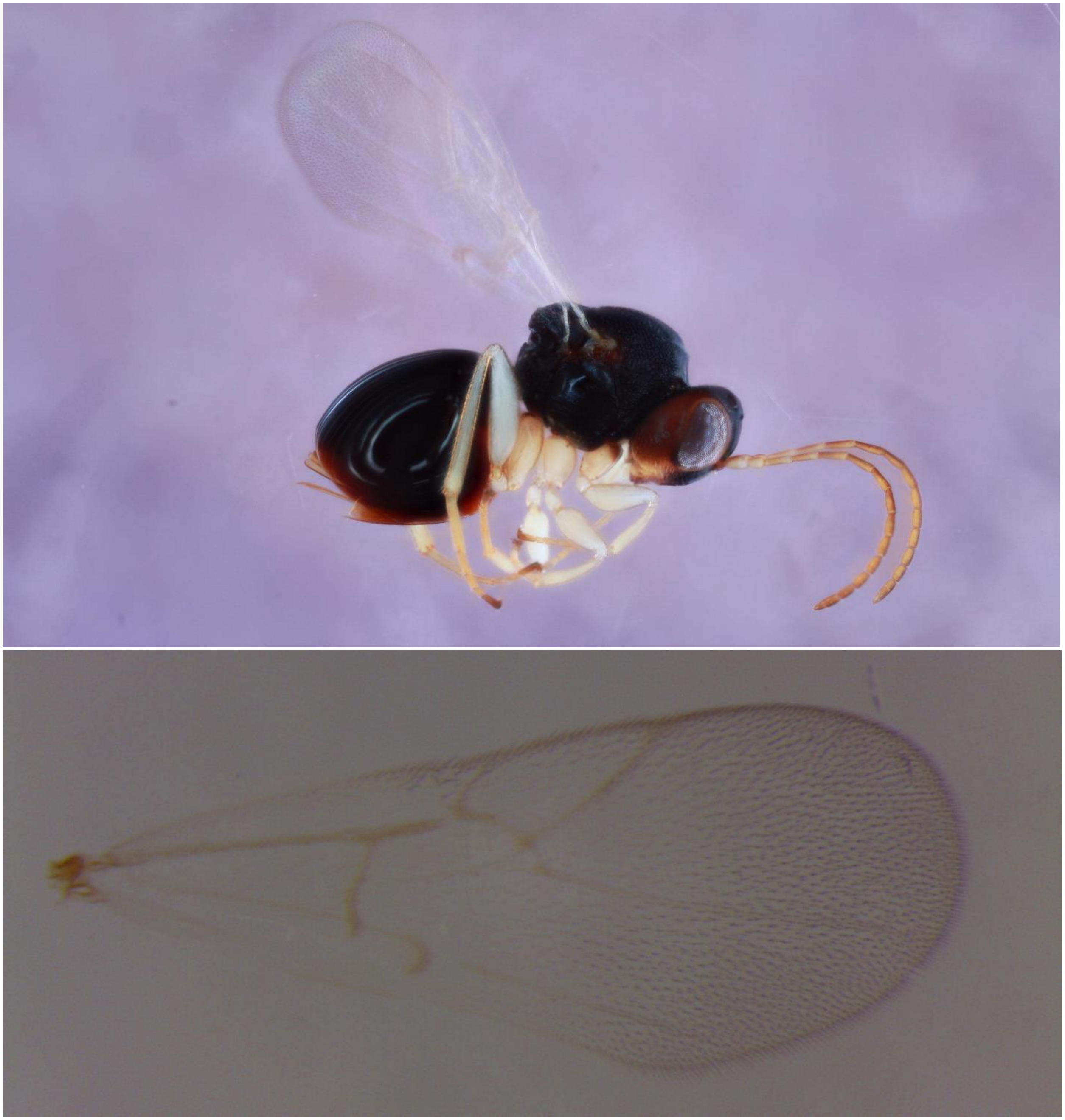

Figure S44. *Synergus campanula* (Clade 26) – 122-020-013 – male
from *Disholcaspis quercusmamma* on *Quercus bicolor*, Iowa City, IA

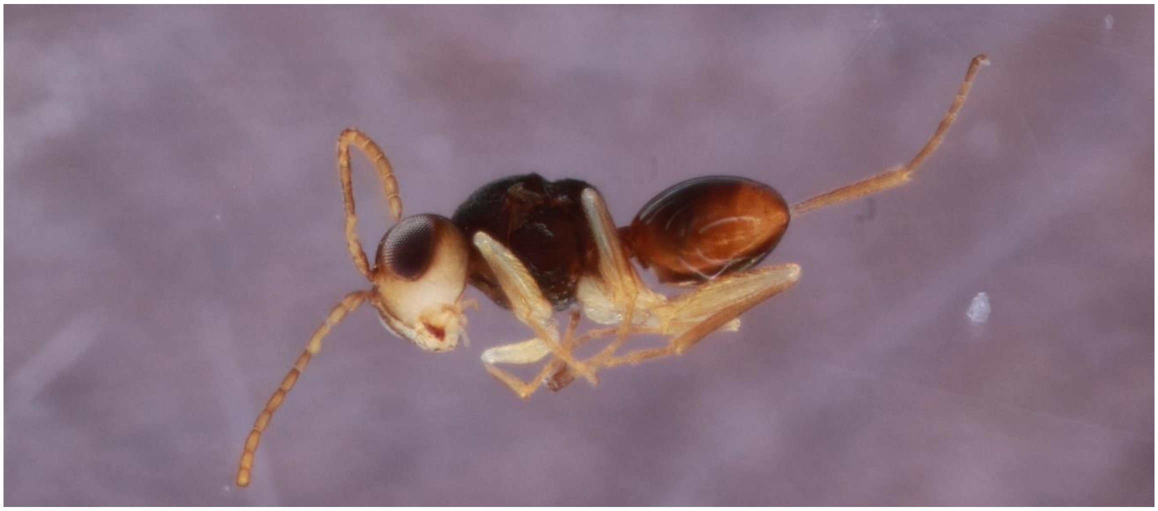

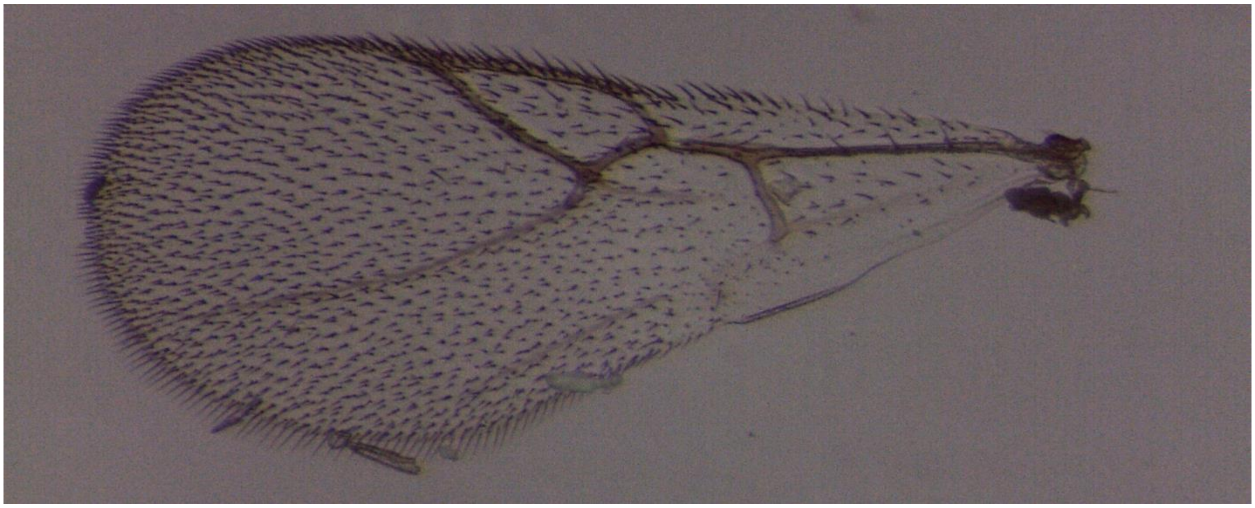

Figure S45. *Synergus campanula* (Clade 27) – 1105-003-001 – female from *Acraspis pezomachoides* on *Quercus alba*, Paducah, KY

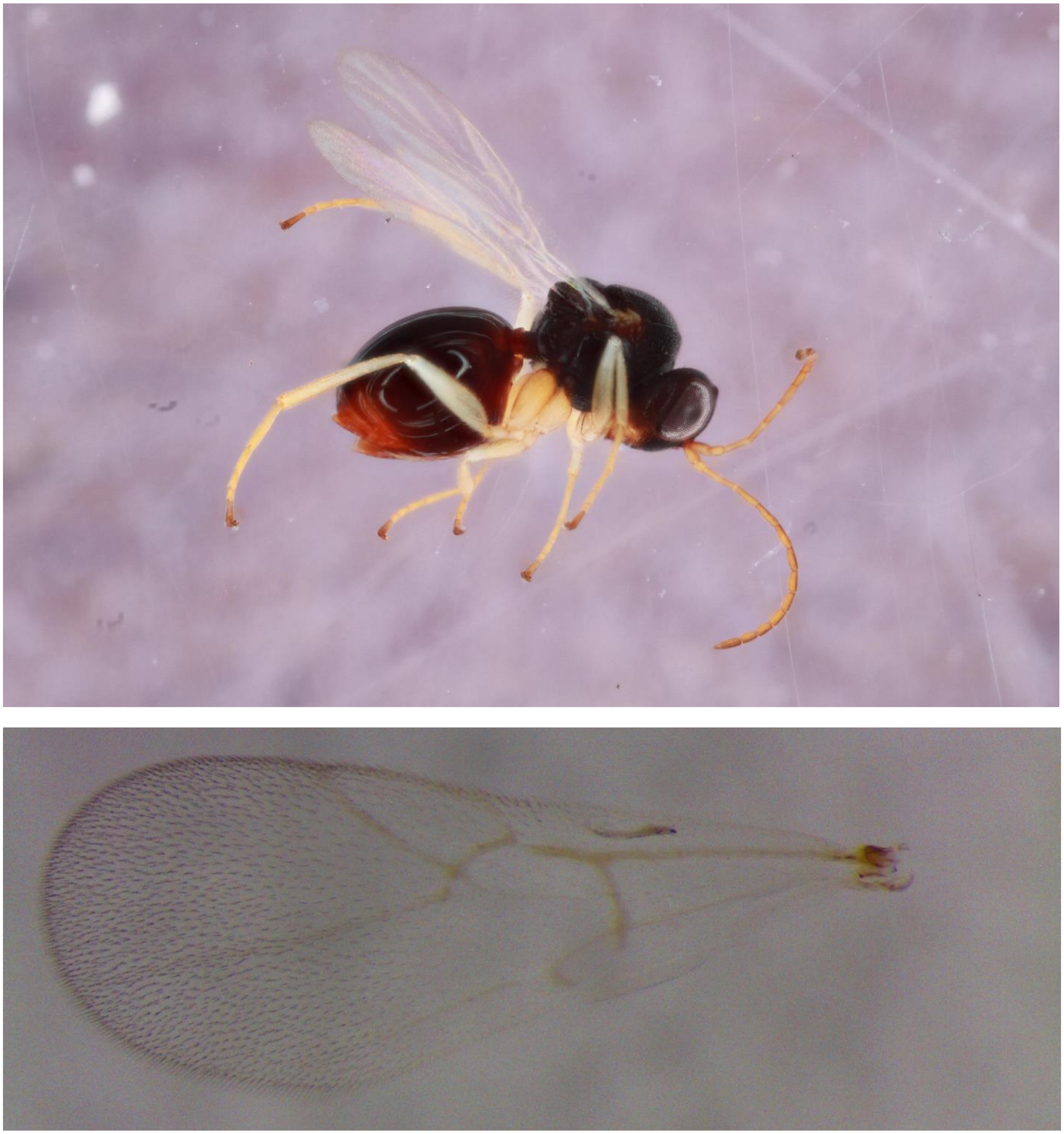

Figure S46. *Synergus campanula* (Clade 27) – 443-003-001 – male
from *Acraspis pezomachoides* on *Quercus alba*, Tiffin, IA

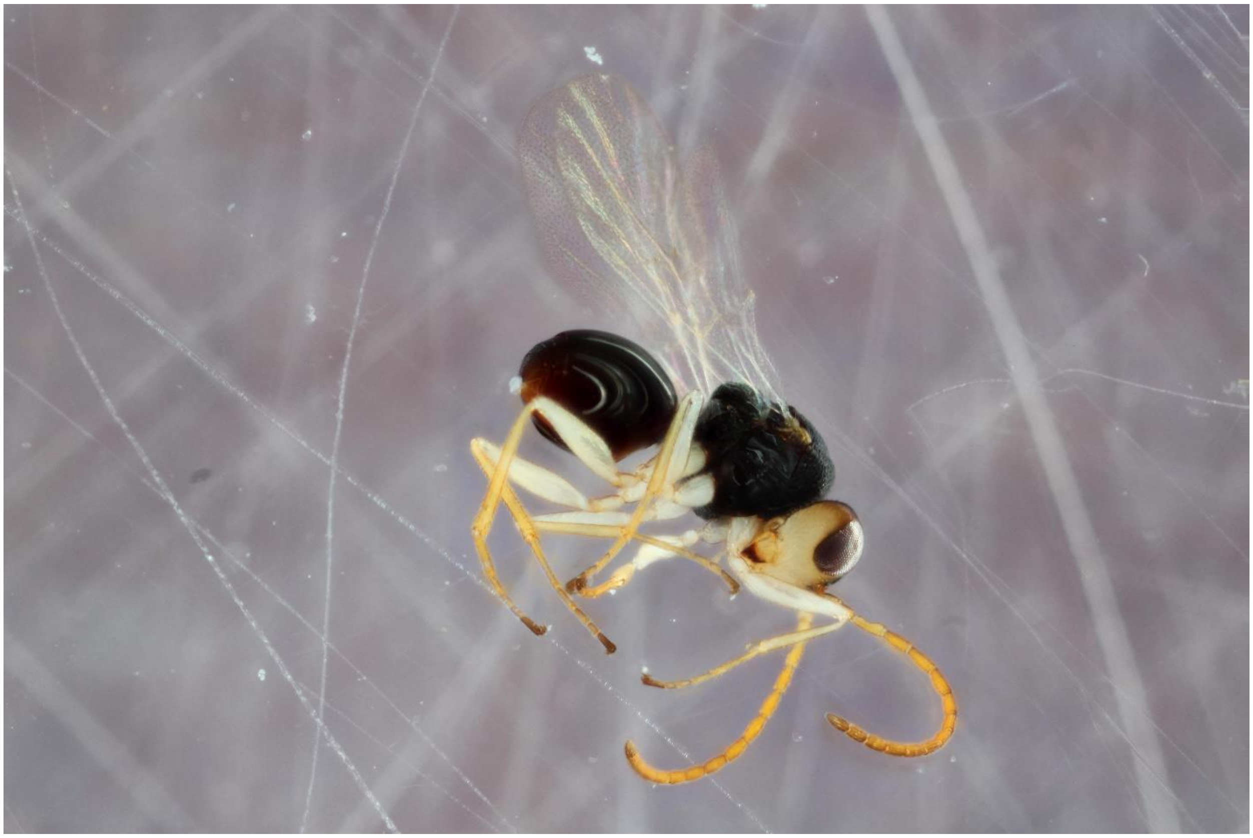

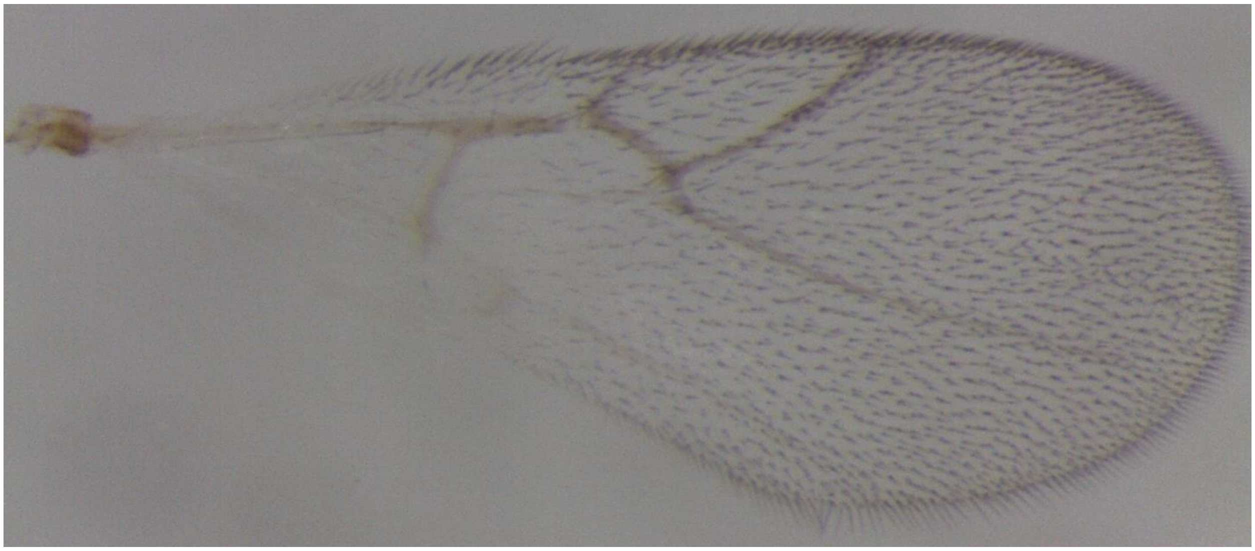
