## Supplemental Figure 48 for "Diversity, host ranges, and potential drivers of speciation among the inquiline enemies of oak gall wasps"

**Figure S48.** ABGD output results based on the alignment of mtCOI sequences for North American *Synergus*. (A) histogram of pairwise distance comparisons of all samples. (B) rank order of pairwise distances. (C) Number of groups determined by ABGD depending on the prior intraspecific divergence cut-off. The yellow boxes represent the initial partition and the red boxes the recursive partition. The liberal (a) and the conservative (b) partitions are highlighted with black boxes.

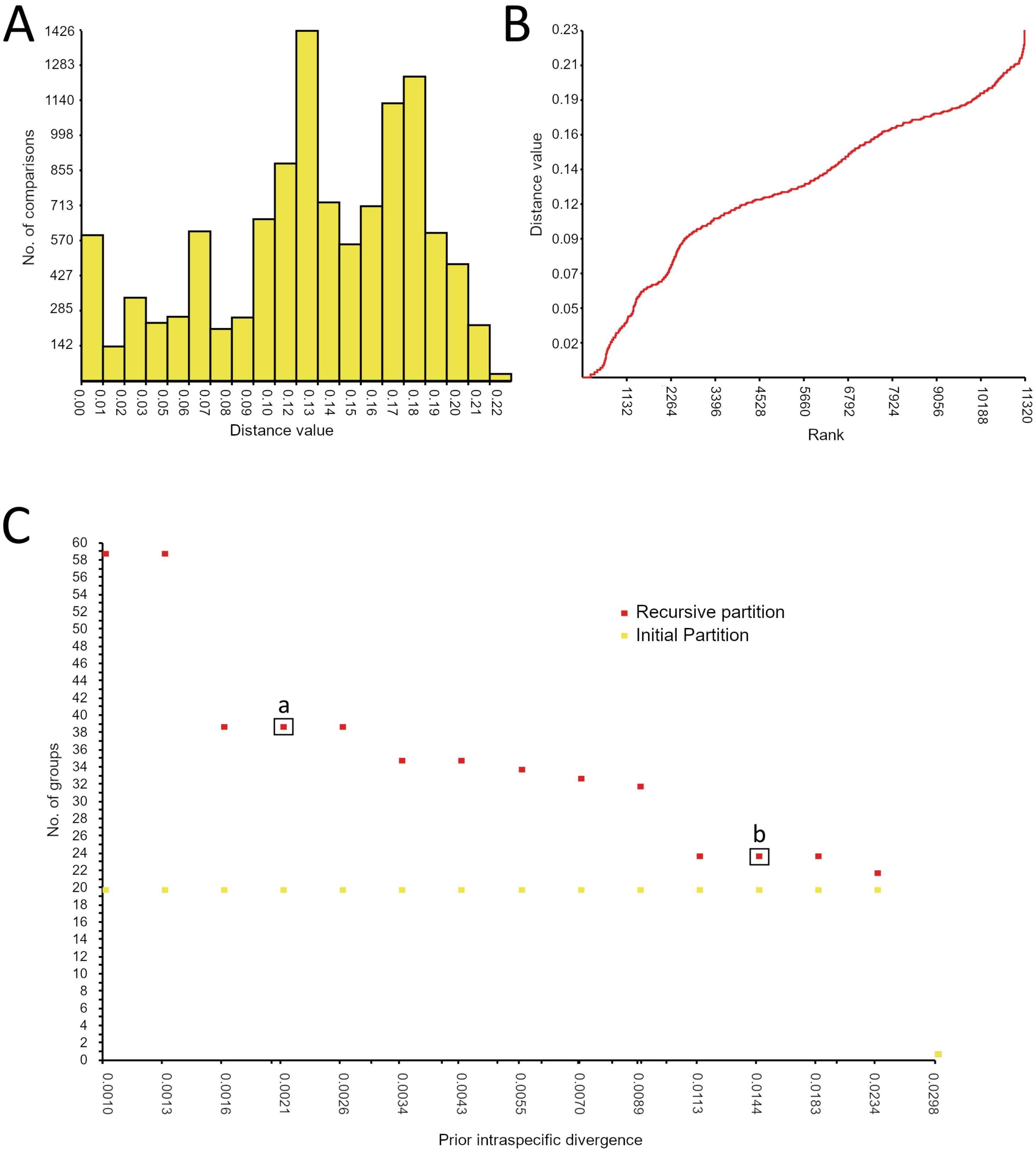
