## Supplemental Figure 49 for "Diversity, host ranges, and potential drivers of speciation among the inquiline enemies of oak gall wasps"

**Figure S49.** Highest supported tree from bPTP model for species delimitation. The values on the branches are Bayesian posterior probabilities supporting whether the clade is a species or not – higher support values indicate higher support for clade being one species. The tips are labeled in the following structure: Lab specific number_Species_Gall host_tree host_Collection location. Ac = *Acraspis*, An = *Andricus*, Am= *Amphibolips*, C = *Callirhytis*, Di = *Disholcaspis*, Dr = *Dryocosmus*, N= *Neuroteras*, Phi = *Philonix*, Phy = *Phylloteras*. For full information about the samples see Supplemental Table 2.

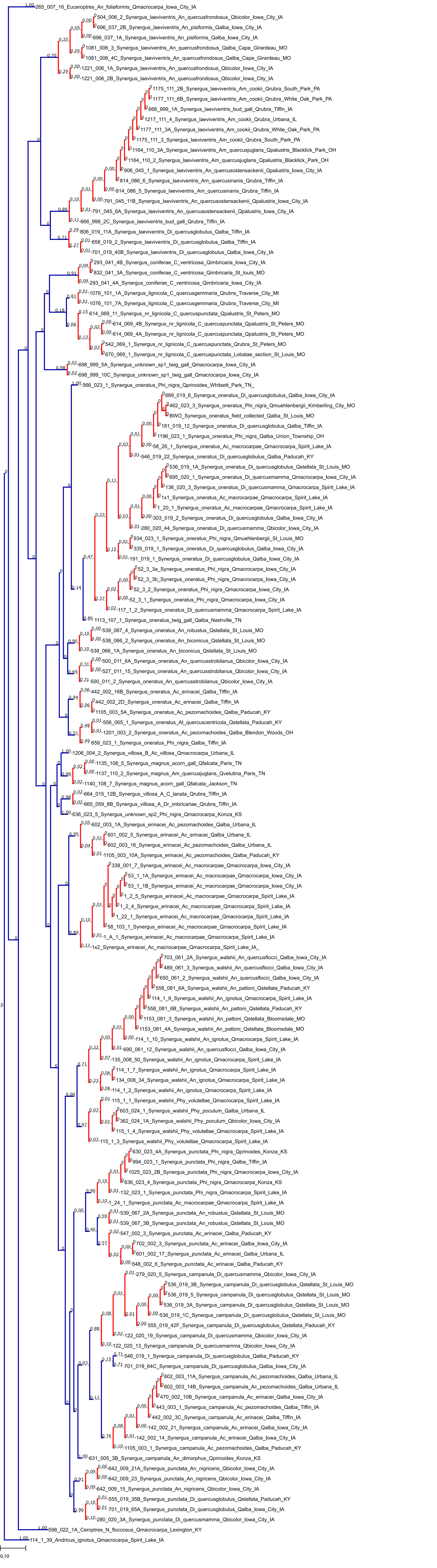
