## Supplemental Figure 50 for "Diversity, host ranges, and potential drivers of speciation among the inquiline enemies of oak gall wasps"

**Figure S50.** Maximum likelihood tree of North American *Synergus* based on mtCOI sequences. The values on the nodes represent bootstrap values. The tips are labeled in the following structure: Lab specific number_Species_Gall host_tree host_Collection location. Ac = *Acraspis*, An = *Andricus*, Am= *Amphibolips*, C = *Callirhytis*, Di = *Disholcaspis*, Dr = *Dryocosmus*, N= *Neuroteras*, Phi = *Philonix*, Phy = *Phylloteras*. For full information about the samples see Supplemental Table 2.

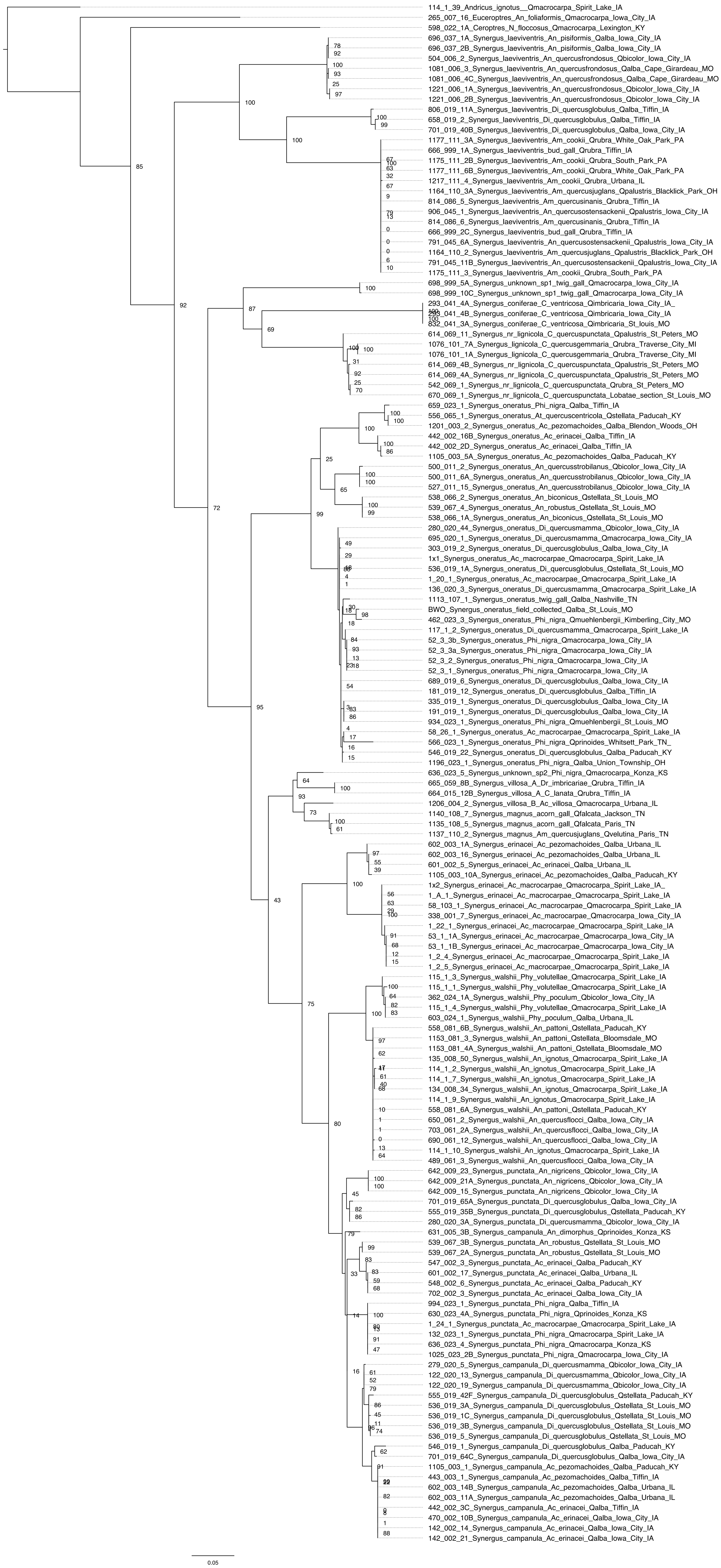
