## Supplemental Figure 51 for "Diversity, host ranges, and potential drivers of speciation among the inquiline enemies of oak gall wasps"

**Figure S51.** Photos of galls collected in this study associated with *Synergus*. The gall photos are grouped based on similar morphology and location found on tree. The numbers following the species name of the gall indicate the *Synergus* clades (see Figure 2) that were reared from that host.

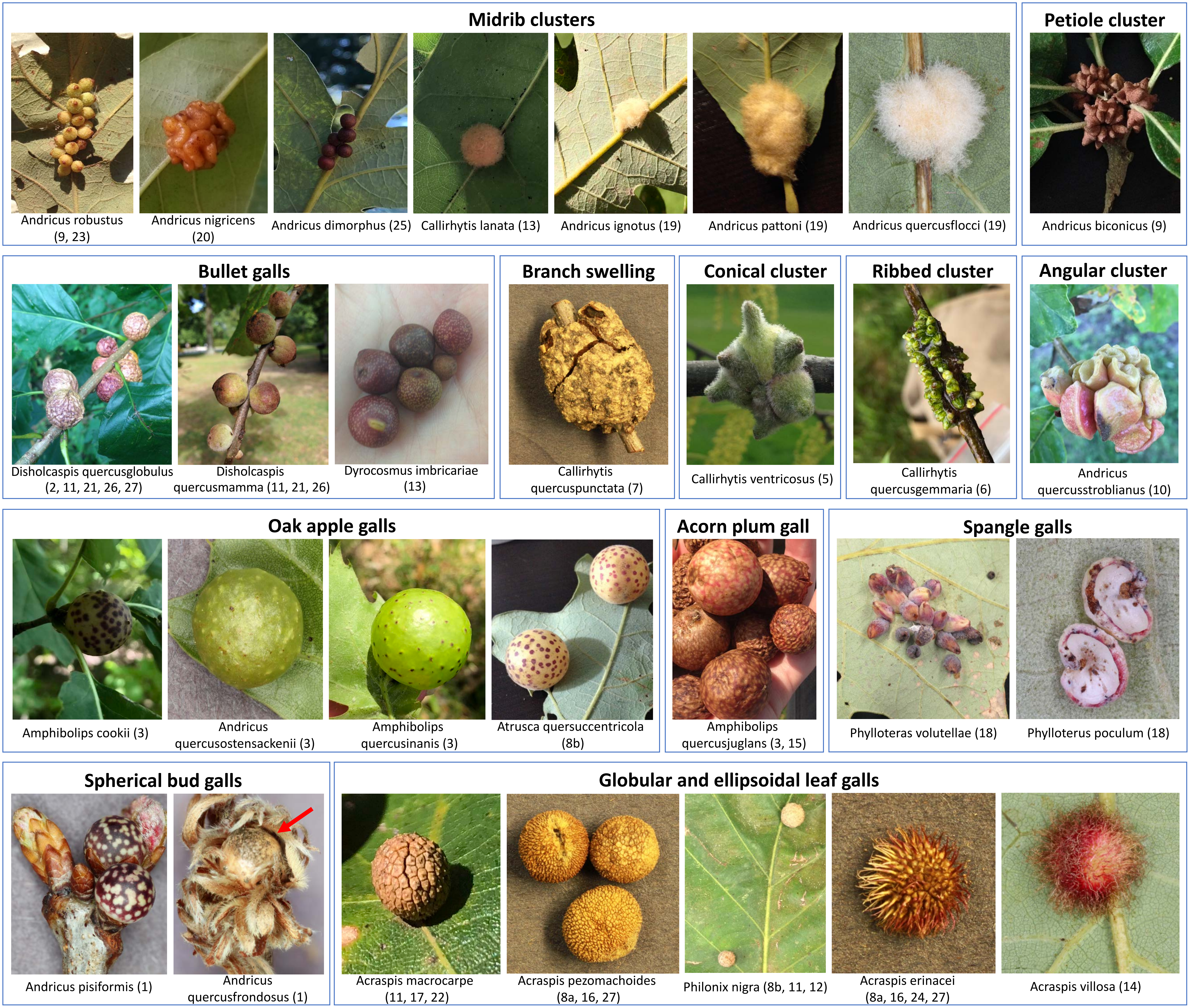
