## Supplemental Figure 52 for "Diversity, host ranges, and potential drivers of speciation among the inquiline enemies of oak gall wasps"

**Figure S52.** Bayesian tree of mtCOI sequences from Nearctic *Synergus* from this study combined with *Synergini* sequences from Acs et al 2010. The gray boxes indicate the Nearctic *Synergus* clades and the green box indicate the Palearctic *Synergus* clade that were collapsed in Figure 6. The values on the tree indicate Bayesian posterior probabilities.

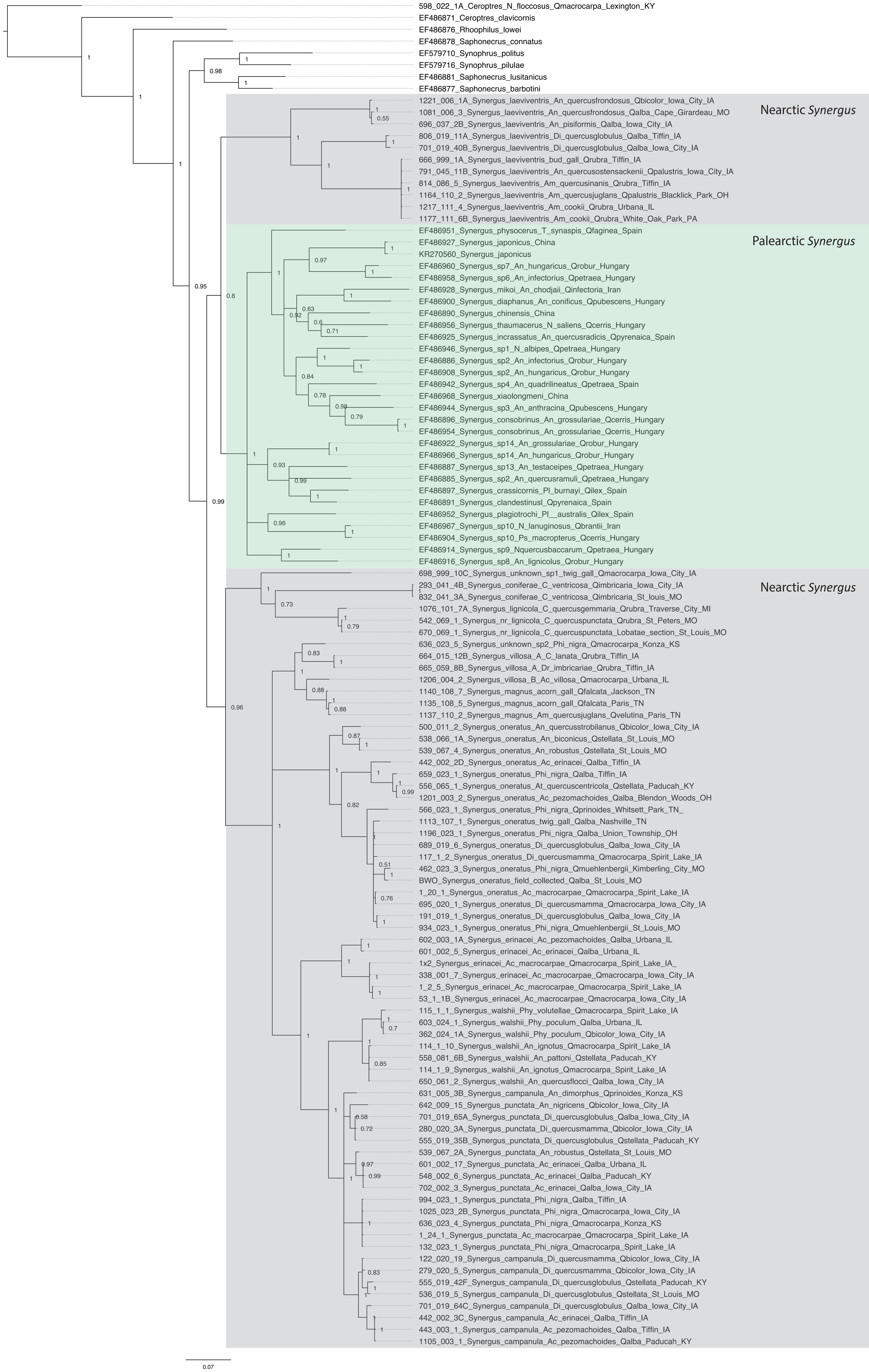
